# The *DGKK* mRNA is a primary target of the Prader-Willi syndrome-associated snoRNA *SNORD116*

**DOI:** 10.64898/2026.10.20.760352

**Authors:** Soline Louis, Oktay Cakil, Audrey Thévenin, Hélène Marty-Capelle, Asmae Maachi, Virginie Marty, Anne Robert, Marie-Eve Chagot, Bruno Charpentier, Xavier Manival, Isabelle Lebars, Hervé Moine, Stéphane Labialle

**Author notes:** Corresponding authors: Stéphane Labialle. These authors contributed equally to this work (Soline Louis, Oktay Cakil). Present address: Cell Biology Division, MRC Laboratory of Molecular Biology, Cambridge CB2 0QH, UK (O. Cakil).

## Abstract

Box C/D small nucleolar RNAs (SNORDs) classically guide site-specific modification of ribosomal RNAs through antisense elements located upstream of their D’ and D boxes, termed ASE1 and ASE2. Yet many orphan SNORDs have no established primary targets, including the paternally expressed multicopy *SNORD116* family implicated in Prader-Willi syndrome. Our previous comparative analysis identified the *DGKK/Dgkk* interaction as the most evolutionarily persistent candidate involving the highly conserved *SNORD116* ASE1 sequence and showed that *SNORD116* depletion increases DGKK RNA abundance in human cells. Here, we re-evaluated this interaction alongside a focused set of recently published candidates and asked whether evolutionary persistence was accompanied by independent evidence of regulation. *DGKK/Dgkk* showed reproducible increases following transient *SNORD116* loss across human iPSC-derived neural cells and in brain tissues from *Snord116*-knockout mice. Its predicted *SNORD116* interaction site combined a marked local depletion of human sequence variation with an overlapping cis-regulatory interval identified in mouse. *SNORD116* deficiency was also associated with increased endogenous DGKκ protein across several mouse brain regions, including the hypothalamus. More broadly, comparison across the candidate set showed that evolutionary persistence, duplex stability and expression response did not systematically align, indicating that sequence-based properties alone do not allow reliable prediction of responses to *SNORD116* loss. Together, these data support *DGKK/Dgkk* as the best-supported primary regulatory target of *SNORD116* to date and establish it as a model for dissecting the regulatory mechanism of this orphan snoRNA family.

## Introduction

Box C/D small nucleolar RNAs (SNORDs) are best known as guides for site-specific 2′-O-methylation. Canonical SNORDs assemble with fibrillarin, NOP56, NOP58 and SNU13/15.5K, and target recognition is mediated by antisense elements located upstream of the D′ and D boxes, termed ASE1 and ASE2, respectively (Cavaillé et al., 1996; Kiss-László et al., 1996; Watkins and Bohnsack, 2012; Henras et al., 2017). Canonical box C/D snoRNA-target recognition relies on a largely continuous guide-target duplex with few mismatches and generally no internal bulges, including a highly constrained 9-bp core located close to the D or D′ box and separated from it by only one or two nucleotides (Chen et al., 2007; Yang et al., 2016; Dupuis-Sandoval et al., 2015). Base pairing positions fibrillarin at a defined nucleotide within the substrate RNA, but sequence-specific recognition by box C/D RNAs is not necessarily coupled to methylation. U3 uses extensive RNA-RNA interactions during pre-rRNA processing, and individual SNORDs have also been implicated in mRNA splicing or 3′-end processing (Kass et al., 1990; Dragon et al., 2002; Falaleeva et al., 2016; Huang et al., 2017). Numerous mammalian SNORDs are classified as “orphans” because no canonical rRNA or snRNA methylation target has been identified for them. Nevertheless, many retain consensus features of box C/D RNAs, raising the possibility that they preserve sequence-specific RNA recognition through their ASE regions. Consistent with this idea, several orphan SNORDs have been proposed to interact with mRNAs or other non-ribosomal RNAs and to influence RNA processing, stability or expression. However, for most orphan SNORDs, the relevant RNA partners, the contribution of individual antisense elements to target recognition, and the functional consequences of these interactions remain poorly defined (Bergeron et al., 2020; Baldini et al., 2021).

The orphan *SNORD116* is of particular interest because loss of its paternal expression contributes substantially to Prader-Willi syndrome (PWS), a neurodevelopmental disorder characterized by neonatal hypotonia and feeding difficulties, impaired postnatal growth, later hyperphagia and obesity, endocrine abnormalities and neurobehavioral alterations (Sahoo et al., 2008; Duker et al., 2010; Tauber and Höybye, 2021). The general organization and imprinting of the PWS-associated region are conserved between human chromosome 15q11-q13 and the corresponding region of mouse chromosome 7. The paternal domain contains several protein-coding genes and non-coding RNAs, including the *SNORD116* and *SNORD115* clusters, which are embedded in repeated introns of the large paternally expressed *SNHG14* transcript. Rare human paternal microdeletions whose common minimal critical region is centered on the *SNORD116* repeat reproduce major PWS features, and mice lacking the paternal *Snord116* cluster show postnatal growth restriction together with metabolic and behavioral abnormalities reminiscent of the disorder (Sahoo et al., 2008; de Smith et al., 2009; Duker et al., 2010; Ding et al., 2008). Despite this strong genetic evidence, the molecular events immediately downstream of SNORD116 loss remain poorly defined.

The human *SNORD116* cluster is composed of 30 paralog genes arranged in tandem, and our previous phylogenetic analyses identified three main sequence subfamilies conserved at least in primates, together with a set of more divergent copies (Baldini et al., 2022; Guibert et al., 2024). The mouse locus has undergone a marked lineage-specific expansion and contained 79 curated paralogs in our previous dataset, but these copies show substantially less sequence diversity than the human repertoire. Molecular and biochemical studies have supported the production of bona fide box C/D snoRNAs from the *SNORD116* copies, and comparative analyses show that the strongest evolutionary constraints within the family affect the basal stem and box elements required for small nucleolar ribonucleoprotein assembly (Bortolin-Cavaillé and Cavaillé, 2012; Baldini et al., 2022). By contrast, the intervening sequences are more variable, except for a major ASE1 sequence that remains deeply conserved across eutherians whereas ASE2 sequences show much greater sequence divergence (Baldini et al., 2022). It suggests that target recognition may not be equivalent across all *SNORD116* copies and that the conserved ASE1 sequence may retain an ancient recognition function. Previous phylogenetic analyses therefore searched for RNA partners whose hybridization potential was maintained together with conserved *SNORD116* antisense sequences across eutherian evolution and identified a restricted group of candidate mRNAs, including *DGKK*, *NHLH2*, *NLGN3* and *RSBN1L* (Kocher et al., 2021; Baldini et al., 2022). Among these, the *DGKK/Dgkk* interaction showed the strongest combination of evolutionary persistence and predicted duplex stability with the conserved ASE1 sequence. Importantly, the evolutionary persistence of the interaction could not be explained solely by independent conservation of the two sequences: in some lineages, synonymous substitutions within *DGKK/Dgkk* were accompanied by compensatory changes in *SNORD116* that preserved base pairing (Baldini et al., 2022). *DGKK* encodes diacylglycerol kinase κ (DGKκ), which phosphorylates diacylglycerol (DAG) to phosphatidic acid (PA) and has been implicated in neuronal signaling and dendritic spine regulation. Its expression is predominantly neuronal and shows a restricted distribution within the brain (Tabet et al., 2016; Habbas et al., 2022). Indeed, in both human and mouse, *DGKK/Dgkk* expression is enriched in the hypothalamus (Sjöstedt et al., 2020), which is of particular interest in the context of PWS that is characterized by hypothalamic dysfunction.

More recent transcriptomic analyses of human PWS tissues and engineered human *SNORD116*-deficient cells have greatly expanded the list of transcripts associated with *SNORD116* loss (Bochukova et al., 2018; Gilmore et al., 2024; Helwak et al., 2024). In parallel, computational searches for RNA complementarity, including snoGloBe-based analyses (Deschamps-Francoeur et al., 2022; Gilmore et al., 2024), and the recently developed snoKARR-seq approach have generated additional candidate interaction partners, including *Snord116* partners in mouse brain (Liu et al., 2025). Most candidates reported so far are mRNAs, and many predicted interactions involve ASE2 sequences, which may reflect genuine biological diversification, but it may also partly result from the greater sequence diversity of ASE2 across the *SNORD116* family. Together, these approaches provide a broad candidate repertoire, but for most proposed targets it remains unclear whether observed expression changes are primary or downstream consequences of *SNORD116* loss and whether the predicted RNA-RNA interaction corresponds to a functionally relevant regulatory site.

Here, we extended our previous phylogenetic analysis and asked whether the *SNORD116/Snord116-DGKK/Dgkk* interaction was supported by independent molecular and functional observations. We re-evaluated the evolutionary persistence of this interaction alongside a focused set of published candidates, examined sequence variation at the corresponding human interaction sites, and compared transcript responses across complementary models of *SNORD116/Snord116* loss. We then tested the predicted *Dgkk* interval by local steric blockade, quantified the DGKκ protein response by targeted proteomics, while assessing the structural context in which ASE1-mediated recognition may occur.

## Results

### Evolutionary analysis distinguishes a persistent *SNORD116-DGKK* interaction

In addition to *DGKK*, we selected candidate *SNORD116*-mRNA interactions reported in previous phylogenetic, transcriptomic or prediction-based studies. We choose candidates with a defined interaction site and additional support from evolutionary analyses, previous studies, or relevance to PWS-related processes. Because the source studies used different algorithms, transcript annotations and definitions of the *SNORD116* recognition sequence, all interactions were reassessed within a common framework. The final comparison comprised 24 interaction sites in 19 reference transcripts (**Suppl. Table 1**). For each human site, we searched the mature SNORD116 repertoire for the most plausible complementary sequence and then followed the corresponding interaction across the eutherian panel used in our previous phylogenetic analysis, except for rhesus macaque, for which the reference was updated to the Indian-origin Mmul_10/rheMac10 assembly (GCF_003339765.1). Core duplexes were defined conservatively from the antisense regions, excluding C/C′/D′/D-box or basal-stem nucleotides (**Suppl. Figure 1**). IntaRNA hybridization energy was used to compare the energetic contribution of base pairing independently of RNA-unfolding penalties (**Suppl. Table 2**). Candidate interactions differed markedly in evolutionary persistence, duplex length and pairing architecture (**Figure 1**). The most persistent ASE1-centered interactions involved *DGKK*, *NLGN3* and *RSBN1L* that were predicted to form uninterrupted duplexes, and *NHLH2* whose interaction remained broadly conserved despite several mismatches and a substantially less canonical pairing architecture. ASE2-centered interactions were more heterogeneous: sites in *BAP1*, *C1QTNF9* and *ANKRD11* persisted across multiple eutherian lineages, whereas others were much more restricted. In particular, *CADPS2* and *DPP10* formed long and energetically favorable human duplexes but showed limited phylogenetic persistence. Several targets, including *ADARB1*, *IRX5*, *SNX33* and *SYK*, contained more than one plausible site, with marked differences in pairing architecture and evolutionary persistence between sites. Overall, hybridization energy, duplex length and pairing architecture were not consistently associated with evolutionary persistence: favorable uninterrupted duplexes occurred among both broadly maintained and lineage-restricted interactions, while the broadly persistent *NHLH2* interaction used a non-canonical pairing architecture.

**Figure 1.**
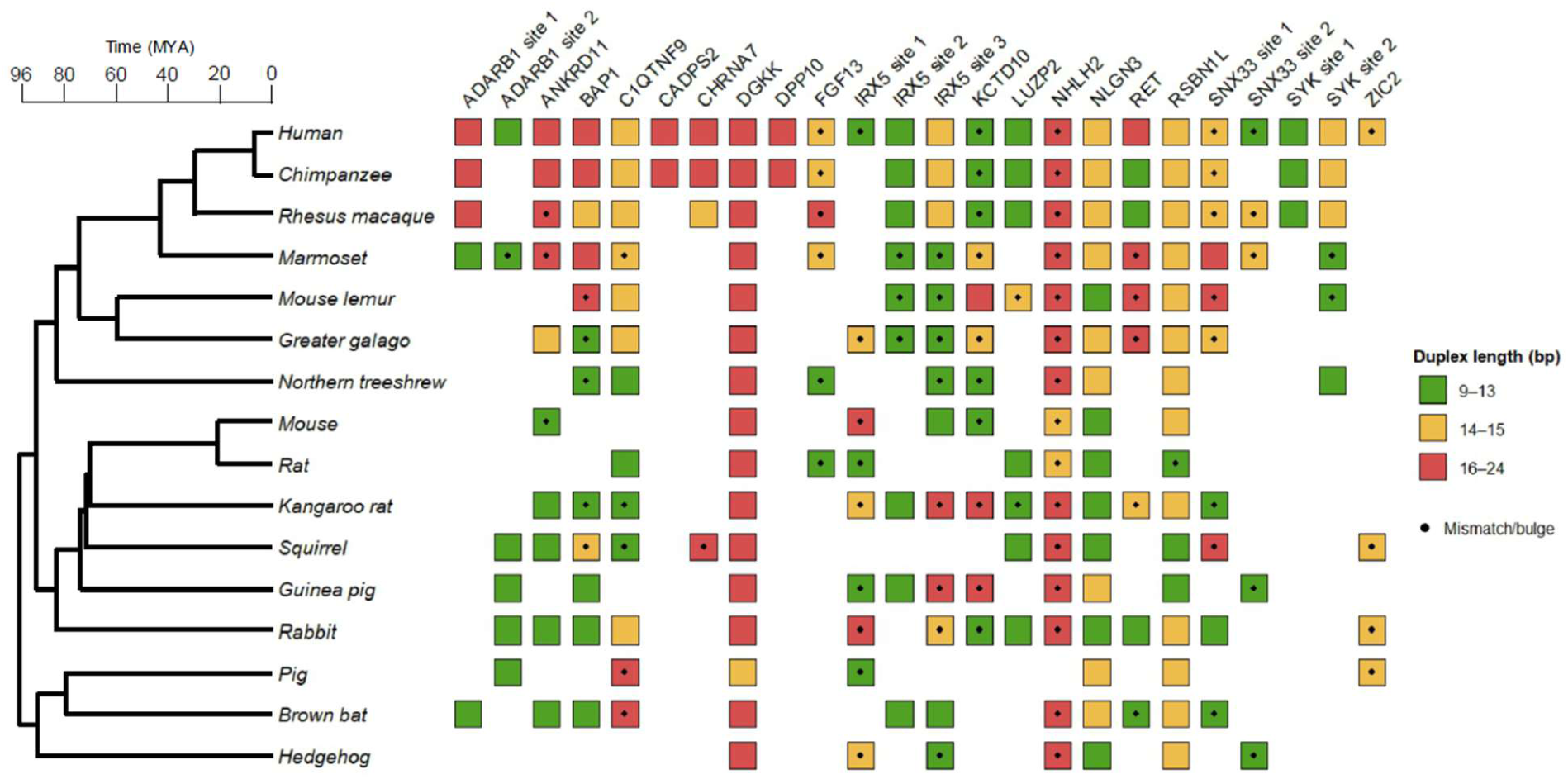
Preservation of candidate *SNORD116*-target RNA duplexes in eutherians. Predicted duplexes were divided into three classes of similar size based on the number of base pairs in each duplex and represented in green, yellow and red. The presence of a mismatch and/or bulge within the duplex is indicated by a dot. The species dendrogram was generated using TimeTree (Kumar et al., 2017). For a detailed presentation of the data, see **Suppl.** Figure 1.

In our selection of candidate target, exonic locations showed greater evolutionary persistence than intronic locations when considering the position of the *SNORD116* interaction site (**Suppl. Table 1**). This difference could not be attributed solely to protein-coding constraint, since highly persistent interactions were also found in untranslated regions, as illustrated by the *NHLH2/Nhlh2* 3’ UTR site. For interactions within coding regions, however, conservation of the target sequence may itself favor persistence of a predicted duplex. Consistent with this, substitutions affecting paired target nucleotides were restricted to third codon positions in several persistent interactions (**Suppl. Table 3**). We therefore searched for a more specific signature of interaction maintenance, namely target-sequence changes accompanied by compensatory changes in *SNORD116* that preserved base pairing. Only *DGKK/Dgkk* provided such a signature: as reported previously, synonymous substitutions within the *DGKK* target site in pig and hedgehog were accompanied by compensatory changes in the corresponding *SNORD116* ASE1 sequence, maintaining the predicted duplex (Baldini et al., 2022). Because these target substitutions are synonymous, preservation of base pairing cannot be explained simply by the conservation of the encoded amino acid sequence. This supports evolutionary maintenance of the *SNORD116-DGKK* interaction beyond constraints imposed by the protein-coding sequence. Its interaction therefore combines broad evolutionary persistence, a continuous ASE1-centered duplex and evidence that pairing capacity has been maintained during target sequence evolution.

### The atypical *SNORD116* C′ sequence impairs canonical C′/D′ folding and SNU13-dependent stabilization

Inspection of the predicted *SNORD116*-target duplexes showed that several interactions could theoretically extend beyond the defined antisense element into neighboring D or D’ box nucleotides (**Figure 2**). For *DGKK/Dgkk*, the conserved interaction is centered on ASE1, although additional pairing toward D’ further stabilizes the predicted duplex. A similar effect was observed for *LUZP2*, whose relatively weak ASE1-centered interaction became substantially more favorable when D’ box proximal nucleotides were included. By contrast, extension of several ASE2-centered interactions toward box D introduced a bulge immediately adjacent to the D box, making these configurations less compatible with canonical box C/D guide geometry. Because D and D’ box nucleotides normally participate in C/D or C’/D’ kink-turn/kink-loop structures, their availability for target pairing is expected to depend on local snoRNA folding. Yet, *SNORD116* is unusual in this respect because its internal C’ motif contains an additional guanosine (UGA<u>G</u>UGA rather than UGAUGA), an insertion conserved across the eutherian orthologs examined. We therefore tested whether this atypical C’/D’ region could form the canonical SNU13-binding K-loop structure expected for a box C/D RNA (van Nues and Watkins, 2017).

**Figure 2.**
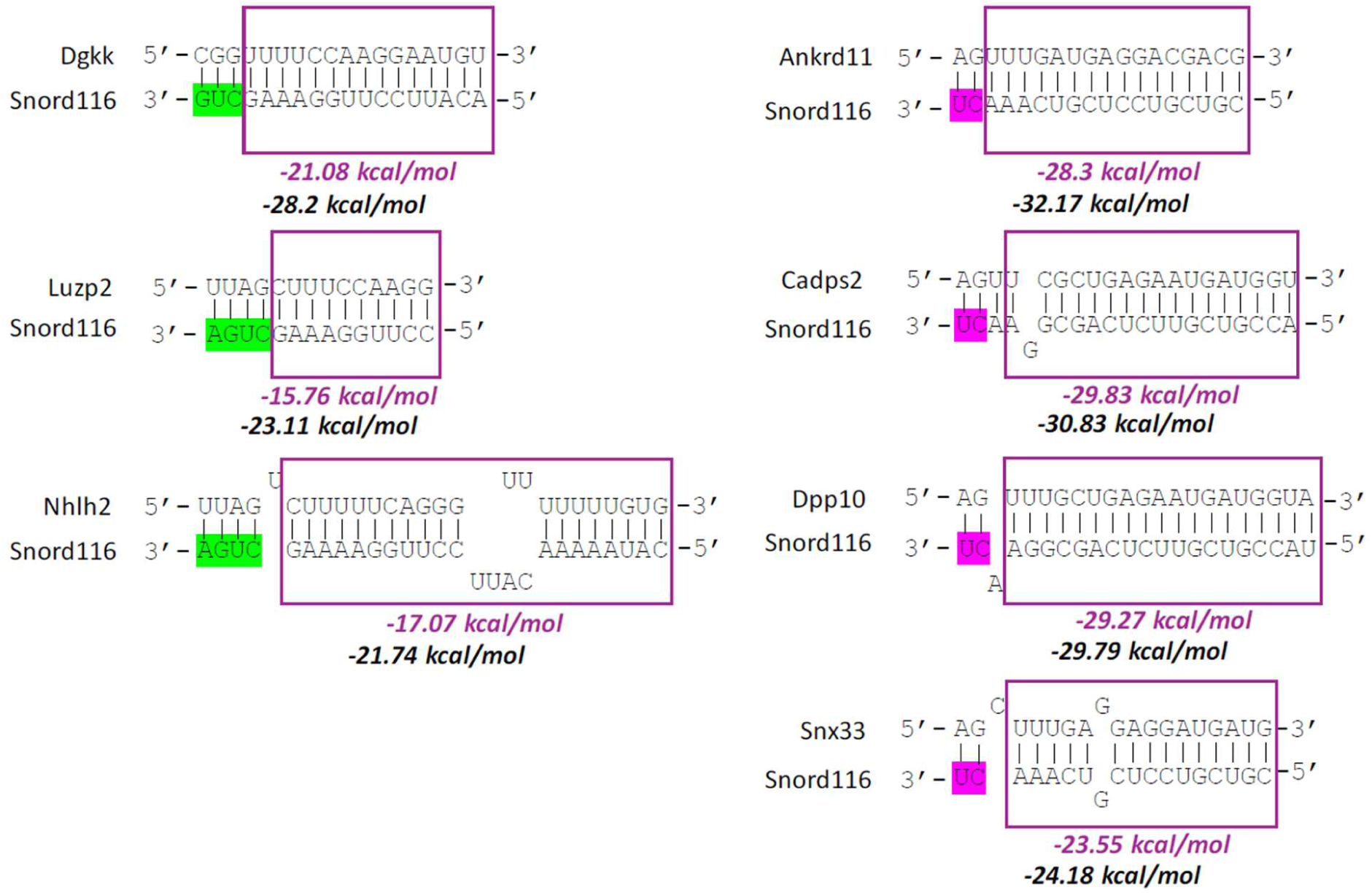
Human *SNORD116*-target RNA duplexes whose base pairing could be extended to D’- or D-box nucleotides. Predicted duplexes are shown with D’- and D-box nucleotides highlighted in green and pink, respectively; predicted hybridization energies before and after extension are indicated in magenta and black, respectively.

Three RNA constructs representing the internal *SNORD116* C’/D’ region were analyzed by Nuclear Magnetic Resonance (NMR). Each contained the D’ box, the atypical C’ motif, adjacent natural nucleotides and two terminal G-C pairs added to stabilize the constructs. One reproduced the highly conserved internal sequence of human *SNORD116-3*, whereas the other two incorporated flanking variants representative of primate subfamilies I and II (SNORD116-3_gI, SNORD116-3_gII), the latter also corresponding to the predominant mouse sequence (**Figure 3A**). In the absence of protein, one-dimensional imino-proton spectra displayed broad and heterogeneous resonances, while 2D NOESY spectra yielded too few correlations for assignment, consistent with a conformationally heterogeneous RNA rather than a single stable K-loop (**Figure 3B**).

**Figure 3.**
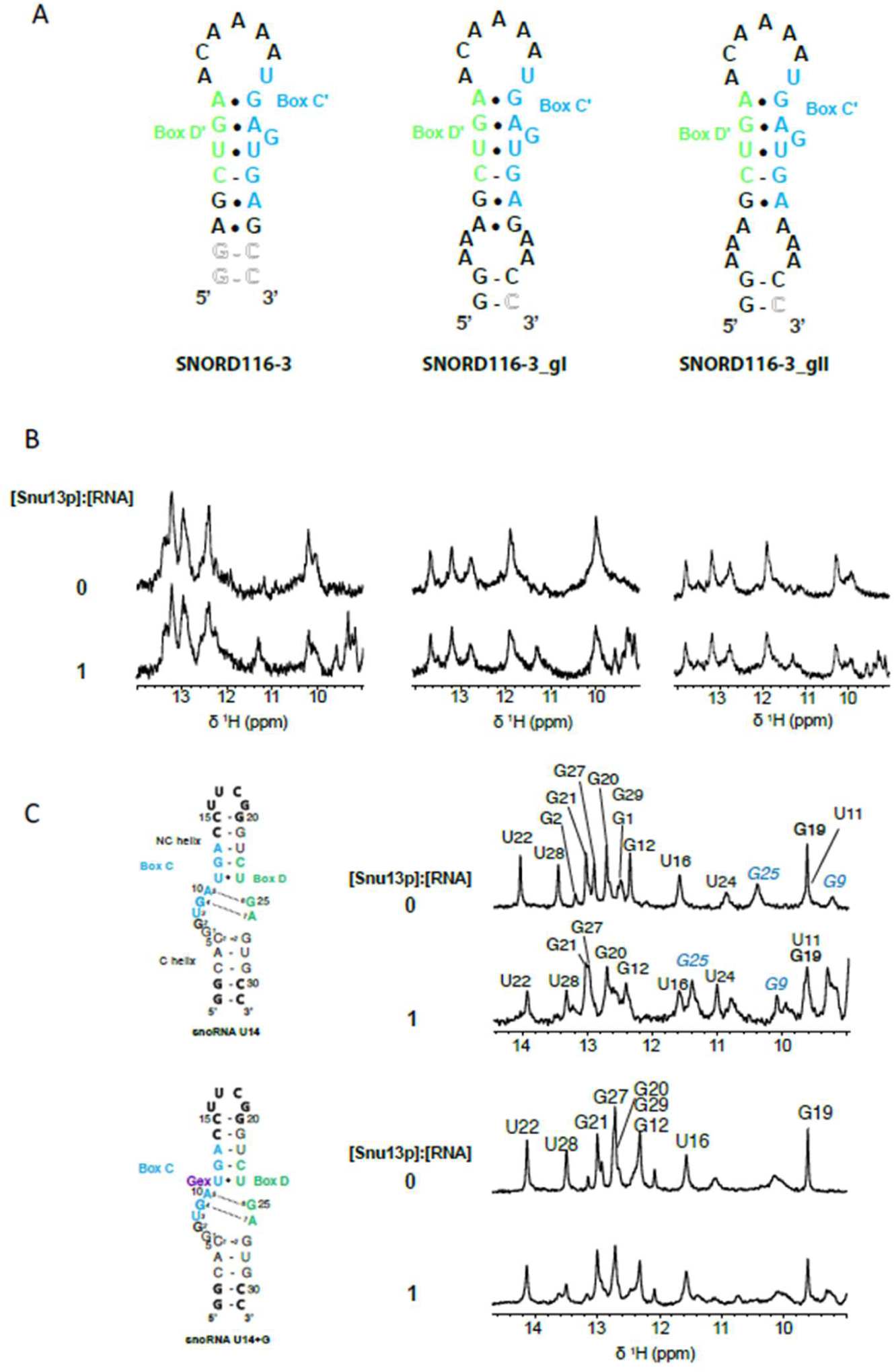
Structural analysis of the *SNORD116* internal C′/D′ region. (A) Proposed K-loop structures of SNORD116-3, SNORD116-3_gI and SNORD116-3_gII, modeled according to the *U18* C’/D’ motif (van Nues and Watkins, 2017). C’- and D’-box sequences are shown in blue and green, respectively; outlined nucleotides correspond to artificial additions. (B) Imino-proton regions of SNORD116-3, SNORD116-3_gI and SNORD116-3_gII in the free state (top) and at a 1:1 Snu13p:RNA ratio (bottom). Spectra were recorded at 15°C in 90:10 H₂O:D₂O; additional signals at the 1:1 ratio arise from the protein. (C) Imino-proton regions of U14 and U14+G in the free state and at a 1:1 Snu13p:RNA ratio, with the corresponding secondary structures shown at left.

We next monitored these RNAs during titration with recombinant yeast Snu13p that shares 69% sequence identity with human SNU13. Because yeast Snu13p has previously been shown to bind the *U14* C/D motif robustly in vitro (Chagot et al., 2019), the *U14*-Snu13p complex was used as a positive control. Whereas Snu13p induced the expected spectral changes in the canonical *U14* C/D module, the three *SNORD116* constructs showed generalized line broadening without the spectral stabilization characteristic of the *U14*-Snu13p complex (**Figure 3B,C**). Introduction of the *SNORD116*-specific extra guanosine into the box C of the *U14* snoRNA (U14+G), whose C/D region forms a K-turn in vitro, destabilized the free K-turn and strongly impaired Snu13p-dependent stabilization (**Figure 3C**). These results indicate that the SNORD116 internal C’/D’ region is poorly competent to form a stable canonical SNU13-stabilized fold, while the U14+G experiment shows that the additional guanosine is sufficient to impair such folding in a canonical C/D context. Although these experiments do not establish the structure of full-length *SNORD116* within its native snoRNA ribonucleoprotein, they suggest that its internal C’/D’ region is less constrained by canonical SNU13-dependent folding than in conventional box C/D RNAs.

### *DGKK/Dgkk* is upregulated across complementary models of *SNORD116/Snord116-*loss

We then compared candidate transcript abundance across complementary models of acute and constitutive *SNORD116/Snord116* loss. Two independent human induced pluripotent stem cells (iPSCs) lines derived from unaffected donors, MCH2-10 (Chamberlain et al., 2010) and YK26 (Zeng et al., 2010), were analyzed following transient *SNORD116* knockdown with a specific Gapmer oligonucleotide, G116, previously used in our work (Baldini et al., 2022). *SNORD116* knockdown was generated at three stages of neuronal differentiation: neural progenitors, neuronal precursors and differentiated neurons. Candidate expression was also examined in *Snord116*-deficient mice using whole brain at postnatal days 3 and 7 and hypothalamus at postnatal day 21 (**Figure 4 and Suppl. Figure 2**).

**Figure 4.**
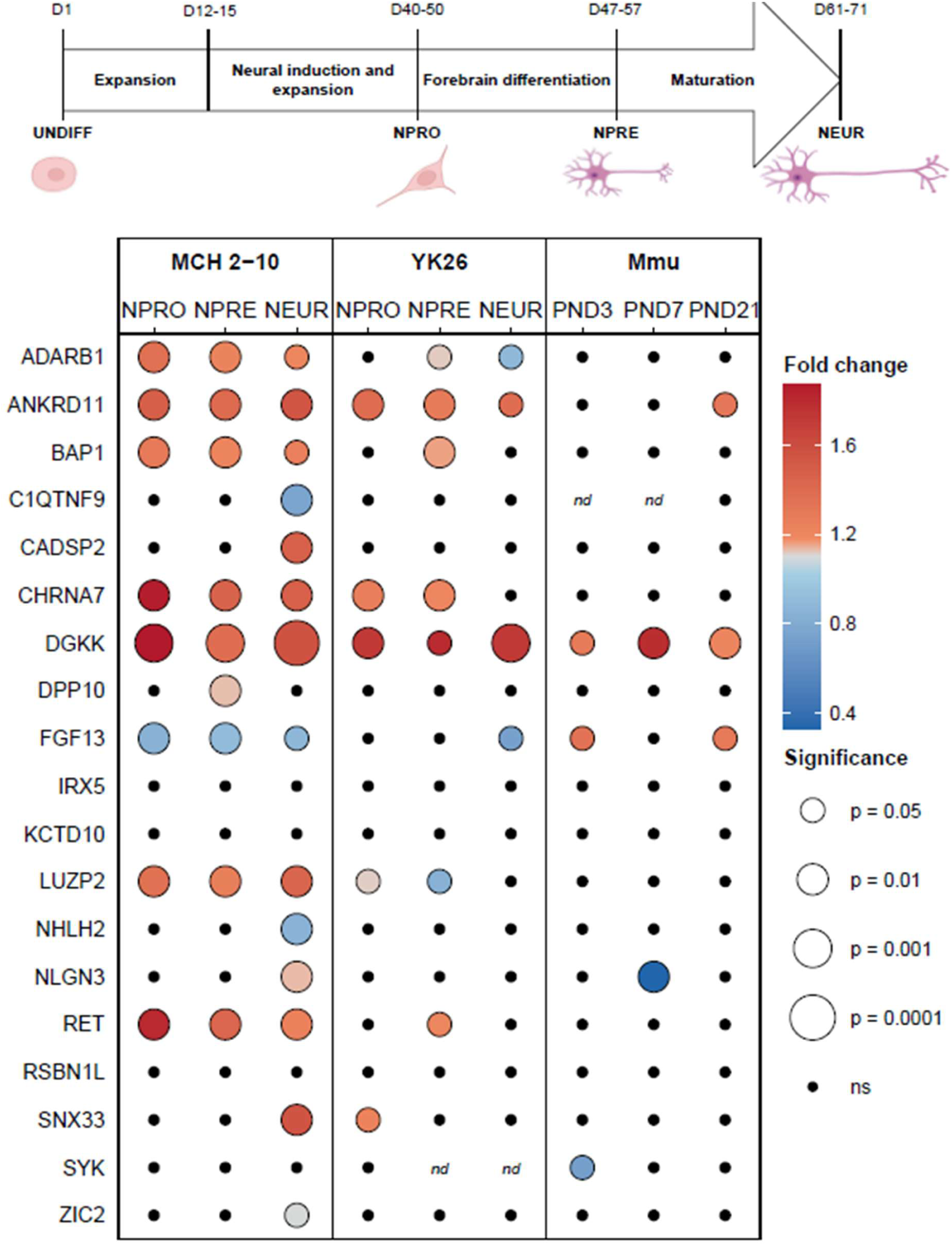
Candidate transcript responses across complementary models of *SNORD116/Snord116* loss. Expression ratios are shown for MCH2-10 and YK26 cells at neural progenitor (NPRO), neuronal precursor (NPRE) and neuronal (NEUR) stages, and for *Snord116*-deficient whole-brain tissues at PND3 and PND7, and hypothalamus at PND21. Ratios correspond to G116/control for human cells and KO/WT for mouse tissues. Color indicates expression fold change and symbol size indicates statistical significance using T-test for YK26 neurons and Mann-Whitney test for all other conditions. Black dots indicate non-significant changes (ns); nd, not detected. For a detailed presentation of the data, see **Suppl.** Figure 2.

*DGKK* showed the most consistent response with significant upregulation in all nine conditions examined. Several other candidates also responded reproducibly, although in fewer conditions. *ANKRD11* was significantly increased in seven of nine conditions, whereas *CHRNA7*, *BAP1* and *RET* were upregulated whenever a significant change was detected. Other transcripts were more context-dependent: *ADARB1* and *LUZP2* were predominantly upregulated but showed one significant change in the opposite direction; *FGF13* decreased in responsive human iPSC conditions but increased in *Snord116*-deficient mouse tissues at PND3 and PND21; and *NLGN3* showed opposite responses in human neurons and mouse brain. *CADPS2*, *DPP10*, *NHLH2*, *C1QTNF9*, *SNX33*, *SYK* and *ZIC2* changed significantly in only one or two conditions, whereas *IRX5*, *KCTD10* and *RSBN1L* remained unchanged.

Because several candidates were originally identified from human transcriptomic studies, whereas their predicted *SNORD116*-mRNA interactions ranged from broadly persistent to largely human-restricted, the mouse experiments provided an opportunity to test whether preservation of a predicted interaction was associated with an expression response to *Snord116* loss. Among the 19 candidates examined, five (*Dgkk*, *Ankrd11*, *Fgf13*, *Nlgn3* and *Syk*) showed significant changes in at least one mouse condition. Of these, only *Dgkk* and *Nlgn3* belonged to the small group of candidates with broadly persistent interaction. Conversely, several interactions with strong evolutionary persistence, including *Nhlh2* and *Rsbn1l*, were not associated with detectable expression changes in the mouse conditions examined. *Fgf13*, by contrast, was significantly upregulated at PND3 and PND21 despite limited persistence of the corresponding predicted interaction. Thus, within our experimental set, persistence of a predicted interaction site in mouse did not consistently predict whether the corresponding transcript responded to *Snord116* loss.

We also compared candidate expression patterns with published transcriptomic studies spanning human PWS tissues, *Snord116*-deficient mice, and human or mouse cellular models of *SNORD116/Snord116* loss (**Suppl. Table 4**). Published datasets showed substantial heterogeneity across experimental systems and, in some cases, between cell backgrounds within the same study. *DGKK/Dgkk* was nevertheless recurrently identified as upregulated across independent *SNORD116/Snord116*-deficient models, including mouse RNA-seq datasets in which it emerged among a relatively limited number of differentially expressed genes (Knott et al., 2022; Marty et al., 2026). Therefore, within our experimental series, *DGKK/Dgkk* was unique in combining broad evolutionary persistence of an uninterrupted ASE1 interaction with significant upregulation whenever a significant change was detected. This combination supported its prioritization for further functional investigation.

### Human sequence constraint and mouse functional assay support a regulatory role for the *DGKK/Dgkk* interaction site

We next asked whether the predicted *SNORD116* interaction sites showed evidence of sequence constraint in human populations. We analyzed exome- and genome-derived single nucleotide variants (SNVs) from gnomAD v4.1.2. Because intronic regions are less well covered in the database, we restricted the analysis to exonic interaction sites. Variants were collapsed to unique SNV-bearing nucleotide positions, and variation within each predicted exonic interaction site was compared with that in the corresponding host exon and complete exonic transcript (**Table 1**). The predicted *DGKK* interaction site showed the strongest local depletion of sequence variation. No SNV-bearing position was identified within the 16-nt ASE1-complementary interval, despite 28 variant-bearing positions in the 103-nt host exon (p = 0.0038). Depletion was also significant relative to the complete exonic transcript (p = 0.0435), and the interaction interval lies at the center of the longest SNV-free stretch identified within the analyzed *DGKK* transcript, which spans 18 nt. Most other candidate sites contained several variant-bearing positions and showed no significant depletion relative to their surrounding exon or transcript. *NHLH2* showed reduced variation relative to its host exon, although it did not reach significance (p = 0.0503), whereas the *ZIC2* site was depleted relative to the complete transcript (p = 0.0357) but not to its host exon (p = 0.1044).

**Table 1:** Sequence variation at predicted *SNORD116*-mRNA interaction sites in human populations. Genetic variation at predicted *SNORD116*-mRNA interaction sites was assessed using SNV data from gnomAD v4.1.2 (GRCh38). All counts refer to distinct nucleotide positions harboring at least one variant. The values were compared with those obtained for the corresponding host exon and the full spliced transcript. Statistical significance was assessed using a one-sided hypergeometric test to determine whether the proportion of positions harboring SNPs was lower within the predicted interaction site than in the corresponding exon or transcript.

| Gene | Site | Transcript | Interaction site (GRCh38) | Hybrid length (nt) | SNV positions in hybrid | Host exon (GRCh38) | Exon length (nt) | SNV positions in exon | p vs exon | Transcript length (nt) | SNV positions in transcript | p vs transcript |
| --- | --- | --- | --- | --- | --- | --- | --- | --- | --- | --- | --- | --- |
| **ADARB1** | Site 1 | ENST00000348831.9 | chr21:45175827-45175842 | 16 | 13 | chr21:45175730-45176664 | 935 | 458 | 0.9985 | 6902 | 1019 | 1 |
|  | Site 2 | ENST00000348831.9 | chr21:45224710-45224722 | 13 | 0 | chr21:45222018-45225627 | 3610 | 196 | 0.4834 | 6902 | 1019 | 0.1251 |
| **ANKRD11** | Site 1 | ENST00000301030.10 | chr16:89285482-89285497 | 16 | 7 | chr16:89279072-89285649 | 6578 | 3941 | 0.1437 | 9301 | 4689 | 0.3888 |
| **BAP1** | Site 1 | ENST00000460680.6 | chr3:52403535-52403550 | 16 | 7 | chr3:52403416-52403894 | 479 | 236 | 0.4236 | 3600 | 1268 | 0.8364 |
| **C1QTNF9** | Site 1 | ENSG00000240654.8 | chr13:24321713-24321727 | 15 | 7 | chr13:24320996-24322531 | 1536 | 267 | 0.9984 | 1849 | 379 | 0.9952 |
| **DGKK** | Site 1 | ENST00000611977.2 | chrX:50401100-50401115 | 16 | 0 | chrX:50401037-50401139 | 103 | 28 | 0.0038 | 7494 | 1332 | 0.0435 |
| **IRX5** | Site 1 | ENST00000394636.9 | chr16:54932778-54932792 | 15 | 8 | chr16:54932498-54932903 | 406 | 191 | 0.7767 | 2398 | 915 | 0.9287 |
|  | Site 2 | ENST00000394636.9 | chr16:54932846-54932856 | 11 | 9 | chr16:54932498-54932903 | 406 | 191 | 0.997 | 2398 | 915 | 0.9995 |
|  | Site 3 | ENST00000394636.9 | chr16:54933466-54933479 | 14 | 8 | chr16:54933077-54934485 | 1409 | 516 | 0.9677 | 2398 | 915 | 0.9573 |
| **KCTD10** | Site 1 | ENST00000228495.11 | chr12:109451669-109451681 | 13 | 8 | chr12:109448655-109451813 | 3159 | 156 | 1 | 3920 | 468 | 1 |
| **NHLH2** | Site 1 | ENST00000320238.3 | chr1:115836504-115836524 | 21 | 4 | chr1:115836377-115838380 | 2004 | 614 | 0.0503 | 2512 | 702 | 0.0895 |
| **NLGN3** | Site 1 | ENST00000358741.4 | chrX:71148855-71148869 | 15 | 2 | chrX:71148846-71148905 | 60 | 12 | 0.3687 | 3975 | 716 | 0.4758 |
| **RSBN1L** | Site 1 | ENST00000334955.13 | chr7:77778627-77778640 | 14 | 4 | chr7:77778530-77783022 | 4493 | 339 | 0.9973 | 6406 | 1287 | 0.8684 |
| **SNX33** | Site 1 | ENST00000308527.6 | chr15:75649427-75649442 | 16 | 13 | chr15:75647912-75650573 | 2662 | 701 | 1 | 8002 | 880 | 1 |
|  | Site 2 | ENST00000308527.6 | chr15:75657020-75657034 | 15 | 12 | chr15:75656962-75662301 | 5340 | 179 | 1 | 8002 | 880 | 1 |
| **ZIC2** | Site 1 | ENST00000376335.8 | chr13:99986692-99986706 | 15 | 3 | chr13:99985323-99986765 | 1443 | 561 | 0.1044 | 2963 | 1361 | 0.0357 |

We further examined the predicted consequences of human sequence variation on *SNORD116*-mRNA hybridization by comparing each reference and alternative target sequence against the complete repertoire of 60 human *SNORD116* ASEs (**Suppl. Table 5**). Across polymorphic sites, individual SNVs frequently weakened the best predicted interaction and altered the repertoire of potential *SNORD116* partners. At every polymorphic site, at least one variant weakened the best predicted interaction, whereas some substitutions strengthened it. Using a threshold of ≤ −20 kcal/mol to define strong predicted interactions, several variants abolished all such interactions, notably at *ADARB1*, *BAP1*, *C1QTNF9*, *IRX5*, *KCTD10*, *RSBN1L* and *SNX33* sites. *NLGN3* was comparatively robust, as none of the four detected variants reduced the number of ASE copies meeting this threshold. These results indicate that naturally occurring substitutions can substantially alter both predicted interaction strength and the repertoire of potential *SNORD116* partners. By contrast, no SNV was detected within the 16-nt *DGKK* interaction interval, which combines broad evolutionary persistence of its ASE1-mediated interaction with marked local depletion of sequence variation in human populations. Notably, the first variant immediately adjacent to the 18-nt *DGKK* SNV-free stretch, chrX:50401118 G>A (rs782276961), falls within the additional D’-dependent extension of the predicted duplex shown in **Figure 2**. On the *DGKK* transcript, this variant corresponds to a C>U substitution and would convert a canonical C-G pair into a terminal U-G wobble, potentially altering duplex stability.

Next, we sought to complement the human population analysis by functionally testing the conserved orthologous *Dgkk* site in mouse cells. We reasoned that, if this site contributes to regulation, local perturbation of the corresponding RNA sequence should affect *Dgkk* abundance. We therefore used the neuroblastoma-derived Neuro2A cells, which express endogenous *Snord116*, to test the regulatory activity of this region. Because basal *Dgkk* expression was low, endogenous *Dgkk* transcription was first increased by CRISPR-activation to provide a robust RNA readout. Then, transient depletion of *Snord116* with the G116 specific Gapmer after *Dgkk* activation reproduced the expected increase in *Dgkk* RNA (**Figure 5A and Suppl. Figure 3**). We then tiled the *Dgkk* transcript with steric-blocking oligonucleotides spanning the conserved *Snord116* interaction site and its immediate flanks (**Figure 5B and Suppl. Figure 3**). The oligonucleotide centered on the predicted interaction interval reduced *Dgkk* RNA abundance, whereas flanking oligonucleotides produced weaker or no effects, suggesting a narrow cis-regulatory region overlapping the predicted *Snord116* interaction site.

**Figure 5.**
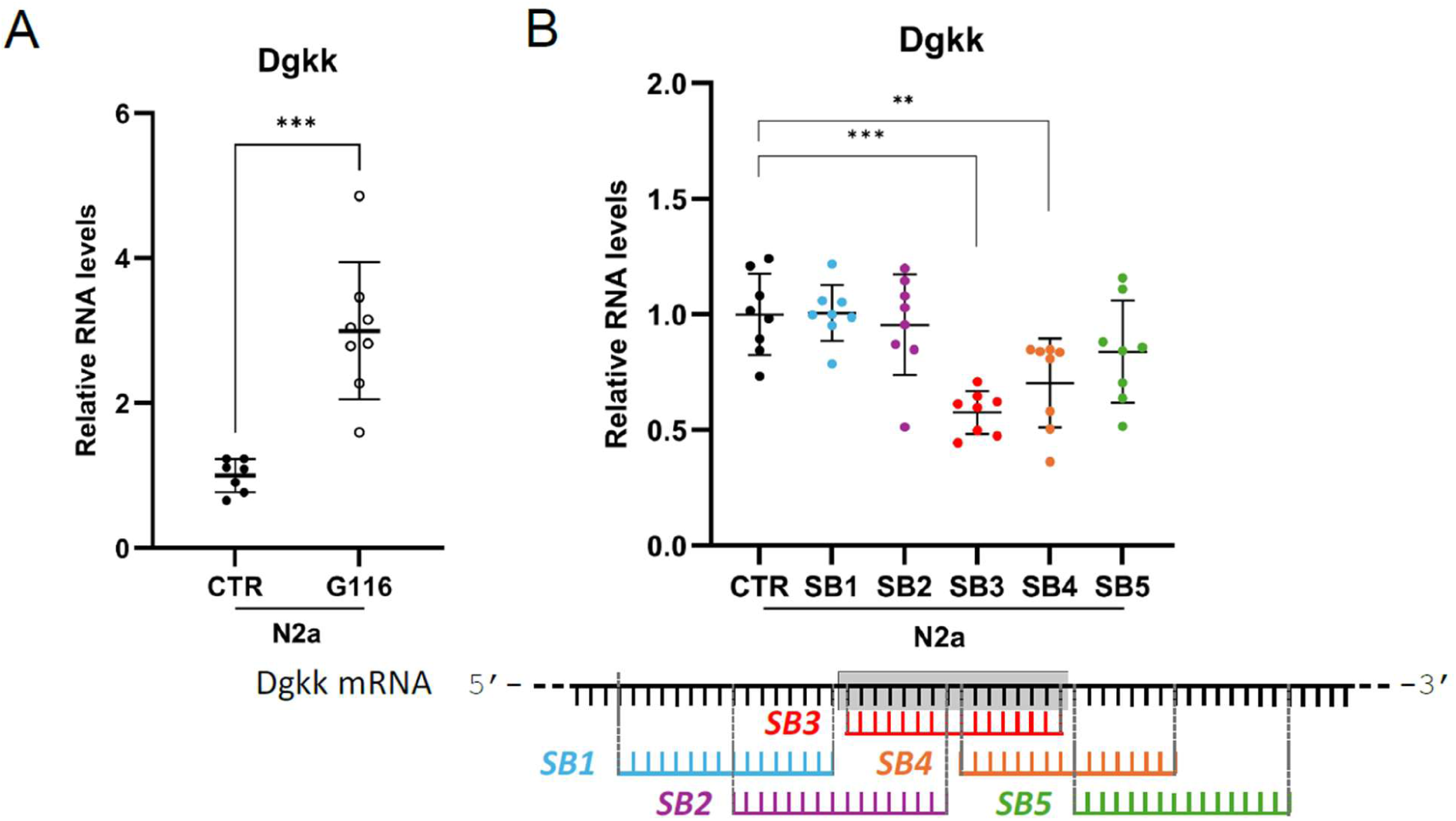
Functional perturbation of the predicted *Snord116-Dgkk* interaction site in Neuro-2a cells. Experiments were performed following CRISPR-mediated activation of endogenous *Dgkk*. (A) Relative *Dgkk* RNA abundance following depletion of endogenous *Snord116* with the G116 Gapmer compared with control-treated cells (n=7-8). Two samples were identified as outliers by the Rout test for *DGKK* in G116 condition and removed; their exclusion did not change the conclusion of the comparison. (B) Relative *Dgkk* RNA abundance following treatment with steric-blocking oligonucleotides SB1-SB5 (n=8). The positions of the five oligonucleotides along the *Dgkk* transcript are shown below the graphs. Data are presented as means ± SD. Mann-Whitney test **P < 0.01, ***P < 0.001.

### *Snord116* loss is associated with increased endogenous DGKκ protein abundance

We next asked whether the *Snord116*-dependent increase in *Dgkk* RNA was accompanied by a change in endogenous DGKκ protein abundance. Because available antibodies lacked sufficient sensitivity or specificity for endogenous DGKκ, we developed a quantitative targeted-proteomics assay based on parallel reaction monitoring with stable isotope-labeled reference peptides (PRM-SIS). The mouse DGKκ signature peptide ISLEFNSR was selected for quantification (**Figure 6A and Suppl. Figure 4**). Assay specificity was established genetically using hypothalamic extracts from *Dgkk*-knockout mice, in which the endogenous peptide signal was absent or reduced to background (**Figure 6B**). Endogenous DGKκ abundance was markedly increased in hypothalamic extracts from *Snord116*-deficient mice relative to wild-type littermates. This increase was observed across the sexes and ages examined (**Figure 6C**). Increased DGKκ abundance was also detected in several additional brain regions (**Figure 6D**). However, the magnitude of the protein increase did not closely parallel *Dgkk* mRNA abundance, which reached significance only in the hypothalamus (**Figure 6E**). This regional specificity is notable given the enrichment of *Dgkk* expression in the hypothalamus and the central contribution of hypothalamic dysfunction to PWS.

**Figure 6.**
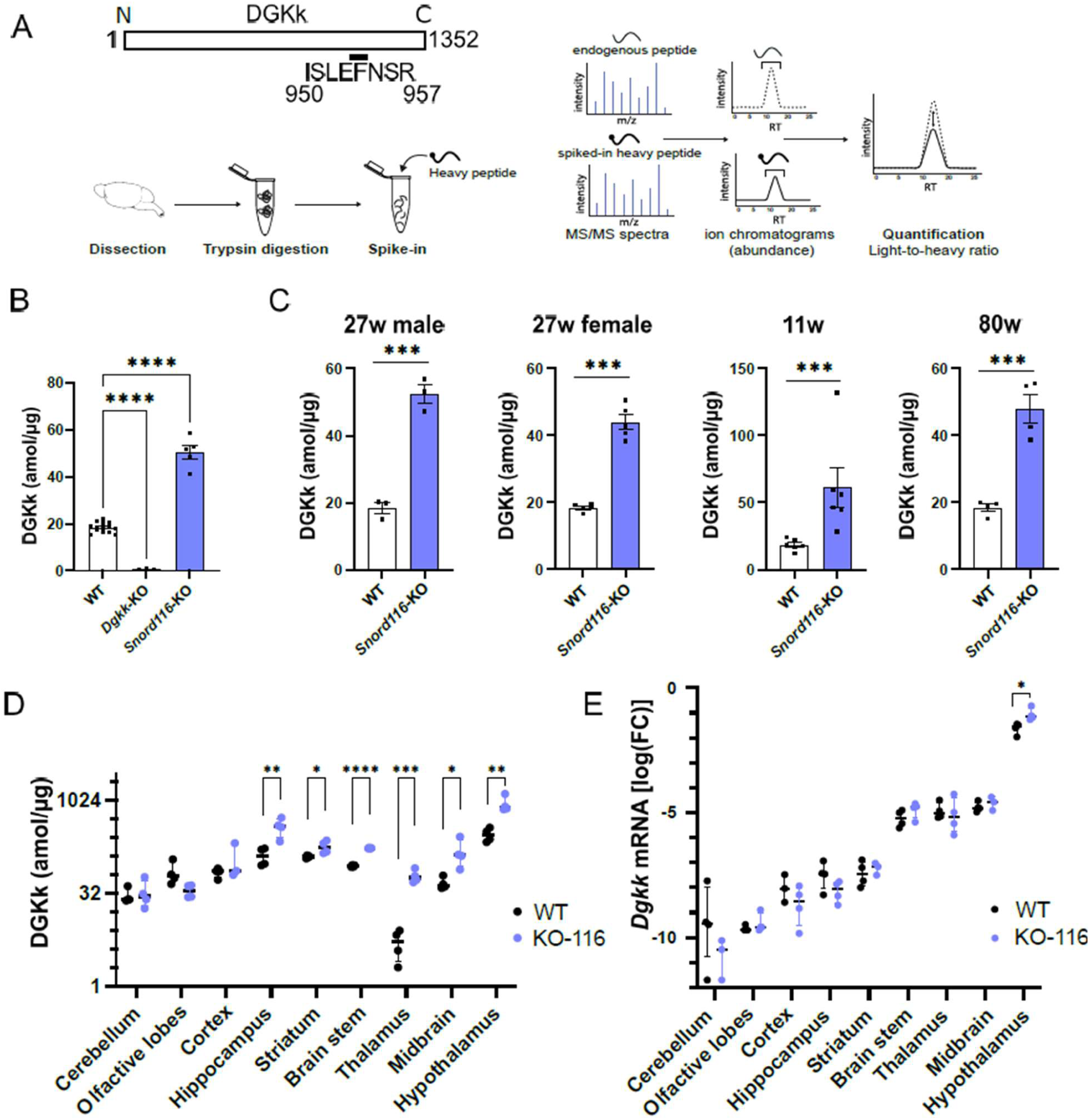
*Snord116* loss is associated with increased endogenous DGKκ protein abundance. (A) PRM-SIS workflow for absolute quantification of endogenous DGKκ using the signature peptide ISLEFNSR and a stable isotope-labeled internal standard. The position of ISLEFNSR within DGKκ and the principle used to distinguish endogenous (light) and isotopically labeled standard (heavy) peptides and derive abundance from the light-to-heavy signal ratio are shown. (B) Absolute DGKκ abundance in hypothalamic extracts from WT, *Dgkk*-KO and *Snord116*-KO mice. Each point represents one mouse. One-way ANOVA with Dunnett’s multiple-comparison test: ****P < 0.0001. (C) DGKκ abundance measured by PRM-SIS in hypothalamic extracts from *Snord116*-KO and WT mice of the indicated age (W, weeks) and sex. Data are means ± SEM; unpaired Student’s t-test, ***P < 0.001. (D) DGKκ abundance across brain regions from *Snord116*-KO and WT mice. Data are means ± SEM; multiple unpaired t-tests with FDR = 1%; *P < 0.05, **P < 0.01, ***P < 0.001, ****P < 0.0001. (E) *Dgkk* mRNA abundance across the same brain regions measured by RT-qPCR. Multiple unpaired t-tests with FDR = 1%; *P < 0.05.

We finally asked whether the increase in DGKκ abundance was accompanied by detectable changes in its lipid substrate or product. Diacylglycerol (DAG) and phosphatidic acid (PA) together with eight additional lipid classes, were quantified in hypothalamic extracts from WT, *Snord116*-KO and *Dgkk*-KO mice using an untargeted mass-spectrometry lipidomic workflow (**Suppl. Figure 5**). No significant genotype-dependent differences were detected in total DAG or PA, or among individual DAG and PA species. Thus, under the conditions examined, the marked increase in DGKκ protein was not accompanied by detectable changes in bulk hypothalamic DAG or PA pools.

## Discussion

Our earlier phylogenetic analysis identified *DGKK* as a strong *SNORD116* candidate target because its predicted interaction with the conserved *SNORD116* ASE1 sequence persisted across eutherian evolution (Baldini et al., 2022). The present study provides several additional and complementary lines of support for this interaction. First, loss of *SNORD116/Snord116* consistently increased *DGKK/Dgkk* RNA abundance across the human and mouse models examined here. Independent transcriptomic studies have also repeatedly identified *Dgkk* as differentially expressed in *Snord116*-deficient mouse tissues, often among a relatively small number of differentially expressed genes (Knott et al., 2022; Marty et al., 2026). If not demonstrative, this recurrence across acute and constitutive loss-of-function models, developmental contexts and species is compatible with *DGKK/Dgkk* being close to the primary molecular consequences of *SNORD116* loss. Together with the human population data, the site-directed perturbation experiments and the increase in endogenous DGKκ protein, these findings support *DGKK/Dgkk* as the best-supported primary regulatory target of *SNORD116/Snord116* identified to date.

Comparison of the broader candidate set also showed that no single sequence-based feature examined was sufficient to predict a reproducible response to *SNORD116* loss. A favorable hybridization energy, a long uninterrupted duplex and broad evolutionary persistence did not consistently predict an expression change in candidate abundance. Nor did an obvious combination of these features emerge as a reliable predictor: some broadly persistent interactions, including those involving *NHLH2* and *RSBN1L*, showed little or no response under the conditions examined, whereas other candidates responded despite more limited evolutionary persistence of the corresponding interaction, as illustrated by *FGF13*. These observations argue against ranking candidate targets from hybridization strength or evolutionary persistence alone, or from a simple additive combination of such properties. Rather, they suggest that the candidate set, assembled largely from expression changes observed in a limited number of models and from sequence-based predictions, is biologically heterogeneous. It may include direct *SNORD116* targets acting across multiple contexts, direct targets with context-dependent effects, transcripts indirectly affected downstream of *SNORD116* loss, and interactions with no detectable regulatory consequence under the conditions examined. Accordingly, sequence-based properties alone are insufficient to distinguish among these possibilities. Conversely, the absence of a detectable change in steady-state RNA abundance does not exclude direct *SNORD116* targeting, as binding could instead affect other RNA parameters such as splicing or stability, or could occur without any regulatory consequence. As an example, although we have previously proposed that *SNORD116* favors *NLGN3* exon 3 inclusion, we did not detect transcripts lacking exon 3 in the human and mouse samples examined here (data not shown), preventing us from assessing any further increase in exon inclusion.

Among the candidates examined, *DGKK* also showed the clearest evolutionary maintenance of predicted base pairing with *SNORD116* ASE1. In pig and hedgehog, synonymous substitutions within the *DGKK/Dgkk* target sequence are accompanied by changes in *SNORD116* ASE1 that preserve the predicted pairing. Because these substitutions do not alter the DGKκ protein sequence, this pattern is difficult to explain by protein-coding constraint alone and is consistent with compensatory evolution maintaining the interaction. The human population data provide complementary evidence at a much shorter evolutionary timescale. The predicted *DGKK* interaction site is unusually depleted of sequence variation and lies within the longest local SNV-free interval of the transcript. This observation does not establish that selection acts specifically to preserve *SNORD116* binding, since the sequence may also be constrained by other features of the transcript, including coding requirements, RNA structure or recognition by other factors. It nevertheless suggests that the predicted interaction overlaps an unusually constrained local RNA element in human populations. Complementary functional evidence on the target side was obtained by steric blocking of the orthologous *Dgkk* sequence in mouse cells. Steric block oligonucleotides centered on the predicted *Snord116* interaction site decreased *Dgkk* level, whereas oligonucleotides targeting adjacent sequences had weaker or no effects. The effect was opposite to that produced by *Snord116* depletion, suggesting that the accessibility of this RNA interval contributes to the regulation of transcript abundance. Occupancy of the region could interfere with *Snord116* binding, with another RNA-RNA or RNA-protein interaction, or with local RNA structure. Together, these experiments suggest the presence of a functional cis-regulatory element overlapping the predicted *SNORD116/Snord116* site. The main unresolved question is whether snoRNA directly binds the mRNA at this site in cells; direct interaction mapping, or synonymous editing of the *DGKK/Dgkk* interaction site to disrupt *SNORD116/Snord116* complementarity, would provide direct tests of the predicted interaction. Yet, direct demonstration of an endogenous *SNORD116*-*DGKK* detection may be challenging, e.g., given the generally low abundance of *DGKK* transcripts across tissues and cell types.

The structural data provide additional clues about how *SNORD116* may recognize RNA targets. In the isolated *SNORD116* C’/D’ module, the RNA failed to adopt a stable canonical kink-loop and was not efficiently stabilized by Snu13p. Introducing the extra guanosine characteristic of the *SNORD116* C’ box into the canonical *U14* C box produced a similar defect, indicating that this nucleotide is sufficient to disrupt canonical folding in this experimental context. Although these experiments do not establish the structure of full-length *SNORD116* in cells, they do suggest that the internal C’/D’ region is less constrained in a canonical SNU13-bound fold and may leave ASE1-adjacent nucleotides more accessible for alternative RNA interactions. Still, whether D’-proximal nucleotides contribute to target recognition remains unresolved and will require dedicated structural and functional approaches. This question also intersects with another unresolved issue: whether mRNA targets, particularly DGKK/Dgkk, undergo fibrillarin-dependent 2’-O-methylation guided by the C’/D’ module. Addressing this question will likely require transcript-specific approaches, such as Nm-VAQ (Tang et al., 2024), among other methods.

The increase in endogenous DGKκ protein extends the effect of *Snord116* loss beyond the changes detected at the RNA level. DGKκ abundance was increased in several brain regions and across the sexes and ages examined, whereas changes in *Dgkk* RNA were limited and reached significance only in the hypothalamus. This discrepancy suggests that steady-state RNA abundance alone may not account for the magnitude of the protein increase. Changes in translation, RNA processing or localization, or protein turnover, could also contribute. Notably, the hypothalamus was the only region showing significant changes at both the *Dgkk* RNA and DGKκ protein levels, a finding of particular relevance given the enrichment of *Dgkk* expression in this region and the central role of hypothalamic dysfunction in PWS. Despite the increase in DGKκ protein, we detected no significant change in total hypothalamic DAG or PA, or among individual DAG and PA species. This lack of a bulk lipid phenotype does not necessarily imply that increased DGKκ abundance is functionally silent. DAG and PA pools are controlled by multiple enzymes, including ten mammalian DGK isoforms, and DGKκ-dependent effects may be restricted to particular cell types, subcellular compartments or transient signaling events that are not resolved by whole-tissue steady-state lipid measurements. Future studies combining cell type-resolved lipid measurements with direct readouts of DGKκ-dependent signaling should determine whether the increase in DGKκ protein produces localized functional effects that are masked in whole-tissue analyses. An important next step will be to determine whether and to what extent *DGKK/Dgkk* dysregulation contributes to PWS-related phenotypes, for example by selective normalization of DGKκ abundance or activity in human PWS-derived iPSCs and *Snord116*-deficient mice, followed by assessment of neuronal function in cell models, of growth, metabolic and behavioral phenotypes in vivo, and transcriptomic, proteomic and lipidomic profiles in both systems. In parallel, synonymous editing of the *Dgkk* interaction site to disrupt *SNORD116* complementarity without altering the DGKκ protein sequence would directly test the functional requirement for the predicted interaction.

More broadly, the present study illustrates that assigning primary targets to orphan snoRNAs requires convergent evidence from complementary approaches, with no single criterion sufficient on its own. In this context, the present findings support *DGKK/Dgkk* as the best-supported primary regulatory target of *SNORD116/Snord116* identified to date, to our knowledge. *DGKK/Dgkk* therefore provides a useful model to investigate the molecular basis of *SNORD116*-dependent mRNA recognition and regulation, a key step toward understanding how this orphan SNORD family exerts its biological functions.

## Materials and Methods

### Evolutionary persistence and RNA-RNA interaction analysis

Candidate interaction sites were taken from the literature or identified by re-analysis of the corresponding human transcript with full-length *SNORD116* sequences using IntaRNA. *SNORD116* orthologous sequences were taken from the curated dataset described in Baldini et al. (2022). Orthologous target regions were retrieved from the UCSC Genome Browser by BLAT using the human target interval and flanking sequence, followed by manual verification of local synteny and transcript orientation. Hybridization Energy values reported by IntaRNA were used to compare base-pairing strength independently of the energetic cost of unfolding RNAs. IntaRNA was run with default parameters for all candidates except *IRX5*, for which the previously proposed interaction was not recovered under the default settings and required relaxed interaction parameters.

### Human population variation at candidate *SNORD116* interaction sites

Genetic variation was assessed using gnomAD v4.1.2 exome and genome datasets mapped to GRCh38. Single-nucleotide variants passing quality filters in at least one dataset were retained; variants present in both exomes and genomes were counted once, multiallelic variants at the same nucleotide were collapsed to a single variant-bearing position, and indels or other non-SNV variants were excluded. Analyses therefore used the number of distinct nucleotide positions carrying at least one PASS SNV, irrespective of allele count or frequency. For each interaction site, local variation was compared with that observed in the host exon and across the complete exonic sequence of the selected reference transcript. Transcript exonic length was reconstructed from HGVS cDNA coordinates. Depletion of sequence variation was tested using a one-sided hypergeometric test.

### Differentiation and oligonucleotide treatment in human induced pluripotent stem cells

Human induced pluripotent stem cells (iPSCs) were maintained in 6-well plates coated with 10 µg/mL Vitronectin XF™ (StemCell Technologies) in mTeSR™ Plus medium (StemCell Technologies). Medium was replaced daily, and cells were passaged every 3-4 days at 80-90% confluence using ReleSR™ (StemCell Technologies), at split ratios ranging from 1:3 to 1:8. The ROCK inhibitor Y-27632 (StemCell Technologies) was added at seeding to improve cell survival. Cells were maintained at 37°C in a humidified atmosphere containing 5% CO₂. For neural induction, iPSCs were differentiated into neural progenitor cells (NPROs) using the STEMdiff™ SMADi Neural Induction Kit (StemCell Technologies) according to the manufacturer’s monolayer protocol. Cells were dissociated using Gentle Cell Dissociation Reagent (StemCell Technologies), and 2 × 10⁶ single cells were seeded per well onto tissue culture-treated 6-well plates (StemCell Technologies, 38016) coated with 15 µg/mL poly-L-ornithine (PLO; Sigma-Aldrich) and 10 µg/mL laminin (Sigma-Aldrich). Cells were cultured in STEMdiff™ Neural Induction Medium supplemented with SMADi and Y-27632, with complete medium replacement daily for 7 days. Early NPROs were then dissociated using Accutase™ (StemCell Technologies) and replated onto PLO/laminin-coated 6-well plates at 2.5 × 10⁵ cells/cm² in STEMdiff™ Neural Induction Medium supplemented with SMADi and Y-27632. Medium was changed daily for a further 7 days. After two passages in STEMdiff™ Neural Induction Medium, NPROs were maintained in STEMdiff™ Neural Progenitor Medium, seeded at 2.5 × 10⁵ cells/cm² for routine maintenance and at 1.5 × 10⁵ cells/cm² for experimental use. To generate forebrain neuronal precursor cells (NPREs), NPROs were dissociated with Accutase™ after one passage and seeded onto PLO/laminin-coated 6-well plates in STEMdiff™ Forebrain Neuron Differentiation Medium (StemCell Technologies), at 2.5 × 10⁵ cells/cm² for routine differentiation or 1.5 × 10⁵ cells/cm² for experiments. Medium was replaced daily for 7 days, until cultures reached 80-90% confluence. For neuronal maturation, NPREs were dissociated using Accutase™ and plated in 6-well plates coated with 15 µg/mL PLO and 5 µg/mL laminin at 5.5 × 10⁵ cells per well in STEMdiff™ Forebrain Neuron Maturation Medium (StemCell Technologies). Cells were maintained in maturation medium for at least 14 days to obtain differentiated neurons. Expression of stage-specific differentiation markers was monitored throughout the differentiation procedure (**Suppl. Figure 6**). For oligonucleotide treatment, cells were transfected using Lipofectamine 3000 (Thermo Fisher Scientific, L3000008). The G116 Gapmer oligonucleotide targeting human SNORD116 (TCACTCATTTTGTTCA) or a negative control B antisens Gapmer from Qiagen (sequence: GCTCCCTTCAATCCAA) was used at a final concentration of 400 nM. Cells were collected 30 h after transfection in RLT lysis buffer (RNeasy Mini Kit, Qiagen) for total RNA extraction.

### CRISPR-mediated activation of *Dgkk* and oligonucleotide treatment in Neuro-2a cells

Neuro-2a cells were maintained at 37 °C in a humidified atmosphere containing 5% CO₂ in DMEM supplemented with 10% serum, 1% L-glutamine and 1% penicillin, 1% streptomycin antibiotics. A 20-nt sgRNA targeting the mouse *Dgkk* locus was designed using CHOPCHOP to maximize predicted on-target activity while minimizing potential off-target effects (Montague et al., 2014). Complementary oligonucleotides containing the *Dgkk*-targeting sequence and BpiI-compatible overhangs were synthesized: CACCGTGGTTCGGGGTCCGGCTCT and AAACCAGAGCCGGACCCCGAACCAC. Oligonucleotides were annealed, phosphorylated with T4 polynucleotide kinase, and ligated into a BpiI-linearized pUC57-attB-U6 sgRNA vector using a cloning strategy adapted from previously described CRISPR sgRNA constructs (Chen et al., 2013). Recombinant plasmids were propagated in *Escherichia coli* DH5α and verified by Sanger sequencing. For transcriptional activation of endogenous *Dgkk*, 2 × 10^5^ Neuro-2a cells per well were seeded in six-well plates. After 24 h, when cells had reached >70% confluence, each well was co-transfected with 1 µg of the *Dgkk*-targeting sgRNA construct and 1 µg of SP-dCas9-VPR, encoding a nuclease-inactive Cas9 fused to the VP64-p65-Rta transcriptional activation domains (Chavez et al., 2015), using 3 µl Lipofectamine 2000 according to the manufacturer’s instructions. Twenty-four hours after CRISPR-mediated activation, the medium was replaced and cells were transfected with the indicated oligonucleotides at a final concentration of 25 nM using a calcium phosphate method with 2 M CaCl₂ and 2× HBS. Steric block oligonucleotides (Qiagen) were used with the following sequences: CCGCTTCTGGCAGGCC (SB1), TTGGAAAACCGCTTCT (SB2), AACATTCCTTGGAAAA (SB3), TCCGAAGGAACATTCC (SB4), and CGACGACTTCCGAAGG (SB5). Cells were incubated for an additional 30 h before lysis in RLT buffer (RNeasy Mini Kit, Qiagen) for total RNA extraction.

### Mouse tissue samples

Brain tissues from *Snord116*-deficient, *Dgkk*-deficient and corresponding wild-type mice were provided by collaborating laboratories in which the respective mouse lines were maintained and experimental procedures were performed under local ethical approvals.

### Transcript abundance

Candidate transcript abundance was measured by RT-qPCR in the indicated human iPSC-derived neural differentiation stages and mouse tissues. For iPSC sample, total RNA was extracted using RNeasy mini kit (Qiagen); for mouse samples, total RNA was extracted by Trizol (Invitrogen) treatment. Then, 1µg of RNA was reverse-transcribed with oligo-dT and random hexamer primers, using Superscript IV (Thermofisher). Quantitative PCR was performed using SybrGreen (BioRad) on a QuantStudio real-time PCR system (Applied Biosystems). Relative transcript abundance was calculated using a relative standard curve method and normalized to GAPDH/Gapdh and RPLP0/Rplp0 genes. All primer pairs are described in **Supplementary Table 6**.

### NMR experiments

Milligram quantities of RNAs were prepared unlabeled by *in vitro* transcription with in-house T7 RNA polymerase from oligonucleotide templates (Milligan *et al*., 1989). The DNA templates were purchased from SIGMA-ALDRICH. Transcription conditions were optimized with varying magnesium concentrations in 25 µL reaction mixtures in 40 mM TRIS-HCl pH 8, 1 mM spermidine, 0.01% Triton X-100, 5 mM DTT and 10% polyethylene glycol (PEG8000), in the presence of 4 mM of each NTP. Preparative *in vitro* transcriptions were next performed with 16 mM MgCl_2_ in 5 mL reaction volumes. The reaction mixture was incubated at 37°C during 4.5 h. After phenol extraction and ethanol precipitation, RNAs were next purified on denaturing polyacrylamide gels followed by electroelution (Wyatt et al., 1991). After ethanol precipitation, RNAs were dialyzed overnight against the buffer used for NMR experiments. Samples were concentrated by lyophilization and resuspended in 90/10 H_2_O/D_2_O for NMR experiments. Each sample was refolded by heating at 95°C (2 min) and snap-cooled at 4°C. The concentration of RNA samples was measured using a Nanodrop Spectrometer and calculated with molar extinction coefficients. NMR experiments were recorded at 600 MHz on an Avance III Bruker spectrometer equipped with a z-gradient TXI cryoprobe. NMR data were processed using TopSpin (Bruker) and analyzed with NMRFAM-SPARKY software packages (Lee et al., 2015). NMR experiments were performed in 50 mM sodium phosphate buffer (pH 6.4). The concentration of RNA samples ranged from 100 to 340 µM. Sample volumes were 150 μL in 3 mm NMR tubes. NMR data were acquired at 15°C. Solvent suppression was achieved using combined “Jump and Return” and WATERGATE sequences (Piotto et al., 1992; Plateau et al., 1982; Sklenar et al., 1993). Two-dimensional NOESY spectra were acquired with mixing time of 50 ms and 400 ms. All titration experiments were performed at 15°C. Complexes were formed by stepwise additions of the protein to RNA monitoring the imino protons region of 1D spectra. For protein titration experiments, recombinant His6-tagged Snu13p was expressed in *Escherichia coli* Rosetta 2(DE3) cells carrying the pRARE2 plasmid and purified as described previously (Chagot et al., 2019). Following immobilized metal-affinity chromatography, the His6 tag was removed by PreScission protease cleavage, and Snu13p was further purified by size-exclusion chromatography on a Superdex 75 column in 50 mM sodium phosphate (pH 6.4), 100 mM NaCl. Peak fractions were used for NMR titrations.

### Targeted quantification of DGKκ by parallel reaction monitoring using a stable isotope-labeled internal standard peptide (PRM-SIS)

Frozen cell pellets or tissues were lysed in RIPA lysis buffer [50 mM Tris-HCl pH 7.4, 0.08 mM CaCl2, 100 mM KCl, 1 mM MgCl2, 1% (v/v) NP-40, 0.5% (m/v) sodium deoxycholate, 1% (m/v) SDS, 1× complete EDTA-free protease inhibitor cocktail]. Protein concentrations were determined using the Bradford assay (Bio-Rad protein assay dye reagent). A total of 20 μg of protein extracts was separated by SDS-PAGE on a 10 % gel for 15 minutes at 150 V. The gel was fixed for 30 min in 10% (v/v) acetic acid and 40% (v/v) ethanol, followed by staining with PageBlue Protein Staining Solution (Thermo Fisher Scientific) for 1 hour. After several washes with ultrapure water, the band corresponding to the expected molecular weight of DGKκ was excised. Gel pieces were transferred to a 96-well microplate and de-stained by three washes with 50 mM ammonium bicarbonate and 50% (v/v) acetonitrile for 10 min at room temperature with agitation at 850 rpm, followed by one wash with 100% acetonitrile under the same conditions. Proteins were reduced with 10 mM DTT for 30 minutes at 56°C and alkylated with 50 mM IAA for 30 min in the dark. Gel pieces were washed again using the same procedure and dried in a SpeedVac for 15 min. Digestion was performed overnight at 37°C with trypsin at 10 ng/μl in 10% (v/v) acetonitrile, 50 mM ammonium bicarbonate, and 2 mM CaCl2. Peptides were extracted first with 0.2% (v/v) formic acid and 50% (v/v) acetonitrile for 15 min at room temperature with agitation at 850 rpm, followed by extraction with 0.2% (v/v) formic acid and 90% (v/v) acetonitrile under the same conditions. Finally, the peptide mixture was dried in a SpeedVac for 3 h, resuspended in 20 μl of 0.1% (v/v) trifluoroacetic acid (TFA), and 0.5 μl was injected into the mass spectrometer. DGKκ-specific “signature” peptides predicted on Skyline software were validated by a global acquisition method (DDA) of the target protein produced in an overexpression system. The target peptide ISLEFNSR was synthesized chemically with heavy isotopes that incorporate 13C- and 15N-labeled C-terminal arginine (R) or lysine (K), identical in sequence and physicochemical properties to the peptides of interest, that are needed for the actual spiking and quantification of the samples. To improve the precision of absolute abundance measurements, an external calibration curve is produced by measuring the signal intensity of the reference peptides at multiple concentration points. Spike-in of samples with heavy-labeled peptides is used as internal standards. Peptide samples were analyzed by nano-liquid chromatography coupled to an Orbitrap Exploris 480 mass spectrometer (Thermo Fisher Scientific) operated in targeted PRM mode. The endogenous DGKκ peptide and its isotopically labeled counterpart were distinguished by their precursor mass shift and quantified from co-eluting fragment-ion chromatograms. PRM data were processed using Skyline. DGKκ abundance was determined from the ratio of endogenous light peptide to the corresponding stable isotope-labeled standard and the known amount of standard peptide. Calibration curves generated from defined peptide amounts were used to verify the linear quantitative range and to interpolate endogenous DGKκ peptide abundance.

### Lipidomic analysis

Hypothalamic lipidomics was performed as described previously (Habbas et al., 2022), using class-specific internal standards and high-resolution mass spectrometry. Lipid abundance was expressed as mol% of total lipids. Total and individual DAG and PA species were compared between WT, *Snord116*-KO and *Dgkk*-KO mice by one-way ANOVA followed, when appropriate, by Student-Newman-Keuls post-hoc tests.

## Statistical analysis

Statistical analyses were performed using GraphPad Prism version 11.1.0. The statistical tests used for each experiment are indicated in the corresponding figure legends

## Data availability

All data supporting the findings of this study are available within the article and its Supplementary Information. Additional data are available from the corresponding author upon reasonable request.

## Acknowledgements

We thank Jérôme Cavaillé and Timofey Rozhdestvensky for providing access to mouse tissue samples used for the PRM-SIS analyses, and Bastien Morlet from the IGBMC Mass Spectrometry Platform for assistance with these analyses. We also thank the NMR Core Facility of the Service Mutualisé de Plateformes (SMP) - Pôle BMS, Université de Lorraine, for access to NMR instrumentation. We thank the University of Connecticut Cell and Genome Engineering Core (UConn Health, Farmington, CT, USA) for providing the iPSC lines.

## Funding

This work was supported by the Centre National de la Recherche Scientifique (CNRS), SATT SAYENS (to S.L.), Foundation for Prader-Willi Research (FPWR) and Fondation Jérôme Lejeune and Foundation for Prader-Willi Research (to H.M. and S.L.), and Fondation ID+ Lorraine (to S.L.).

## Competing interests

The authors declare no competing interests.

## Legend of supplementary figure

**Supplementary Figure 1. Evolutionary persistence of predicted *SNORD116*-mRNA interactions across eutherian mammals.** Predicted duplexes between *SNORD116* and the selected candidate target sites are shown across the 16 eutherian species included in the comparative analysis. For each species, the most favorable interaction identified between the orthologous target sequence and the corresponding *SNORD116* repertoire is represented. Species sharing the same predicted duplex are grouped together. A slash indicates that no interaction meeting the selection criteria was identified.

**Supplementary Figure 2. Candidate transcript responses across complementary models of *SNORD116/Snord116* loss.** Individual RT-qPCR measurements underlying the summary shown in Figure 4 are presented for the indicated candidate transcripts. Human candidate expression was measured in MCH2-10 and YK26 cells at the neural progenitor (NPRO, n=8-10), neuronal precursor (NPRE, n=8-10) and neuronal (NEUR, n=8-15 for MCH2-10, n=3 for YK26) stages following *SNORD116* depletion. Mouse orthologues were analyzed in *Snord116*-deficient and WT whole brain at PND3 (n=6) and PND7 (n=6) and hypothalamus at PND21 (n=7). Data are presented as means ± SD. * P<0.05, **P < 0.01, ***P < 0.001 in T-test for YK26 neurons and Mann-Whitney test for the other conditions. For mouse comparisons, isolated outliers identified by Rout test were removed: one sample for *Adarb1* in KO at PND21, and one sample for *Irx5* in KO at PND7. Their exclusion did not change the conclusion of the corresponding WT versus *Snord116*-KO comparison.

**Supplementary Figure 3. S*n*ord116 levels following oligonucleotide treatment in Neuro-2a cells**. (A) Relative *Snord116* RNA abundance following treatment with the G116 Gapmer or control oligonucleotide (n=7-10). (B) Relative *Snord116* RNA abundance following treatment with steric-blocking oligonucleotides SB1-SB5 or control oligonucleotide (8-10). Data are presented as means ± SD. Mann-Whitney test * P<0.05. For mouse comparisons, one sample in CTR and one sample in G116 were identified as outliers by Rout test and removed; their exclusion did not change the conclusion of the analysis.

**Supplementary Figure 4. PRM-SIS validation of DGKκ quantification in brain extracts.** (A) Fragment-ion spectrum of the mouse DGKκ signature peptide ISLEFNSR, showing the assigned fragment ions. (B) Genetic validation of DGKκ detection using WT and *Dgkk*-KO hypothalamic extracts. Chromatographic signals for the stable isotope-labeled reference peptide and endogenous peptide are shown. (C) Calibration curve used for absolute quantification of the endogenous DGKκ peptide.

**Supplementary Figure 5. Lipidomic analysis of Snord116-KO hypothalamus.** (A) Lipid-class composition of the analyzed samples, expressed as percentage of total lipids. (B) Total diacylglycerol (DAG) abundance in hypothalamic extracts from WT, *Snord116*-KO and *Dgkk*-KO mice. (C) Total phosphatidic acid (PA) abundance. (D) Individual DAG species. (E) Individual PA species. Data are expressed as mol% of total lipids and shown as mean ± SEM. Homogeneity of variance was assessed using the Fisher–Snedecor test. For each lipid class or species, groups were compared by ANOVA; when significant (α = 0.05), Student-Newman-Keuls post hoc tests were applied using a corrected significance threshold of α/3.

**Supplementary Figure 6. Expression of differentiation markers during neural differentiation of human iPSCs.** Expression of pluripotency markers (*OCT4, NANOG*), neural progenitor markers (*PAX6, NES*), neuronal development markers (*TUBB3, MAP2*) and the forebrain identity marker *FOXG1* was measured at the undifferentiated (UNDIFF), neural progenitor (NPRO), neuronal precursor (NPRE) and neuronal (NEUR) stages in MCH2-10 (A) and YK26 (B) cells. Data are shown as relative RNA levels. Mean ± SD of n=3-4 independent samples analyzed by T-test.

## Notes

### Competing Interest Statement

The authors have declared no competing interest.

## References

Baldini L, Charpentier B, Labialle S. 2021. Emerging Data on the Diversity of Molecular Mechanisms Involving C/D snoRNAs. Non-Coding RNA 7(2):30. doi:10.3390/ncrna7020030.

Baldini L, Robert A, Charpentier B, Labialle S. 2022. Phylogenetic and molecular analyses identify SNORD116 targets involved in the Prader-Willi syndrome. Molecular Biology and Evolution 39(1):msab348. doi:10.1093/molbev/msab348.

Batzli K et al. 2026. Role of Snord116 in pituitary growth hormone deficiency of Prader-Willi syndrome. JCI Insight 11(18):e205467. doi:10.1172/jci.insight.205467.

Bergeron D, Fafard-Couture É, Scott MS. 2020. Small nucleolar RNAs: continuing identification of novel members and increasing diversity of their molecular mechanisms of action. Biochemical Society Transactions 48(2):645–656. doi:10.1042/BST20191046.

Bochukova EG et al. 2018. A transcriptomic signature of the hypothalamic response to fasting and BDNF deficiency in Prader-Willi syndrome. Cell Reports 22(13):3401–3408. doi:10.1016/j.celrep.2018.03.018.

Bortolin-Cavaillé M-L, Cavaillé J. 2012. The SNORD115 (H/MBII-52) and SNORD116 (H/MBII-85) gene clusters at the imprinted Prader–Willi locus generate canonical box C/D snoRNAs. Nucleic Acids Research 40(14):6800–6807. doi:10.1093/nar/gks321.

Cavaillé J, Nicoloso M, Bachellerie JP. 1996. Targeted ribose methylation of RNA in vivo directed by tailored antisense RNA guides. Nature 383:732–735. doi:10.1038/383732a0.

Chagot ME, Quinternet M, Rothé B, et al. 2019. The yeast C/D box snoRNA U14 adopts a “weak” K-turn-like conformation recognized by the Snu13 core protein in solution. Biochimie 164:70–82. doi:10.1016/j.biochi.2019.03.014.

Chamberlain SJ, Chen PF, Ng KY, et al. 2010. Induced pluripotent stem cell models of the genomic imprinting disorders Angelman and Prader-Willi syndromes. Proceedings of the National Academy of Sciences USA 107(41):17668–17673. doi:10.1073/pnas.1004487107.

Chavez A, Scheiman J, Vora S, et al. 2015. Highly efficient Cas9-mediated transcriptional programming. Nature Methods 12:326–328. doi:10.1038/nmeth.3312.

Chen B, Gilbert LA, Cimini BA, et al. 2013. Dynamic imaging of genomic loci in living human cells by an optimized CRISPR/Cas system. Cell 155:1479–1491. doi:10.1016/j.cell.2013.12.001.

Chen CL, Perasso R, Qu LH, Amar L. 2007. Exploration of pairing constraints identifies a 9 base-pair core within box C/D snoRNA-rRNA duplexes. Journal of Molecular Biology 369(3):771–783. doi:10.1016/j.jmb.2007.03.052.

de Smith AJ, Purmann C, Walters RG, et al. 2009. A deletion of the HBII-85 class of small nucleolar RNAs (snoRNAs) is associated with hyperphagia, obesity and hypogonadism. Human Molecular Genetics 18:3257–3265. doi:10.1093/hmg/ddp263.

Deschamps-Francoeur G, Couture S, Abou-Elela S, Scott MS. 2022. The snoGloBe interaction predictor reveals a broad spectrum of C/D snoRNA RNA targets. Nucleic Acids Research 50(11):6067–6083. doi:10.1093/nar/gkac475.

Ding F, Li HH, Zhang S, et al. 2008. SnoRNA Snord116 (Pwcr1/MBII-85) deletion causes growth deficiency and hyperphagia in mice. PLoS ONE 3(3):e1709. doi:10.1371/journal.pone.0001709.

Dragon F, Gallagher JEG, Compagnone-Post PA, et al. 2002. A large nucleolar U3 ribonucleoprotein required for 18S ribosomal RNA biogenesis. Nature 417:967–970. doi:10.1038/nature00769.

Duker AL, Ballif BC, Bawle EV, et al. 2010. Paternally inherited microdeletion at 15q11.2 confirms a significant role for the SNORD116 C/D box snoRNA cluster in Prader-Willi syndrome. European Journal of Human Genetics 18(11):1196–1201. doi:10.1038/ejhg.2010.102.

Dupuis-Sandoval F, Poirier M, Scott MS. 2015. The emerging landscape of small nucleolar RNAs in cell biology. WIREs RNA 6(4):381–397. doi:10.1002/wrna.1284.

Falaleeva M, Pages A, Matuszek Z, et al. 2016. Dual function of C/D box small nucleolar RNAs in rRNA modification and alternative pre-mRNA splicing. Proceedings of the National Academy of Sciences USA 113(12):E1625–E1634. doi:10.1073/pnas.1519292113.

Gilmore RB, Liu Y, Stoddard CE, et al. 2024. Identifying key underlying regulatory networks and predicting targets of orphan C/D box SNORD116 snoRNAs in Prader-Willi syndrome. Nucleic Acids Research 52(22):13757–13774. doi:10.1093/nar/gkae1129.

Guibert M, Marty-Capelle H, Robert A, et al. 2024. Coordinated evolution of the SNORD115 and SNORD116 tandem repeats at the imprinted Prader-Willi/Angelman locus. NAR Molecular Medicine 1(1):ugad003. doi:10.1093/narmme/ugad003.

Habbas K, Cakil O, Zámbó B, et al. 2022. AAV-delivered diacylglycerol kinase DGKκ achieves long-term rescue of fragile X syndrome mouse model. EMBO Molecular Medicine 14(5):e14649. doi:10.15252/emmm.202114649.

Helwak A, Turowski T, Spanos C, Tollervey D. 2024. Roles of SNORD115 and SNORD116 ncRNA clusters during neuronal differentiation. Nature Communications 15:10427. doi:10.1038/s41467-024-54573-8.

Henras AK, Plisson-Chastang C, Humbert O, Romeo Y, Henry Y. 2017. Synthesis, function, and heterogeneity of snoRNA-guided posttranscriptional nucleoside modifications in eukaryotic ribosomal RNAs. The Enzymes 41:169–213. doi:10.1016/bs.enz.2017.03.007.

Huang C, Shi J, Guo Y, et al. 2017. A snoRNA modulates mRNA 3′-end processing and regulates the expression of a subset of mRNAs. Nucleic Acids Research 45(15):8647–8660. doi:10.1093/nar/gkx651.

Kass S, Tyc K, Steitz JA, Sollner-Webb B. 1990. The U3 small nucleolar ribonucleoprotein functions in the first step of preribosomal RNA processing. Cell 60(6):897–908. doi:10.1016/0092-8674(90)90338-F.

Kiss-László Z, Henry Y, Bachellerie JP, Caizergues-Ferrer M, Kiss T. 1996. Site-specific ribose methylation of preribosomal RNA: a novel function for small nucleolar RNAs. Cell 85(7):1077–1088. doi:10.1016/S0092-8674(00)81308-2.

Knott B, Kocher MA, Paz HA, et al. 2022. Dietary conjugated linoleic acid reduces body weight and fat in Snord116m+/p− and Snord116m−/p− mouse models of Prader-Willi syndrome. Nutrients 14(4):860. doi:10.3390/nu14040860.

Kocher MA, Huang FW, Le E, Good DJ. 2021. Snord116 post-transcriptionally increases Nhlh2 mRNA stability: implications for human Prader-Willi syndrome. Human Molecular Genetics 30(12):1101–1110. doi:10.1093/hmg/ddab103.

Kumar S, Stecher G, Suleski M, Hedges SB. 2017. TimeTree: a resource for timelines, timetrees, and divergence times. Molecular Biology and Evolution 34(7):1812–1819. doi:10.1093/molbev/msx116.

Kummerfeld DM, Skryabin BV, Brosius J, Vakhrushev SY, Rozhdestvensky TS. 2022. Reference genes across nine brain areas of wild type and Prader-Willi syndrome mice: assessing differences in Igfbp7, Pcsk1, Nhlh2 and Nlgn3 expression. International Journal of Molecular Sciences 23(15):8729. doi:10.3390/ijms23158729.

Lee W, Tonelli M, Markley JL. 2015. NMRFAM-SPARKY: enhanced software for biomolecular NMR spectroscopy. Bioinformatics 31:1325–1327. doi:10.1093/bioinformatics/btu830.

Liu B, Wu T, Miao BA, et al. 2025. snoRNA-facilitated protein secretion revealed by transcriptome-wide snoRNA target identification. Cell 188(2):465–483.e22. doi:10.1016/j.cell.2024.10.046.

Lorenz R, Bernhart SH, Höner zu Siederdissen C, et al. 2011. ViennaRNA Package 2.0. Algorithms for Molecular Biology 6:26. doi:10.1186/1748-7188-6-26.

Mann M, Wright PR, Backofen R. 2017. IntaRNA 2.0: enhanced and customizable prediction of RNA-RNA interactions. Nucleic Acids Research 45(W1):W435–W439. doi:10.1093/nar/gkx279.

Marty V, Hebras J, Boursereau R, et al. 2026. Evolutionary and biological insights into the paternally expressed Snord116-Ipw-Snord115 gene array at the imprinted Prader-Willi syndrome domain. bioRxiv. doi:10.64898/2026.07.28.739784.

Milligan JF, Uhlenbeck OC. 1989. Synthesis of small RNAs using T7 RNA polymerase. Methods in Enzymology 180:51–62.

Montague TG, Cruz JM, Gagnon JA, Church GM, Valen E. 2014. CHOPCHOP: a CRISPR/Cas9 and TALEN web tool for genome editing. Nucleic Acids Research 42:W401–W407. doi:10.1093/nar/gku410.

Piotto M, Saudek V, Sklenár V. 1992. Gradient-tailored excitation for single-quantum NMR spectroscopy of aqueous solutions. Journal of Biomolecular NMR 2:661–665.

Plateau P, Gueron M. 1982. Exchangeable proton NMR without base-line distortion, using new strong-pulse sequences. Journal of the American Chemical Society 104:7310–7311. doi:10.1021/ja00389a067.

Rivière JC, Marty V, Cavaillé J, Verret L. 2026. Loss of Snord115 mitigates Snord116-driven sleep abnormalities in mouse models of Prader-Willi syndrome. Neurobiology of Disease 228:107551. doi:10.1016/j.nbd.2026.107551.

Sahoo T, del Gaudio D, German JR, et al. 2008. Prader-Willi phenotype caused by paternal deficiency for the HBII-85 C/D box small nucleolar RNA cluster. Nature Genetics 40:719–721. doi:10.1038/ng.158.

Sjöstedt E, Zhong W, Fagerberg L, et al. 2020. An atlas of the protein-coding genes in the human, pig, and mouse brain. Science 367(6482):eaay5947. doi:10.1126/science.aay5947.

Sklenar V, Piotto M, Leppik R, Saudek V. 1993. Gradient-tailored water suppression for 1H-15N HSQC experiments optimized to retain full sensitivity. Journal of Magnetic Resonance, Series A 102:241–245. doi:10.1006/jmra.1993.1098.

Tabet R, Moutin E, Becker JAJ, et al. 2016. Fragile X Mental Retardation Protein (FMRP) controls diacylglycerol kinase activity in neurons. Proceedings of the National Academy of Sciences USA 113(26):E3619–E3628. doi:10.1073/pnas.1522631113.

Tang Y, Wu Y, Wang S, Lu X, Gu X, Li Y, Yang F, Xu R, Wang T, Jiao Z, Wu Y, Liu L, Chen JQ, Wang Q, Chen Q. 2024. An integrative platform for detection of RNA 2’-O-methylation reveals its broad distribution on mRNA. Cell Reports Methods 4(3):100721. doi:10.1016/j.crmeth.2024.100721.

Tauber M, Höybye C. 2021. Endocrine disorders in Prader-Willi syndrome: a model to understand and treat hypothalamic dysfunction. Lancet Diabetes & Endocrinology 9(4):235–246. doi:10.1016/S2213-8587(21)00002-4.

van Nues RW, Watkins NJ. 2017. Unusual C′ /D′ motifs enable box C/D snoRNPs to modify multiple sites in the same rRNA target region. Nucleic Acids Research 45(4):2016–2028. doi:10.1093/nar/gkw842.

Watkins NJ, Bohnsack MT. 2012. The box C/D and H/ACA snoRNPs: key players in the modification, processing and dynamic folding of ribosomal RNA. WIREs RNA 3(3):397–414. doi:10.1002/wrna.117.

Wyatt JR, Chastain M, Puglisi JD. 1991. Synthesis and purification of large amounts of RNA oligonucleotides. BioTechniques 11:764–769.

Yang Z, Lin J, Ye K. 2016. Box C/D guide RNAs recognize a maximum of 10 nt of substrates. Proceedings of the National Academy of Sciences USA 113(39):10878–10883. doi:10.1073/pnas.1604872113.

Zeng H, Guo M, Martins-Taylor K, et al. 2010. Specification of region-specific neurons including forebrain glutamatergic neurons from human induced pluripotent stem cells. PLoS ONE 5(7):e11853. doi:10.1371/journal.pone.0011853.

