## Supplementary Figures for "The *DGKK* mRNA is a primary target of the Prader-Willi syndrome-associated snoRNA *SNORD116*"

Supplementary Figure 1

### ADARB1/Adarb1 (site1)

**Human**

```
Adarb1    5'–CAAUGGGGGUGGUGGU–3'
             ||||||||||||||||
Snord116  3'–GUUACUCCUACUACCA–5'
```

**-25.79 kcal/mol**

**Chimpanzee**

```
Adarb1    5'–CCAUCAUCCUCAUUGU–3'
             ||||||||||||||||
Snord116  3'–GGUGGUGGGGGUAACA–5'
```

**-25.79 kcal/mol**

**Rhesus macaque**

```
Adarb1    5'–CAAUGGGGGUGGUGGU–3'
             ||||||||||||||||
Snord116  3'–GUUACUCCUACUACCA–5'
```

**-22.37 kcal/mol**

**Marmoset**

```
Adarb1    5'–AUGGGGGUGGUGG–3'
             |||||||||||||
Snord116  3'–UACUCCUACUGCC–5'
```

**-16.58 kcal/mol**

**Brown bat**

```
Adarb1    5'–AUGGGGGUG–3'
             |||||||||
Snord116  3'–UACCCUUAC–5'
```

**-12.69 kcal/mol**

Greater Galago, Guinea pig, Hedgehog, Kangaroo rat, Mouse, Mouse lemur, Northern treeshrew, Pig, Rabbit, Rat, Squirrel

/

### ADARB1/Adarb1 (Site 2)

**Human**

```
Adarb1    5'–UUUCGGGGAGCC–3'
             ||||||||||||
Snord116  3'–AAGGUUCCUUGG–5'
```

**-15.84 kcal/mol**

**Marmoset**

```
                      G
Adarb1    5'–UUCAGGGA  GC–3'
             ||||||||  ||
Snord116  3'–AGGUUCCU  CG–5'
```

**-11.77 kcal/mol**

**Squirrel**

```
Adarb1    5'–UUUCCAAGGAAU–3'
             ||||||||||||
Snord116  3'–AAAGGUUCCUUA–5'
```

**-16.13 kcal/mol**

**Guinea Pig**

```
Adarb1    5'–UUCCAAGGAAUG–3'
             ||||||||||||
Snord116  3'–AAGGUUCCUUAC–5'
```

**-18.20 kcal/mol**

**Rabbit**

```
Adarb1    5'–UCCAAGGAAUGU–3'
             ||||||||||||
Snord116  3'–AGGUUCCUUACA–5'
```

**-18.80 kcal/mol**

**Pig**

```
Adarb1    5'–GGGGAGGCGG–3'
             ||||||||||
Snord116  3'–CUCCUCUGCC–5'
```

**-19.46 kcal/mol**

Brown bat, Chimpanzee, Greater Galago, Hedgehog, Kangaroo rat, Mouse lemur, Mouse, Northern treeshrew, Rat, Rhesus macaque

/

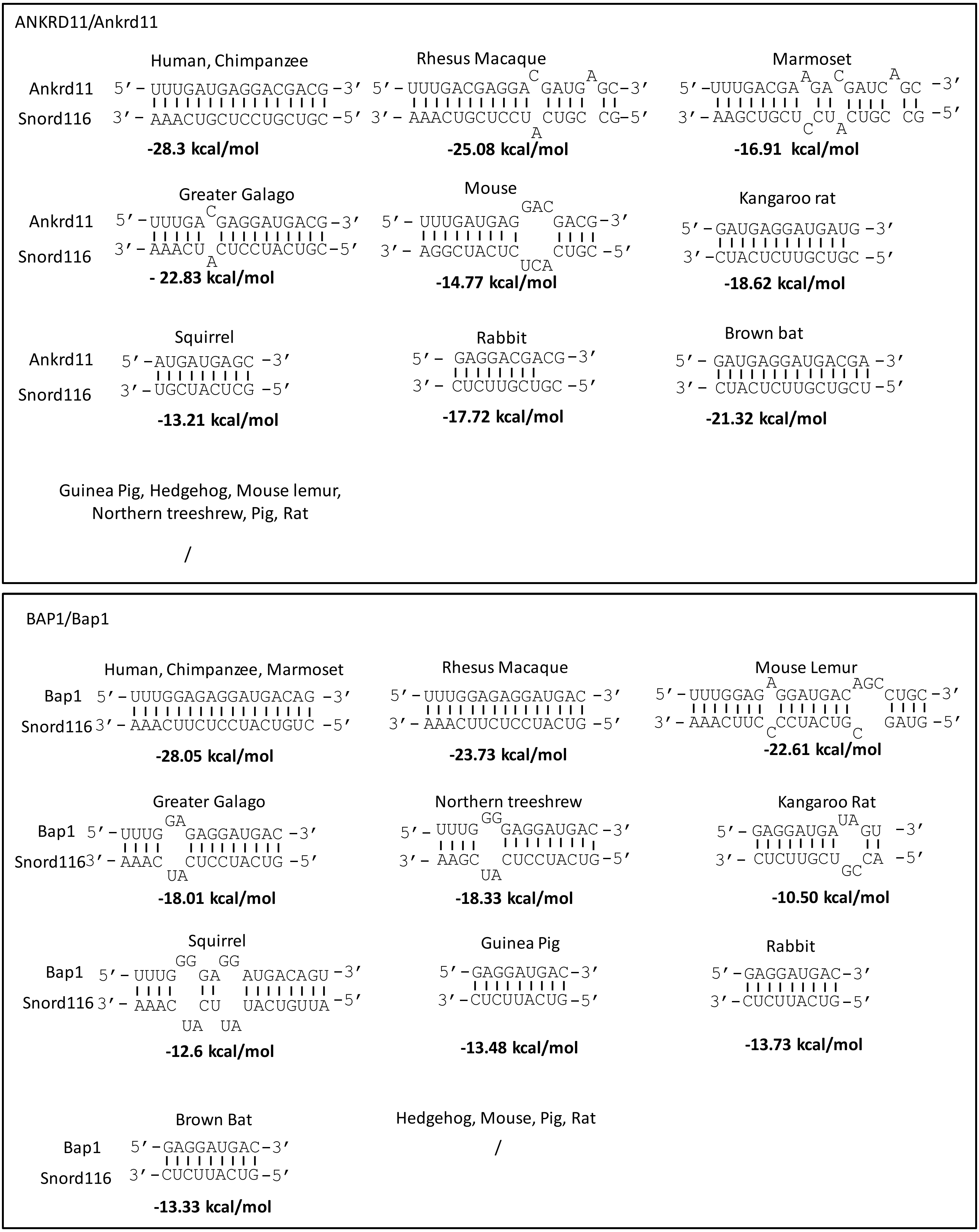
ANKRD11/Ankrd11
Human, Chimpanzee
Ankrd11 5′–UUUGAUGAGGACGACG–3′
Snord116 3′–AAACUGCUCCUGCUGC–5′
-28.3 kcal/mol
Rhesus Macaque
5′–UUUGACGAGGA C GAUG A GC–3′
3′–AAACUGCUCCU A CUGC CG–5′
-25.08 kcal/mol
Marmoset
5′–UUUGACGA A GA C GAUC A GC –3′
3′–AAGCUGCU C CU A CUGC CG –5′
-16.91 kcal/mol
Greater Galago
Ankrd11 5′–UUUGA C GAGGAUGACG–3′
Snord116 3′–AAACU A CUCCUACUGC–5′
- 22.83 kcal/mol
Mouse
5′–UUUGAUGAG GAC GACG–3′
3′–AGGCUACUC UCA CUGC–5′
-14.77 kcal/mol
Kangaroo rat
5′–GAUGAGGAUGAUG–3′
3′–CUACUCUUGCUGC–5′
-18.62 kcal/mol
Squirrel
Ankrd11 5′–AUGAUGAGC –3′
Snord116 3′–UGCUACUCG –5′
-13.21 kcal/mol
Rabbit
5′– GAGGACGACG–3′
3′– CUCUUGCUGC–5′
-17.72 kcal/mol
Brown bat
5′– GAUGAGGAUGACGA –3′
3′– CUACUCUUGCUGCU–5′
-21.32 kcal/mol
Guinea Pig, Hedgehog, Mouse lemur, Northern treeshrew, Pig, Rat
/
BAP1/Bap1
Human, Chimpanzee, Marmoset
Bap1 5′–UUUGGAGAGGAUGACAG –3′
Snord116 3′–AAACUUCUCCUACUGUC –5′
-28.05 kcal/mol
Rhesus Macaque
5′–UUUGGAGAGGAUGAC –3′
3′–AAACUUCUCCUACUG –5′
-23.73 kcal/mol
Mouse Lemur
5′–UUUGGAG A GGAUGAC AGC CUGC–3′
3′–AAACUUC C CCUACUG C GAUG –5′
-22.61 kcal/mol
Greater Galago
Bap1 5′–UUUG GA GAGGAUGAC –3′
Snord116 3′–AAAC UA CUCCUACUG –5′
-18.01 kcal/mol
Northern treeshrew
5′– UUUG GG GAGGAUGAC–3′
3′– AAGC UA CUCCUACUG–5′
-18.33 kcal/mol
Kangaroo Rat
5′– GAGGAUGA UA GU –3′
3′– CUCUUGCU GC CA –5′
-10.50 kcal/mol
Squirrel
Bap1 5′– UUUG GG GA GG AUGACAGU –3′
Snord116 3′– AAAC UA CU UA UACUGUUA –5′
-12.6 kcal/mol
Guinea Pig
5′–GAGGAUGAC–3′
3′–CUCUUACUG–5′
-13.48 kcal/mol
Rabbit
5′–GAGGAUGAC–3′
3′–CUCUUACUG–5′
-13.73 kcal/mol
Brown Bat
Bap1 5′–GAGGAUGAC–3′
Snord116 3′–CUCUUACUG–5′
-13.33 kcal/mol
Hedgehog, Mouse, Pig, Rat
/

### C1QTNF/C1qtnf9

Human
```
C1qtnf9   5'- UGAUGAGGACGAUG -3'
              ||||||||||||||
Snord116  3'- ACUGCUCCUGCUGC -5'
```
**-25.17 kcal/mol**

Chimpanzee
```
5'- UGAUGAGGACGAUG-3'
    ||||||||||||||
3'- ACUGCUCCUGCUGC-5'
```
**-25.10 kcal/mol**

Rhesus Macaque
```
5'- UGAUGAGGACGAUG-3'
    ||||||||||||||
3'- ACUGCUCCUGCUGC-5'
```
**-25.55 kcal/mol**

Marmoset
```
                        C
C1qtnf9   5'- CCGAUGAGGA GAUG -3'
              |||||||||| ||||
Snord116  3'- GGCUACUCCU CUGC -5'
                        U
```
**-24.14 kcal/mol**

Mouse Lemur
```
5'-CGAUGAGGAUGACG-3'
   ||||||||||||||
3'-GCUACUCCUACUGC-5'
```
**-27.22 kcal/mol**

Greater Galago
```
5'-CGAUGAGGACGACG-3'
   ||||||||||||||
3'-GCUACUCCUGCUGC-5'
```
**-28.62 kcal/mol**

Northern treeshrew
```
C1qtnf9   5'- GAUGAGGAUGAUG -3'
              |||||||||||||
Snord116  3'- CUACUCCUACUGC -5'
```
**-22.7 kcal/mol**

Rat
```
5'- GAUGACGAUA -3'
    ||||||||||
3'- CUACUGUUAU -5'
```
**-12.17 kcal/mol**

Kangaroo rat
```
       C
5'-CGA GAGGACGACG -3'
   |||  ||||||||||
3'-GCU CUCUUGCUGC -5'
      A
```
**-20.86 kcal/mol**

Rabbit
```
C1qtnf9   5'- CGAUGAGGACGACG -3'
              ||||||||||||||
Snord116  3'- GCUACUCUUGCUGC -5'
```
**-25.17 kcal/mol**

Pig
```
       G  G        C
5'-GUUU CG AUGAGGA GACG-3'
   |||| || ||||||| ||||
3'-CAAA GC UACUCCU CUGC-5'
```
**-20.83 kcal/mol**

Brown bat
```
        G
5'-UUGC GAUGAGGAUGAUGA -3'
   ||||  ||||||||||||||
3'-AAUG CUACUCUUGCUGCU -5'
        G
```
**-25.22 kcal/mol**

Squirrel
```
                     UGA
C1qtnf9   5'- UUGCAGA   GG -3'
              |||||||   ||
Snord116  3'- AGUGUCU   CC -5'
                     UCA
```
**-8.71kcal/mol**

Guinea Pig, Hedgehog, Mouse

/

### CADPS2/Cadps2

Human, Chimpanzee
```
Cadps2    5'- UUCGCUGAGAAUGAUGGU -3'
              ||||||||||||||||||
Snord116  3'- AGGCGACUCUUGCUGCCA -5'
```
**-29.83 kcal/mol**

Brown bat, Greater Galago, Guinea pig, Hedgehog, Kangaroo rat, Marmoset, Mouse lemur, Mouse, Northern treeshrew, Pig, Rabbit, Rat, Rhesus macaque, Squirrel

/

### CHRNA7/Chrna7

Human
```
Chrna7    5'- GAUGGGGGUGAUGGUGUAG -3'
              |||||||||||||||||||
Snord116  3'- CUGCUCCUACUGCCAUAUC -5'
```
**-31.53 kcal/mol**

Chimpanzee
```
5'-AUGGGGGUGAUGGUGUAG -3'
   ||||||||||||||||||
3'-UACUCCUACUACCAUAUC -5'
```
**-28.73 kcal/mol**

Rhesus Macaque
```
5'- GAUGGGGGUGAUGGU-3'
    |||||||||||||||
3'- CUGCUCCUACUGCCA-5'
```
**-24.5 kcal/mol**

Squirrel
```
              A
Chrna7    5'-GAU GGAUGUGAUGGUGUAG -3'
             ||| ||||||||||||||||
Snord116  3'-CUA CUUAUACUGUUACAUC -5'
```
**-20.83 kcal/mol**

Brown bat, Chimpanzee, Greater Galago, Guinea pig, Hedgehog, Kangaroo rat, Marmoset, Mouse lemur, Mouse, Northern treeshrew, Pig, Rabbit, Rat

/

DGKK/Dgkk

Human, Chimpanzee
```
Dgkk      5'-UUUUCCAAGGAAUGUU -3'
             ||||||||||||||||
Snord116  3'-AAAAGGUUCCUUACAA -5'
```
**-22.2 kcal/mol**

Rhesus Macaque
```
5'-UUUUCCAAGGAAUGUU -3'
   ||||||||||||||||
3'-AAAAGGUUCCUUACAA -5'
```
**-22.08 kcal/mol**

Marmoset
```
5'-UUUUCCAAGGAAUGUU -3'
   ||||||||||||||||
3'-AAAAGGUUCCUUACAA -5'
```
**-22.09 kcal/mol**

Mouse Lemur
```
Dgkk      5'-UUUUCCAAGGAAUGUU -3'
             ||||||||||||||||
Snord116  3'-AAAAGGUUCCUUACAA -5'
```
**-22.06 kcal/mol**

Greater Galago
```
5'-UUUUCCAAGGAAUGUU -3'
   ||||||||||||||||
3'-AAAAGGUUCCUUACAA -5'
```
**-22.38 kcal/mol**

Northern treeshrew
```
5'-UUUUCCAAGGAAUGUU -3'
   ||||||||||||||||
3'-AAAAGGUUCCUUACAA -5'
```
**-21.67 kcal/mol**

Mouse
```
Dgkk      5'-UUUUCCAAGGAAUGUU -3'
             ||||||||||||||||
Snord116  3'-AAAAGGUUCCUUACAA -5'
```
**-22.03 kcal/mol**

Rat
```
5'-UUUUCCAAGGAAUGUU -3'
   ||||||||||||||||
3'-AAAAGGUUCCUUACAA -5'
```
**-21.75 kcal/mol**

Kangaroo rat
```
5'-UUUUCCAAGGAAUGUU -3'
   ||||||||||||||||
3'-AAAAGGUUCCUUACAA -5'
```
**-22.34 kcal/mol**

Squirrel
```
Dgkk      5'-UUUUCCAAGGAAUGUU -3'
             ||||||||||||||||
Snord116  3'-AAAAGGUUCCUUACAA -5'
```
**-22.04 kcal/mol**

Guinea pig
```
5'- UUUUCCAAGGAAUGUU -3'
    ||||||||||||||||
3'- AAAAGGUUCCUUACAA -5'
```
**-22.10 kcal/mol**

Rabbit
```
5'-UUUUCCAAGGAAUGUU -3'
   ||||||||||||||||
3'-AAAAGGUUCCUUACAA -5'
```
**-22.37 kcal/mol**

Pig
```
Dgkk      5'-UUUCCAAGGAAUGCU -3'
             |||||||||||||||
Snord116  3'-AAAGGUUCCUUACGA -5'
```
**-23.83 kcal/mol**

Hedgehog
```
5'- UUUUCCAAGGAAUGCU -3'
    ||||||||||||||||
3'- AAAAGGUUCCUUACGA -5'
```
**-24.73 kcal/mol**

Brown bat
```
5'-UUUUCCAAGGAAUGUU -3'
   ||||||||||||||||
3'-AAAAGGUUCCUUACAA -5'
```
**-22.08 kcal/mol**

DPP10/Dpp10

Human, Chimpanzee
```
Dpp10     5'-UUUGCUGAGAAUGAUGGUA-3'
             |||||||||||||||||||
Snord116  3'-AGGCGACUCUUGCUGCCAU-5'
```
**-29.27 kcal/mol**

Brown bat, Greater Galago, Guinea pig, Hedgehog, Kangaroo rat, Marmoset, Mouse lemur, Mouse, Northern treeshrew, Pig, Rabbit, Rat, Rhesus macaque, Squirrel

/

FGF13/Fgf13

Human
```
                    U
Fgf13     5'-GUGGCAC GAAGUGAU -3'
             ||||||| ||||||||
Snord116  3'-CACCGUG CUUCACUA -5'
                    U
```
**-25.59 kcal/mol**

Chimpanzee
```
          U
5'-GUGGCAC GAAGUGAU -3'
   ||||||| ||||||||
3'-CACCGUG CUUCACUA -5'
          U
```
**-25.61 kcal/mol**

Rhesus Macaque
```
       CAC       U
5'-GAAG   GGCAC GAAGUGAUU -3'
   ||||   ||||| |||||||||
3'-CUUC   CCGUG CUUCACUAA -5'
       A         U
```
**-25.23 kcal/mol**

Marmoset
```
                   UG
Fgf13     5'-GUGGCAC  AAGUGAUU -3'
             |||||||  ||||||||
Snord116  3'-CACCGUG  UUUACUAA -5'
                   UU
```
**-21.50 kcal/mol**

Northern treeshrew
```
5'-GUG UGGUACUGAG-3'
   ||| ||||||||||
3'-CAC ACCAUGACUC-5'
      U
```
**-20.33 kcal/mol**

Rat
```
              ACU
5'-AAUGGUG       GUGC-3'
   |||||||       ||||
3'-UUAUCGC       CACG-5'
```
**-9.25 kcal/mol**

Brown bat, Greater Galago, Guinea pig, Hedgehog, Kangaroo rat, Mouse lemur, Mouse, Pig, Rabbit, Squirrel

/

IRX5/Irx5 (site 1)

Human

```
            U     C
Irx5      5'-UGACG GGACG CG -3'
             ||||| ||||| ||
Snord116  3'-ACUGC CCUGC GC -5'
                 U     U
```

**-18.3 kcal/mol**

Greater Galago

```
          CGU      CC
5'- GAUGA    GGACG   GCGG-3'
    |||||    |||||   ||||
3'- CUACU    CCUGC   UGCC-5'
```

**-17.3 kcal/mol**

Mouse

```
                 A  CC
5'-GAUGAC GUGG CG   ACGGA-3'
   |||||| |||| ||   |||||
3'-CUACUG CACC GC   UGUCU-5'
         U        CA
```

**-21.39 kcal/mol**

Rat

```
                  C     A
Irx5      5'-AGAUGA GUGG CG-3'
             ||||||  |||| ||
Snord116  3'-UCUACU  CACC GC-5'
                   AU
```

**-12.85 kcal/mol**

Kangaroo rat

```
          C     C CC
5'-GAUGA GUGGA G  GCGG-3'
   ||||| ||||| |  ||||
3'-CUACU CGCCU C  UGCC-5'
```

**-16.17 kcal/mol**

Guinea pig

```
5'- GUG GACGCCG CG -3'
    ||| ||||||| ||
3'- CAC CUGUGGU GU -5'
       A       A
```

**-14.92 kcal/mol**

Rabbit

```
              A      CGU       CC
Irx5      5'-CA GAUGA   GGACG   GCGG-3'
             || |||||   |||||   ||||
Snord116  3'-GU CUACU   CUUGC   UGCC-5'
               G
```

**-15.6 kcal/mol**

Pig

```
          C U
5'- GAUGA G GGACG-3'
    ||||| | |||||
3'- CUACU C UCUGC-5'
           C
```

**-12.37 kcal/mol**

Hedgehog

```
        CC      CC C
5'-GAUGA  UGGACG  G GG-3'
   |||||  ||||||  | ||
3'-CUACU  AUCUGC  C CC-5'
        CU      CU
```

**-13.2 kcal/mol**

Brown bat, Chimpanzee, Marmoset, Mouse lemur, Northern treeshrew, Rhesus macaque, Squirrel

/

IRX5/Irx5 (site 2)

Human, Chimpanzee

```
Irx5      5'- GACGAGGACGA -3'
              |||||||||||
Snord116  3'- CUGCUCCUGCU -5'
```

**-21.84 kcal/mol**

Rhesus macaque

```
5'-CGACGAGGA-3'
   |||||||||
3'-GCUGCUCCU-5'
```

**-16.28 kcal/mol**

Marmoset

```
5'-CGACGAGGA-3'
   |||||||||
3'-GCUGCUCCU-5'
```

**-16.22 kcal/mol**

Mouse Lemur

```
                C
Irx5      5'- GAA GACGAGGA -3'
              |||  ||||||||
Snord116  3'- CUU CUGUUCCU -5'
```

**-14.32 kcal/mol**

Greater Galago

```
       C
5'- GA GAGGACGA -3'
    ||  ||||||||
3'- CU CUCCUGCU -5'
      A
```

**-16.4 kcal/mol**

Kangaroo rat

```
5'- AGAACGACG -3'
    |||||||||
3'- UCUUGCUGC -5'
```

**-14.04 kcal/mol**

Guinea pig

```
Irx5      5'- AGAAUGACG-3'
              |||||||||
Snord116  3'- UCUUACUGC-5'
```

**-12.61 kcal/mol**

Brown bat

```
           G
5'-GAGGACGA GA GG-3'
   |||||||| || ||
3'-CUCUUGCU CU CC-5'
           G  G
```

**-18.17 kcal/mol**

Hedgehog, Mouse, Northern treeshrew, Pig, Rabbit, Rat, Squirrel

/

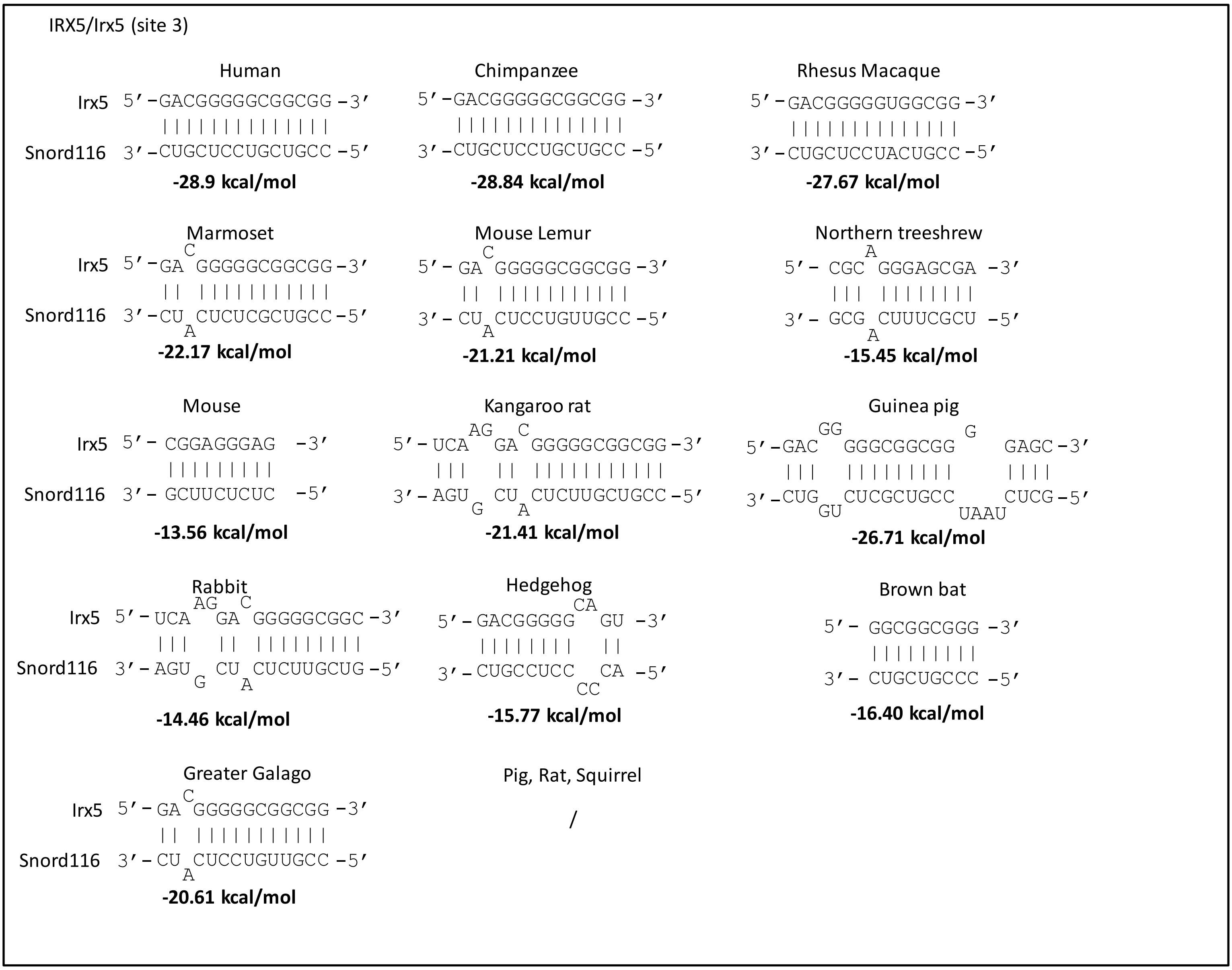

IRX5/Irx5 (site 3)
Human
Irx5 5′-GACGGGGGCGGCGG-3′
Snord116 3′-CUGCUCCUGCUGCC-5′
-28.9 kcal/mol
Chimpanzee
5′-GACGGGGGCGGCGG-3′
3′-CUGCUCCUGCUGCC-5′
-28.84 kcal/mol
Rhesus Macaque
5′-GACGGGGGUGGCGG-3′
3′-CUGCUCCUACUGCC-5′
-27.67 kcal/mol
Marmoset
Irx5 5′-GA C GGGGGCGGCGG-3′
Snord116 3′-CU A CUCUCGCUGCC-5′
-22.17 kcal/mol
Mouse Lemur
5′-GA C GGGGGCGGCGG-3′
3′-CU A CUCCUGUUGCC-5′
-21.21 kcal/mol
Northern treeshrew
5′-CGC A GGGAGCGA-3′
3′-GCG A CUUUCGCU-5′
-15.45 kcal/mol
Mouse
Irx5 5′-CGGAGGGAG-3′
Snord116 3′-GCUUCUCUC-5′
-13.56 kcal/mol
Kangaroo rat
5′-UCA AG GA C GGGGGCGGCGG-3′
3′-AGU G CU A CUCUUGCUGCC-5′
-21.41 kcal/mol
Guinea pig
5′-GAC GG GGGCGGCGG G GAGC-3′
3′-CUG GU CUCGCUGCC UAAU CUCG-5′
-26.71 kcal/mol
Rabbit
Irx5 5′-UCA AG GA C GGGGGCGGC-3′
Snord116 3′-AGU G CU A CUCUUGCUG-5′
-14.46 kcal/mol
Hedgehog
5′-GACGGGGG CA GU-3′
3′-CUGCCUCC CC CA-5′
-15.77 kcal/mol
Brown bat
5′-GGCGGCGGG-3′
3′-CUGCUGCCC-5′
-16.40 kcal/mol
Greater Galago
Irx5 5′-GA C GGGGGCGGCGG-3′
Snord116 3′-CU A CUCCUGUUGCC-5′
-20.61 kcal/mol
Pig, Rat, Squirrel
/


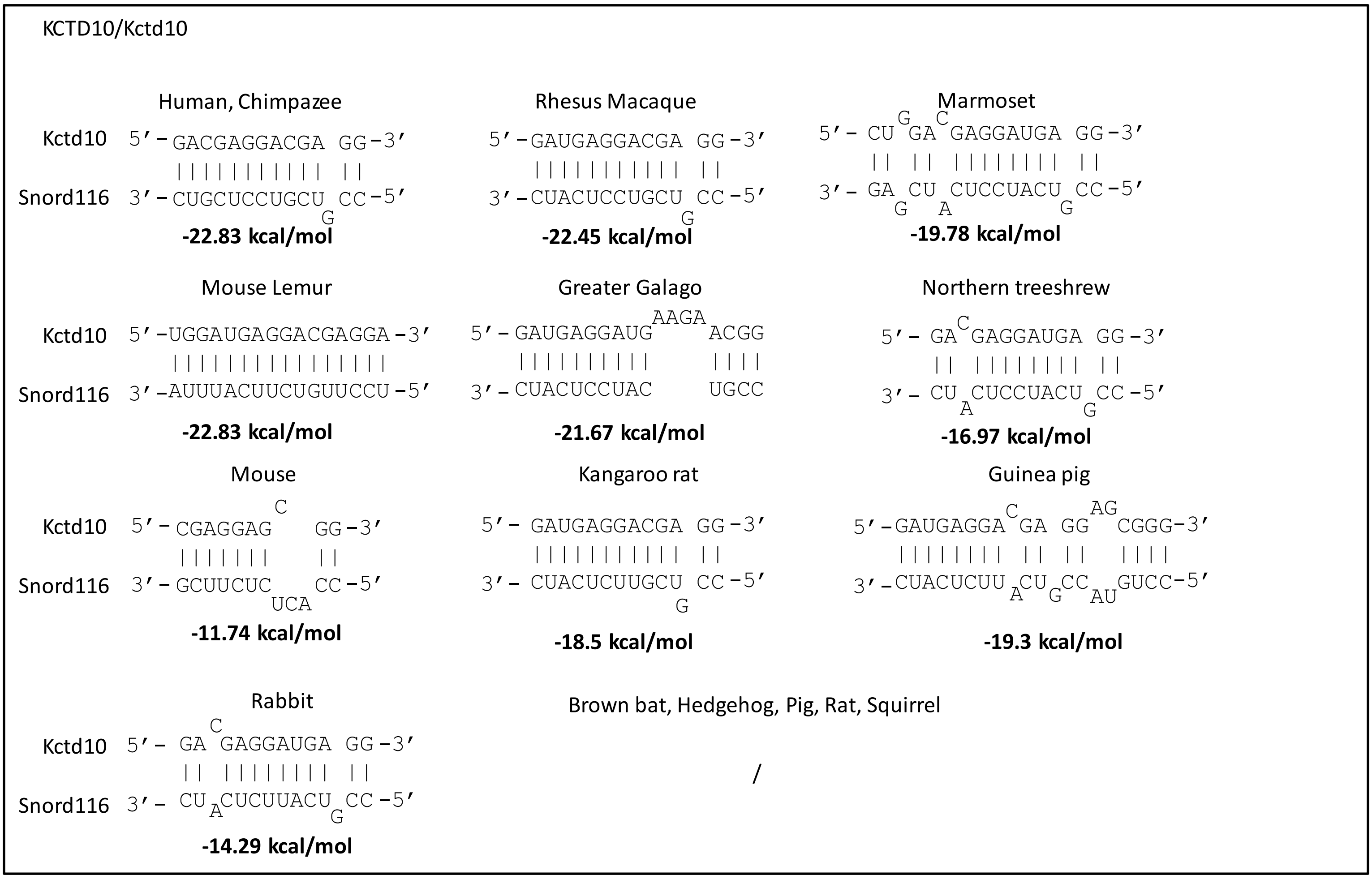

KCTD10/Kctd10
Human, Chimpazee
Kctd10 5′-GACGAGGACGA GG-3′
Snord116 3′-CUGCUCCUGCU G CC-5′
-22.83 kcal/mol
Rhesus Macaque
5′-GAUGAGGACGA GG-3′
3′-CUACUCCUGCU G CC-5′
-22.45 kcal/mol
Marmoset
5′-CU G GA C GAGGAUGA GG-3′
3′-GA G CU A CUCCUACU G CC-5′
-19.78 kcal/mol
Mouse Lemur
Kctd10 5′-UGGAUGAGGACGAGGA-3′
Snord116 3′-AUUUACUUCUGUUCCU-5′
-22.83 kcal/mol
Greater Galago
5′-GAUGAGGAUG AAGA ACGG
3′-CUACUCCUAC UGCC
-21.67 kcal/mol
Northern treeshrew
5′-GA C GAGGAUGA GG-3′
3′-CU A CUCCUACU G CC-5′
-16.97 kcal/mol
Mouse
Kctd10 5′-CGAGGAG C GG-3′
Snord116 3′-GCUUCUC UCA CC-5′
-11.74 kcal/mol
Kangaroo rat
5′-GAUGAGGACGA GG-3′
3′-CUACUCUUGCU G CC-5′
-18.5 kcal/mol
Guinea pig
5′-GAUGAGGA C GA GG AG CGGG-3′
3′-CUACUCUU A CU G CC AU GUCC-5′
-19.3 kcal/mol
Rabbit
Kctd10 5′-GA C GAGGAUGA GG-3′
Snord116 3′-CU A CUCUUACU G CC-5′
-14.29 kcal/mol
Brown bat, Hedgehog, Pig, Rat, Squirrel
/

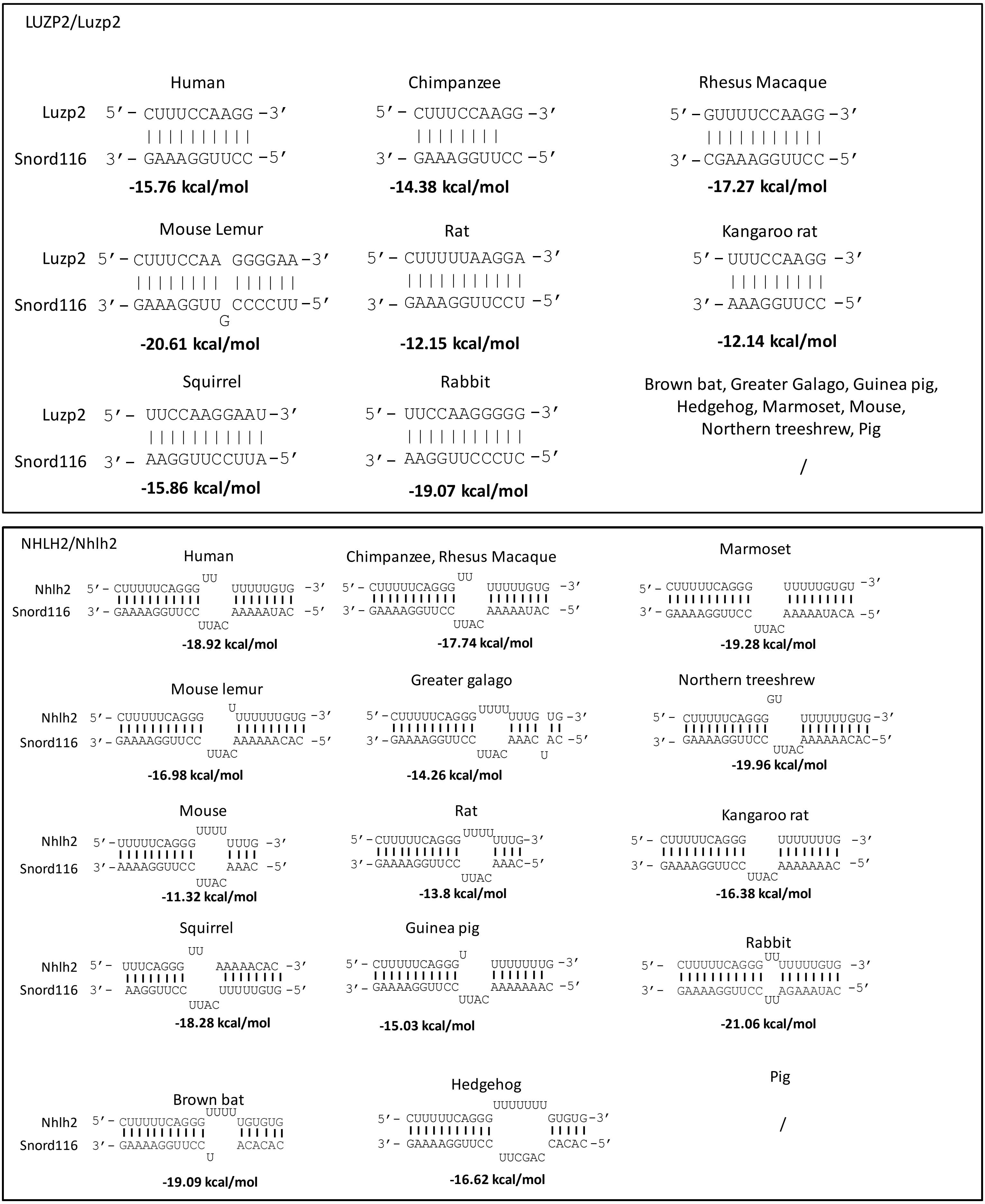
LUZP2/Luzp2
Human
Luzp2 5′ – CUUUCCAAGG –3′
Snord116 3′ – GAAAGGUUCC –5′
-15.76 kcal/mol
Chimpanzee
5′ – CUUUCCAAGG –3′
3′ – GAAAGGUUCC –5′
-14.38 kcal/mol
Rhesus Macaque
5′ – GUUUUCCAAGG –3′
3′ – CGAAAGGUUCC –5′
-17.27 kcal/mol
Mouse Lemur
Luzp2 5′ – CUUUCCAA GGGGAA –3′
Snord116 3′ – GAAAGGUU CCCCUU –5′
G
-20.61 kcal/mol
Rat
5′ – CUUUUUAAGGA –3′
3′ – GAAAGGUUCCU –5′
-12.15 kcal/mol
Kangaroo rat
5′ – UUUCCAAGG –3′
3′ – AAAGGUUCC –5′
-12.14 kcal/mol
Squirrel
Luzp2 5′ – UUCCAAGGAAU –3′
Snord116 3′ – AAGGUUCCUUA –5′
-15.86 kcal/mol
Rabbit
5′ – UUCCAAGGGGG –3′
3′ – AAGGUUCCCUC –5′
-19.07 kcal/mol
Brown bat, Greater Galago, Guinea pig, Hedgehog, Marmoset, Mouse, Northern treeshrew, Pig
/
NHLH2/Nhlh2
Human
UU
Nhlh2 5′ – CUUUUUCAGGG UUUUUGUG –3′
Snord116 3′ – GAAAAGGUUCC AAAAAUAC –5′
UUAC
-18.92 kcal/mol
Chimpanzee, Rhesus Macaque
UU
5′ – CUUUUUCAGGG UUUUUGUG –3′
3′ – GAAAAGGUUCC AAAAAUAC –5′
UUAC
-17.74 kcal/mol
Marmoset
5′ – CUUUUUCAGGG UUUUUGUGU –3′
3′ – GAAAAGGUUCC AAAAAUACA –5′
UUAC
-19.28 kcal/mol
Mouse lemur
U
Nhlh2 5′ – CUUUUUCAGGG UUUUUUGUG –3′
Snord116 3′ – GAAAAGGUUCC AAAAAACAC –5′
UUAC
-16.98 kcal/mol
Greater galago
UUUU
5′ – CUUUUUCAGGG UUUG UG –3′
3′ – GAAAAGGUUCC AAAC AC –5′
UUAC U
-14.26 kcal/mol
Northern treeshrew
GU
5′ – CUUUUUCAGGG UUUUUUGUG –3′
3′ – GAAAAGGUUCC AAAAAACAC –5′
UUAC
-19.96 kcal/mol
Mouse
UUUU
Nhlh2 5′ – UUUUUCAGGG UUUG –3′
Snord116 3′ – AAAAGGUUCC AAAC –5′
UUAC
-11.32 kcal/mol
Rat
UUUU
5′ – CUUUUUCAGGG UUUG –3′
3′ – GAAAAGGUUCC AAAC –5′
UUAC
-13.8 kcal/mol
Kangaroo rat
5′ – CUUUUUCAGGG UUUUUUUG –3′
3′ – GAAAAGGUUCC AAAAAAAC –5′
UUAC
-16.38 kcal/mol
Squirrel
UU
Nhlh2 5′ – UUUCAGGG AAAAACAC –3′
Snord116 3′ – AAGGUUCC UUUUUGUG –5′
UUAC
-18.28 kcal/mol
Guinea pig
U
5′ – CUUUUUCAGGG UUUUUUUG –3′
3′ – GAAAAGGUUCC AAAAAAAC –5′
UUAC
-15.03 kcal/mol
Rabbit
UU
5′ – CUUUUUCAGGG UUUUUGUG –3′
3′ – GAAAAGGUUCC AGAAAUAC –5′
UU
-21.06 kcal/mol
Brown bat
UUUU
Nhlh2 5′ – CUUUUUCAGGG UGUGUG
Snord116 3′ – GAAAAGGUUCC ACACAC
U
-19.09 kcal/mol
Hedgehog
UUUUUUU
5′ – CUUUUUCAGGG GUGUG –3′
3′ – GAAAAGGUUCC CACAC –5′
UUCGAC
-16.62 kcal/mol
Pig
/

NLGN3/Nlgn3

**Human,Chimpanzee**

```
Nlgn3     5'- UUUCCAAGGAAUGC -3'
              ||||||||||||||
Snord116  3'- AAAGGUUCCUUACG -5'
```

**-22.62 kcal/mol**

**Rhesus Macaque**

```
5'- UUUCCAAGGAAUGC -3'
    ||||||||||||||
3'- AAAGGUUCCUUACG -5'
```

**-22.93 kcal/mol**

**Marmoset**

```
5'- UUUCCAAGGAAUGC-3'
    ||||||||||||||
3'- AAAGGUUCCUUACG-5'
```

**-23.24 kcal/mol**

**Mouse Lemur**

```
Nlgn3     5'- UUCCAAGGAAUGC  -3'
              |||||||||||||
Snord116  3'- AAGGUUCCUUACG  -5'
```

**-22.41 kcal/mol**

**Greater Galago**

```
5'- UUUCCAAGGAAUGC -3'
    ||||||||||||||
3'- AAAGGUUCCUUACG -5'
```

**-23.41 kcal/mol**

**Northern treeshrew**

```
5'- UUUCCAAGGAAUGU -3'
    ||||||||||||||
3'- AAAGGUUCCUUACA -5'
```

**-21.38 kcal/mol**

**Mouse**

```
Nlgn3     5'- UUUCCAAGGAAUG  -3'
              |||||||||||||
Snord116  3'- AAAGGUUCCUUAC  -5'
```

**-19.70 kcal/mol**

**Rat, Kangaroo Rat**

```
5'- UUUCCAAGGAAUG  -3'
    |||||||||||||
3'- AAAGGUUCCUUAC  -5'
```

**-19.68 kcal/mol**

**Squirrel**

```
5'- UUUCCAAGGAAUG  -3'
    |||||||||||||
3'- AAAGGUUCCUUAC  -5'
```

**-19.54 kcal/mol**

**Guinea pig**

```
Nlgn3     5'- UUUCCAAGGAAUGC -3'
              ||||||||||||||
Snord116  3'- AAAGGUUCCUUACG -5'
```

**-23.42 kcal/mol**

**Rabbit**

```
5'- UUUCCAAGGAAUGU -3'
    ||||||||||||||
3'- AAAGGUUCCUUACA -5'
```

**-19.60 kcal/mol**

**Pig**

```
5'- UUUCCAAGGAAUGC -3'
    ||||||||||||||
3'- AAAGGUUCCUUACG -5'
```

**-23.13 kcal/mol**

**Brown bat**

```
Nlgn3     5'- UUUCCAAGGAAUGC -3'
              ||||||||||||||
Snord116  3'- AAAGGUUCCUUACG -5'
```

**-22.93 kcal/mol**

**Hedgehog**

```
5'- UUUCCAAGGAAUGC -3'
    ||||||||||||||
3'- AAAGGUUCCUUACG -5'
```

**-22.45 kcal/mol**

RET/Ret

Human
```
Ret       5'-AGGUGGGGAUGGUGGU -3'
             ||||||||||||||||
Snord116  3'-UCUACUCCUACUACCA -5'
```
**-26.12 kcal/mol**

Chimpanzee
```
5'-GGUGGGGAUGGUGGU -3'
   |||||||||||||||
3'-CUGCUCCUACUGCCA -5'
```
**-21.3 kcal/mol**

Rhesus macaque
```
5'- GGUGAUGGUG -3'
    ||||||||||
3'- CUACUGCCAC -5'
```
**-15.65 kcal/mol**

Marmoset
```
                C  U          G
Ret       5'-GAGG AG GGUGGUGGU AUGGUGGUU   -3'
             |||| || ||||||||| |||||||||
Snord116  3'-CUUC UC CUACUGUCA UAUCACCAA   -5'
```
**-24.8 kcal/mol**

Mouse Lemur
```
          G
5'- GAGGA GA GGUGUGGAG -3'
    ||||| || |||||||||
3'- CUCUU CU CCAUAUCUC -5'
         G  G
```
**-23.28 kcal/mol**

Greater Galago
```
                         U
Ret       5'- UGGUG GUGA GGUGGUGGUGA -3'
              ||||| |||| |||||||||||
Snord116  3'- ACUAC CACU CUACCAUCACU -5'
                   U
```
**-28.32 kcal/mol**

Rabbit
```
5'- GUGAUGGGAA -3'
    ||||||||||
3'- CACUACUCUU -5'
```
**-13.14 kcal/mol**

Brown bat
```
      GA
5'-AGU  UGCUGAUG-3'
   |||  ||||||||
3'-UCA  AUGACUAC-5'
```
**-10.03 kcal/mol**

Kangaroo rat
```
                   GU
Ret       5'- AUGGUG  AAUGGUGGU -3'
              ||||||  |||||||||
Snord116  3'- UGCUAC  UUGCUGCCA -5'
                    UC
```
**-14.26 kcal/mol**

Guinea pig, Hedgehog, Kangaroo rat, Mouse, Northern treeshrew, Pig, Rat, Squirrel

/

RSBN1L/Rsbn1l

Human, Chimpanzee, Rhesus Macaque, Marmoset, Mouse Lemur, Greater Galago, Northern treeshrew, Mouse, Kangaroo Rat, Rabbit
```
Rsbn1l    5'-  UUCCAAGGAAUGUU -3'
               ||||||||||||||
Snord116  3'-  AAGGUUCCUUACAA -5'
```
**-20.6 kcal/mol**

Rat
```
          A
5'-  UUCCA GGAAUGUU -3'
     ||||| ||||||||
3'-  AAGGU CCUUACAA -5'
          U
```
**-16.39 kcal/mol**

Squirrel
```
Rsbn1l    5'-UUCCAAGGAAUGU -3'
             |||||||||||||
Snord116  3'-AAGGUUCCUUACA -5'
```
**-20.1 kcal/mol**

Guinea Pig
```
5'-UUCCAAGGAAUGU -3'
   |||||||||||||
3'-AAGGUUCCUUACA -5'
```
**-20.2 kcal/mol**

Pig
```
5'-UUCCAAGGAAUGUU -3'
   ||||||||||||||
3'-AAGGUUCCUUACGA -5'
```
**-20.5 kcal/mol**

Brown bat
```
Rsbn1l    5'-UUCCAAGGAAUGUU-3'
             ||||||||||||||
Snord116  3'-AAGGUUCCUUACAA-5'
```
**-20.47 kcal/mol**

Hedgehog
```
5'-UUCCAAGGAAUGUU -3'
   ||||||||||||||
3'-AAGGUUCCUUACGA -5'
```
**-20.58 kcal/mol**

SNX33/Snx33 (site 1)

Human
```
                G
Snx33     5'-UUUGA GAGGAUGAUG -3'
             ||||| ||||||||||
Snord116  3'-AAACU CUCCUGCUGC -5'
                  G
```
**-23.47 kcal/mol**

Chimpanzee
```
          G
5'-UUCGA GAGGAUGAUG -3'
   ||||| ||||||||||
3'-AAGCU CUCCUACUGC -5'
        G
```
**-25.67 kcal/mol**

Rhesus Macaque
```
           UUG        U
5'- UGAUGA    GGAUGAC  GG -3'
    ||||||    ||||||| ||
3'- ACUACU    CCUGCUG CC -5'
```
**-19.2 kcal/mol**

Marmoset
```
Snx33     5'-UUCGAGGAGGAUGAUG -3'
             ||||||||||||||||
Snord116  3'-AAGCUCCUCCUACUGC -5'
```
**-29.54 kcal/mol**

Mouse lemur
```
          GG  A
5'-CUUUGA  AGG GGAUGAUG-3'
   ||||||  ||| ||||||||
3'-GAAACU  UCC CCUACUGC-5'
```
**-21.79 kcal/mol**

Greater Galago
```
           CUG        U
5'- UGAUGA    GGAUGAC  GG -3'
    ||||||    ||||||| ||
3'- ACUACU    CCUACUG CC -5'
```
**-19.6 kcal/mol**

Kangaroo rat
```
                   U
Snx33     5'- GGAUGAC GG -3'
              ||||||| ||
Snord116  3'- CUUGCUG CC -5'
```
**-9.62 kcal/mol**

Squirrel
```
           UU          AC
5'- AUGAUGA  GGGAUG      UGGG -3'
    |||||||  ||||||      ||||
3'- UGCUACU  CUCUGC      ACCC -5'
           CG      CAUC
```
**-18.98 kcal/mol**

Rabbit
```
5'- GAGGACGACG -3'
    ||||||||||
3'- CUCUUGCUGC -5'
```
**-17.57 kcal/mol**

Brown bat
```
                  AUC
Snx33     5'- UGAUG   GGGAUGAC  -3'
              |||||   ||||||||
Snord116  3'- ACUAC   CCUUACUG  -5'
```
**-15.18 kcal/mol**

Guinea pig, Hedgehog, Mouse, Northern treeshrew, Pig, Rat

/

SNX33/Snx33 (site 2)

Human
```
                  CA
Snx33     5'-GACGAGG  GACG -3'
             |||||||  ||||
Snord116  3'-CUGCUCC  CUGC -5'
                    UG
```
**-18.5 kcal/mol**

Rhesus Macaque
```
           CA
5'- GACGAGG  GAUGGCAU-3'
    |||||||  ||||||||
3'- CUGCUCC  CUGCCGUA-5'
           UA
```
**-26.38 kcal/mol**

Marmoset
```
           CA
5'- GACGAGG  GAUGGCA -3'
    |||||||  |||||||
3'- CUGCUCC  CUGCCGU -5'
           UA
```
**-25.36 kcal/mol**

Guinea pig
```
Snx33     5'- AGGAUGA GG  -3'
              ||||||| ||
Snord116  3'- UCUUACU CC  -5'
                     G
```
**-8.6 kcal/mol**

Hedgehog
```
           U
5'- AGGCGGA GG-3'
    ||||||| ||
3'- UCUGCCU CC-5'
           C
```
**-12.57 kcal/mol**

Brown bat, Chimpanzee, Greater Galago, Kangaroo rat, Mouse lemur, Mouse, Northern treeshrew, Pig, Rabbit, Rat, Squirrel

/

SYK/Syk (site 1)

Human, Chimpanzee, Rhesus macaque

```
Syk       5'- GUGGCACAGAA -3'
              |||||||||||
Snord116  3'- CACCGUGUCUU -5'
```

**-19.38 kcal/mol**

Brown bat, Greater Galago, Guinea pig, Hedgehog, Kangaroo rat, Marmoset, Mouse lemur, Mouse, Northern treeshrew, Pig, Rabbit, Rat, Squirrel

/

SYK/Syk (site 2)

Human, Chimpanzee, Rhesus Macaque

```
Syk       5'- UUUCCAAGGAAUGUA -3'
              |||||||||||||||
Snord116  3'- AAAGGUUCCUUACAU -5'
```

**-21.87 kcal/mol**

Marmoset

```
        G
5'- UUC AAGGAAUG -3'
    |||  ||||||||
3'- AAG UUCCUUAC -5'
       G
```

**-15.5 kcal/mol**

Mouse Lemur

```
          A
5'- UUCCAA GAAUGCA -3'
    ||||||  |||||||
3'- AAGGUU CUUACGU -5'
          C
```

**-16.36 kcal/mol**

Northern treeshrew

```
Syk       5'- UUUCCAAGGAAUA -3'
              |||||||||||||
Snord116  3'- AAAGGUUCCUUAU -5'
```

**-17.6 kcal/mol**

Brown bat, Greater Galago, Guinea pig, Hedgehog, Kangaroo rat, Mouse, Pig, Rabbit, Rat, Squirrel

/

ZIC2/Zic2

Human

```
                        AU
Zic2      5'- AAGUGGUUUG  CAUU -3'
              ||||||||||  ||||
Snord116  3'- UUCACCAAAC  GUGA -5'
                        GA
```

**-17.02 kcal/mol**

Squirrel

```
              AU
5'- AAGUGGUUUG  CAUU -3'
    ||||||||||  ||||
3'- UUCACCAAAC  GUGA -5'
              CA
```

**-15.9 kcal/mol**

Rabbit

```
              AU
5'- AAGUGGUUUG  CAUU -3'
    ||||||||||  ||||
3'- UUCACCAAAC  GUGA -5'
              GA
```

**-15.9 kcal/mol**

Brown bat, Chimpanzee, Greater Galago, Guinea pig, Hedgehog, Kangaroo rat, Marmoset, Mouse lemur, Mouse, Northern treeshrew, Rat, Rhesus macaque, Pig

/

Supplementary Figure 2

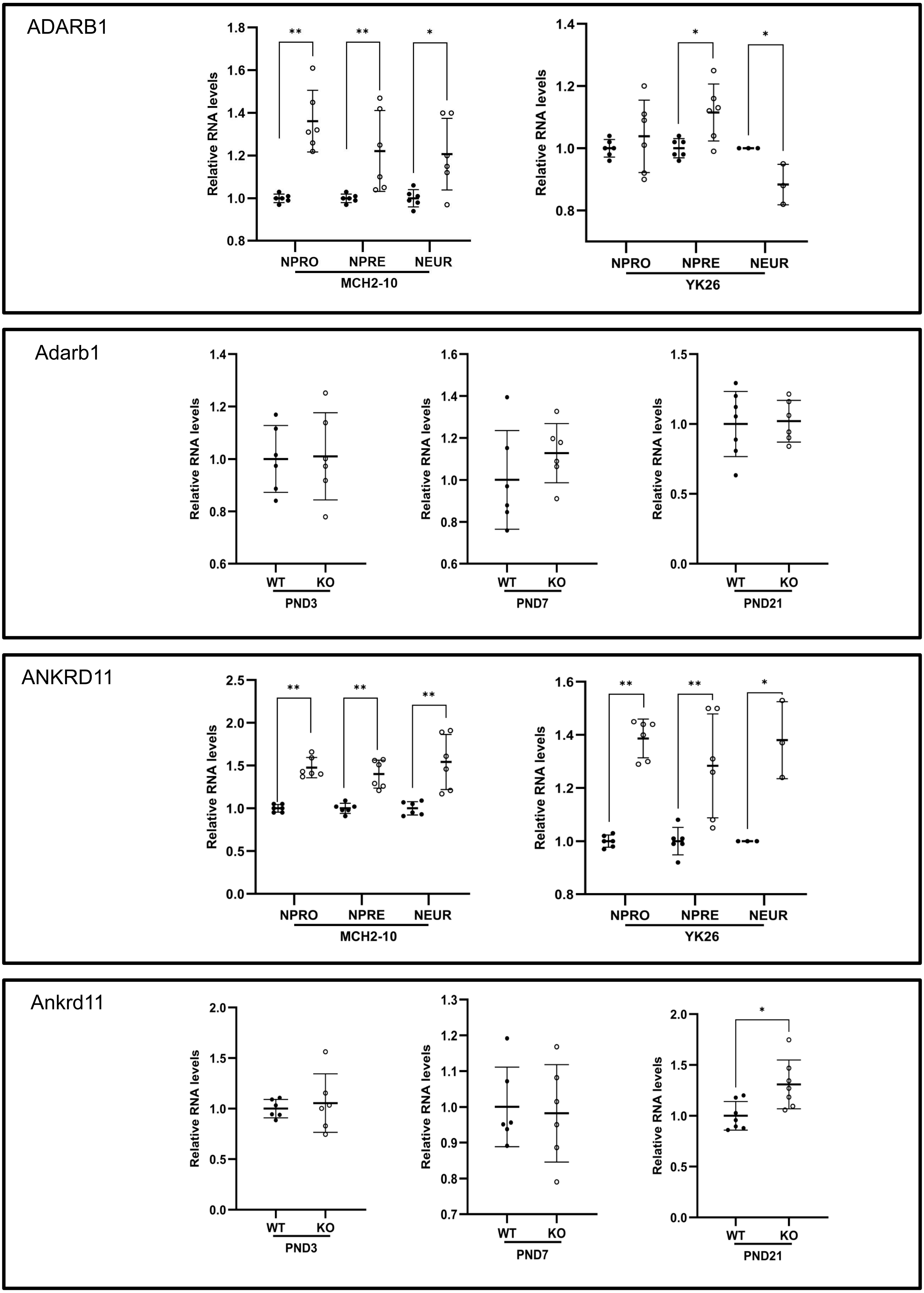

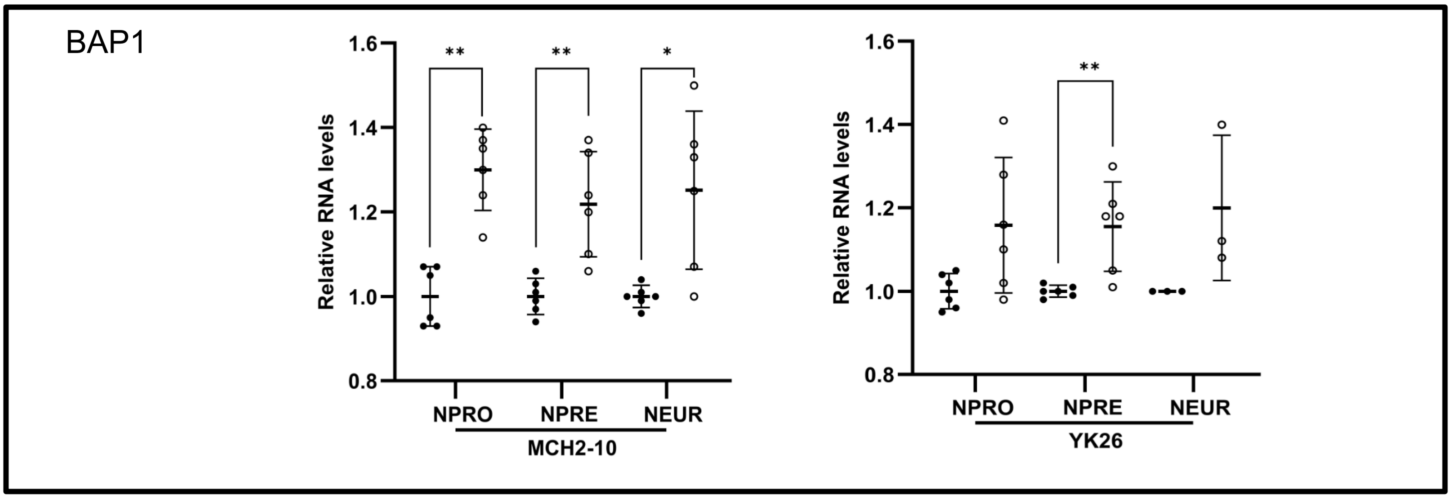
BAP1
Relative RNA levels
**
**
*
NPRO
NPRE
NEUR
MCH2-10
YK26

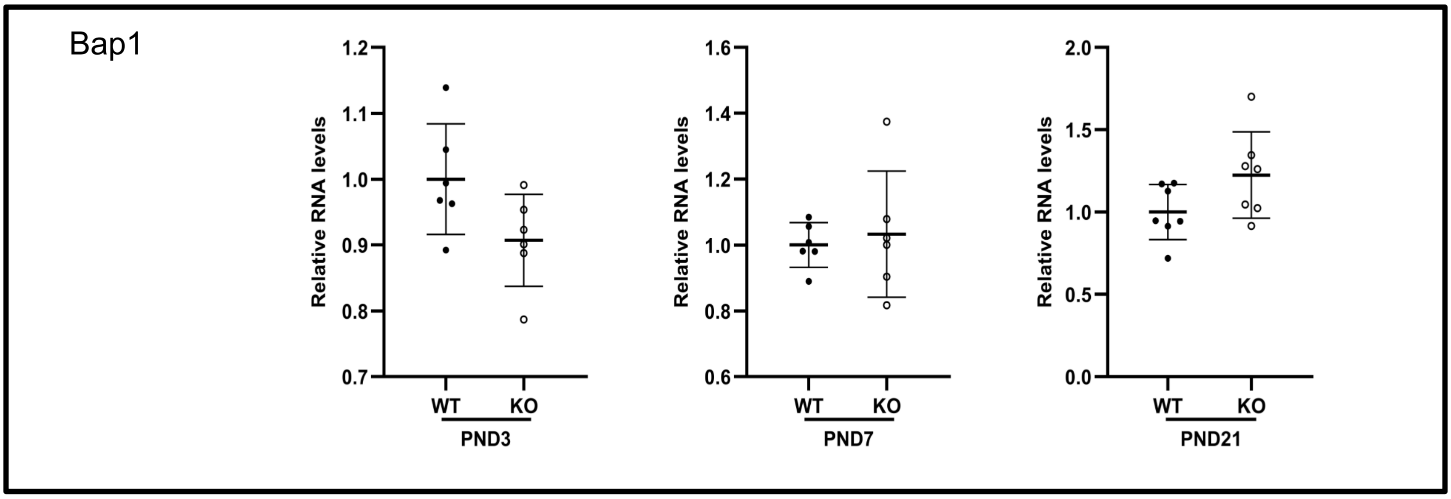
Bap1
Relative RNA levels
WT
KO
PND3
PND7
PND21

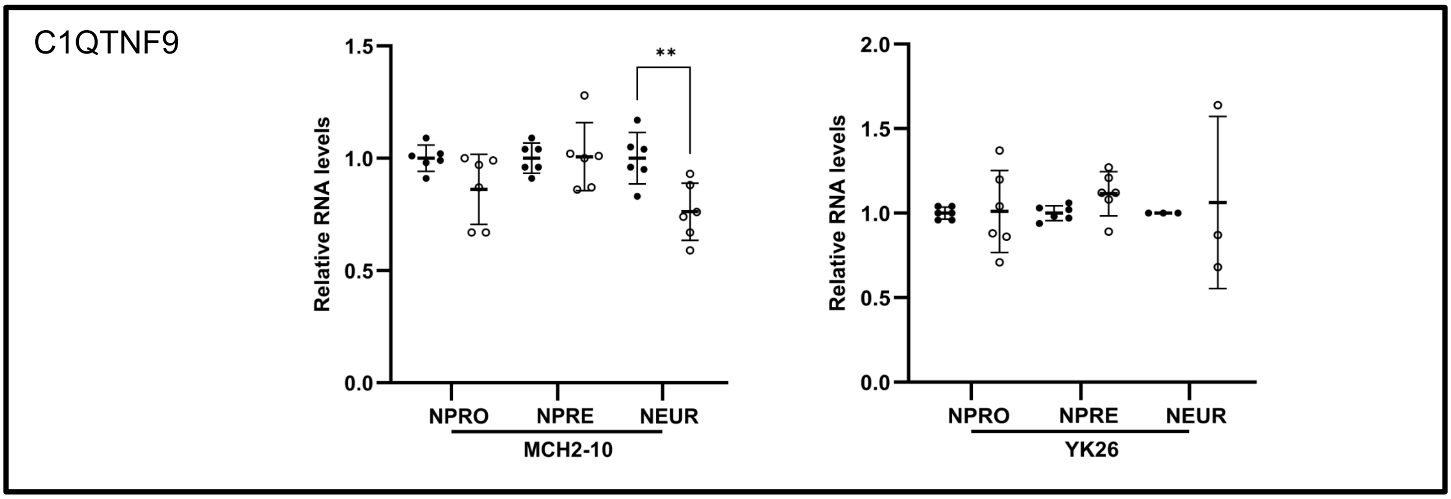
C1QTNF9
Relative RNA levels
**
NPRO
NPRE
NEUR
MCH2-10
YK26

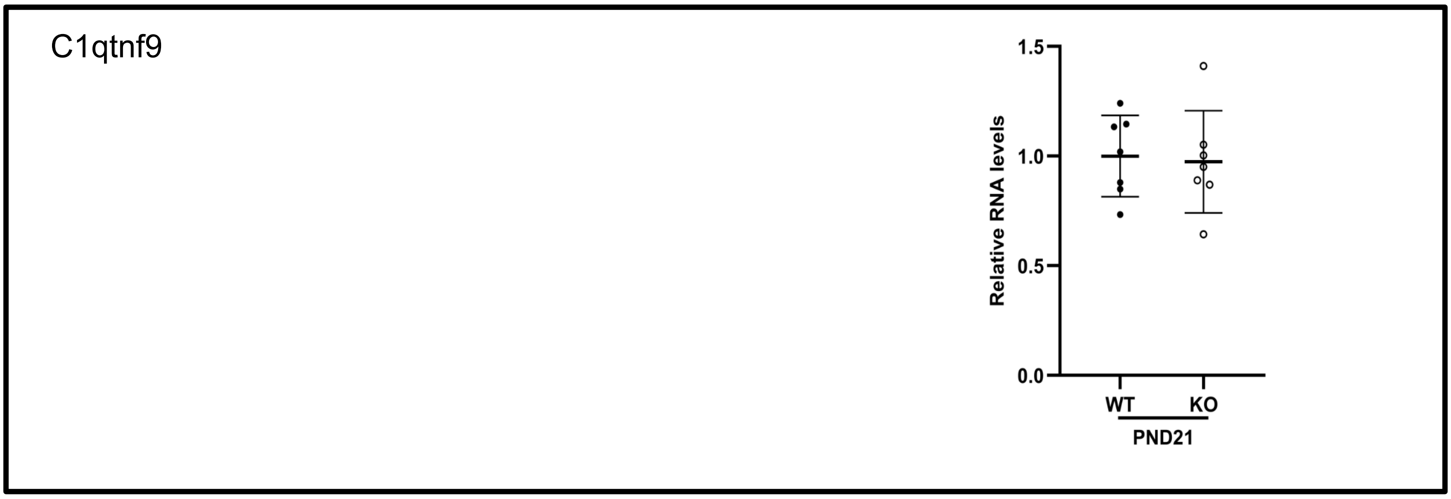
C1qtnf9
Relative RNA levels
WT
KO
PND21

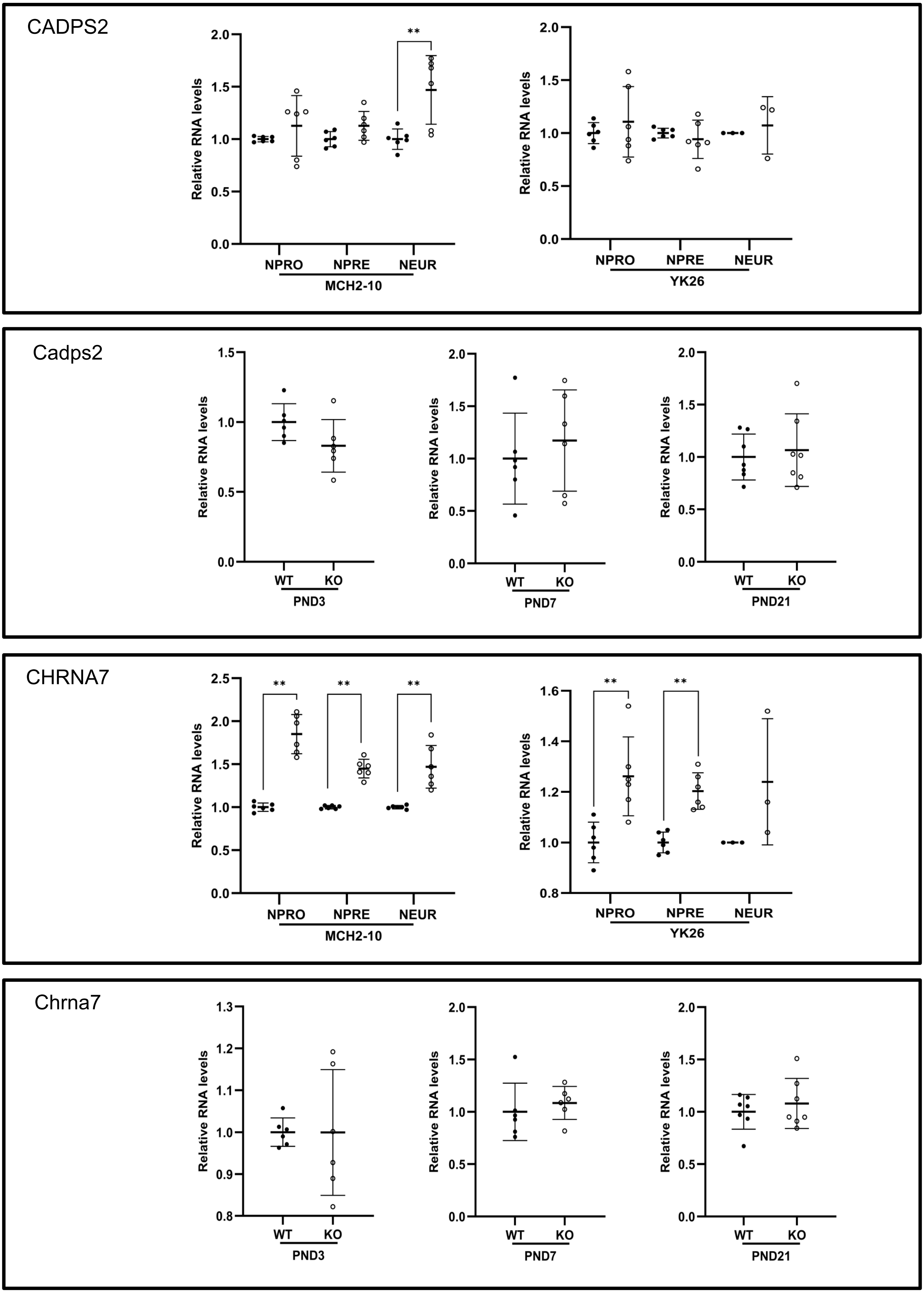
CADPS2
Relative RNA levels
2.0
1.5
1.0
0.5
0.0
**
NPRO
NPRE
NEUR
MCH2-10
YK26
Cadps2
WT
KO
PND3
PND7
PND21
CHRNA7
2.5
1.6
1.4
1.2
0.8
Chrna7
1.3
1.1
0.9

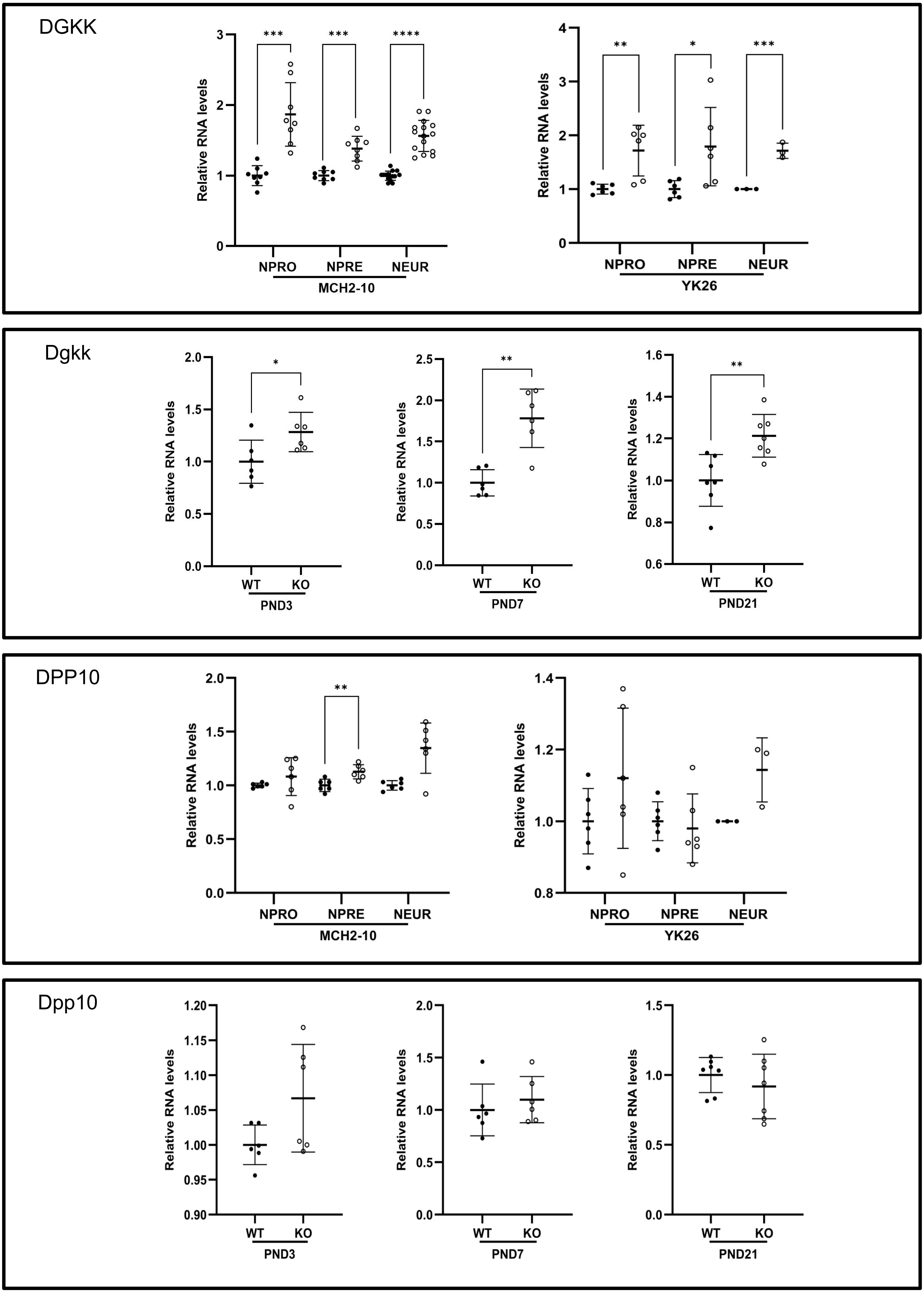
DGKK
Relative RNA levels
NPRO
NPRE
NEUR
MCH2-10
YK26
***
****
**
*
Dgkk
WT
KO
PND3
PND7
PND21
DPP10
Dpp10

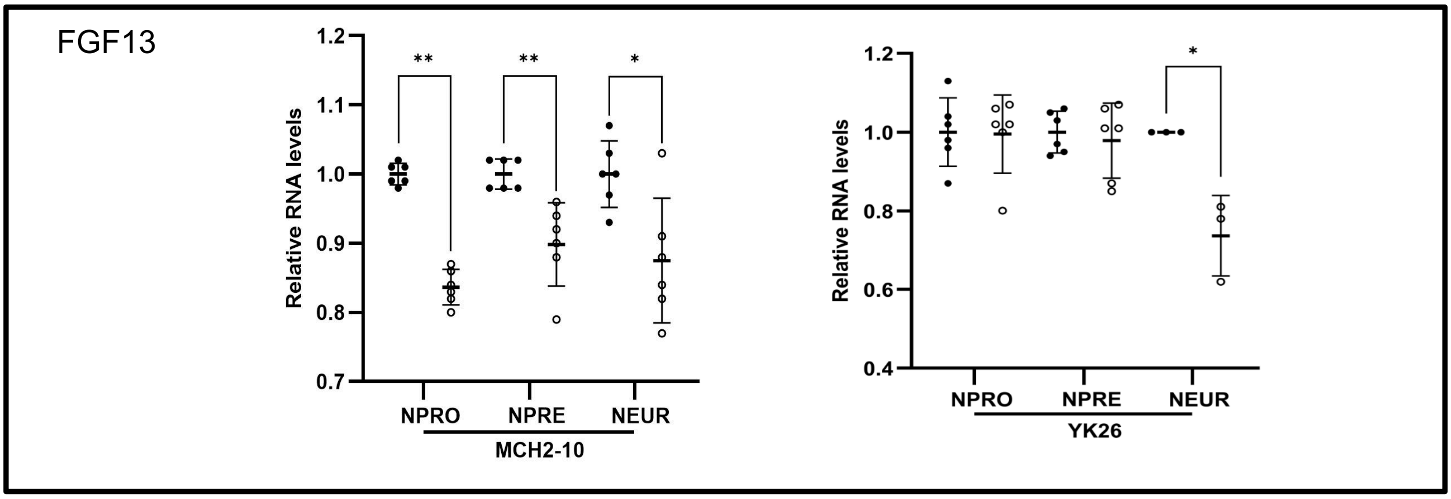
FGF13
Relative RNA levels
NPRO
NPRE
NEUR
MCH2-10
YK26
**
*

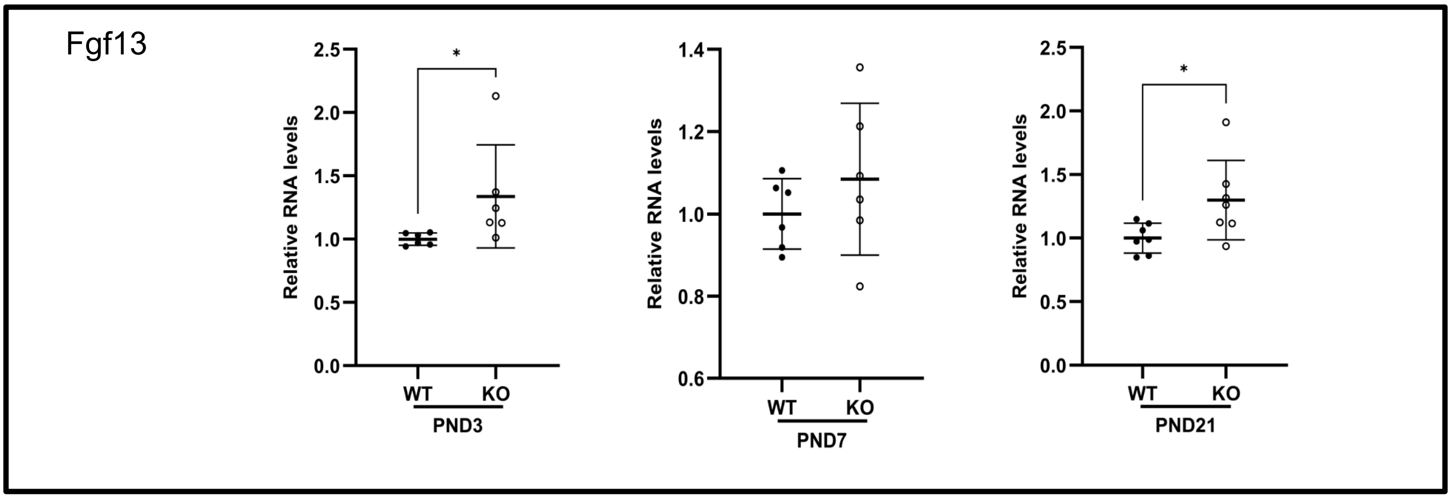
Fgf13
Relative RNA levels
WT
KO
PND3
PND7
PND21
*

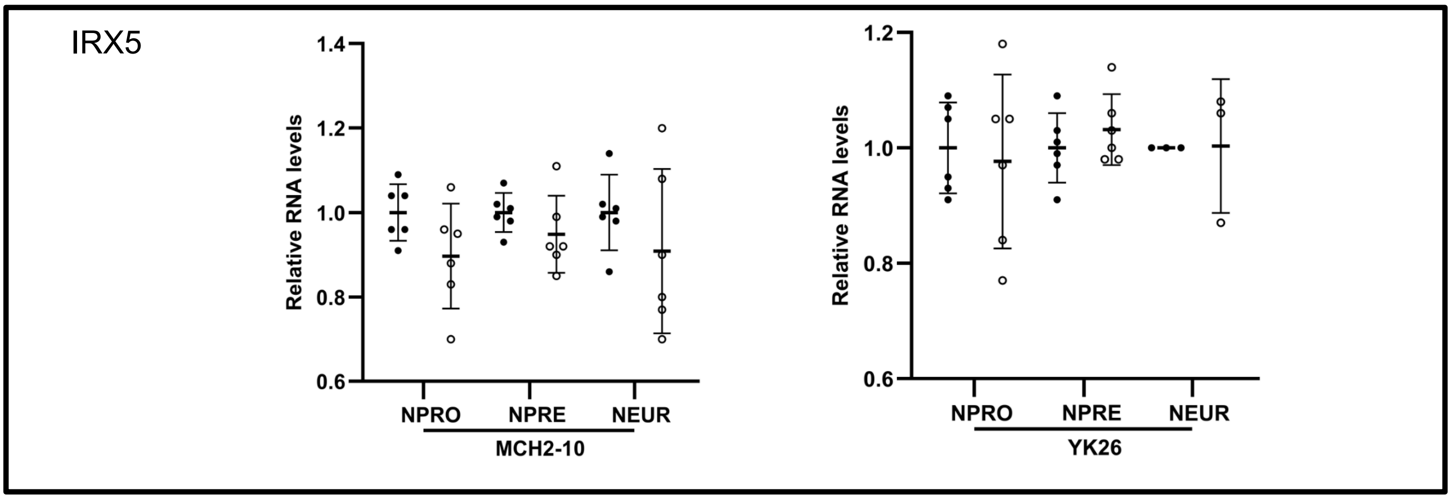
IRX5
Relative RNA levels
NPRO
NPRE
NEUR
MCH2-10
YK26

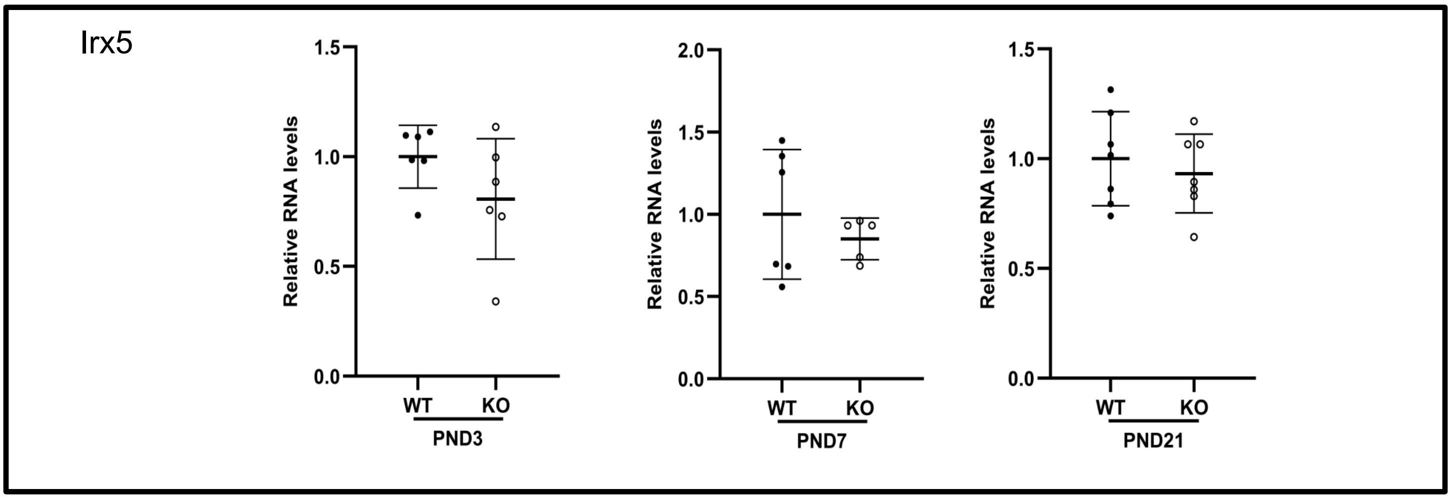
Irx5
Relative RNA levels
WT
KO
PND3
PND7
PND21

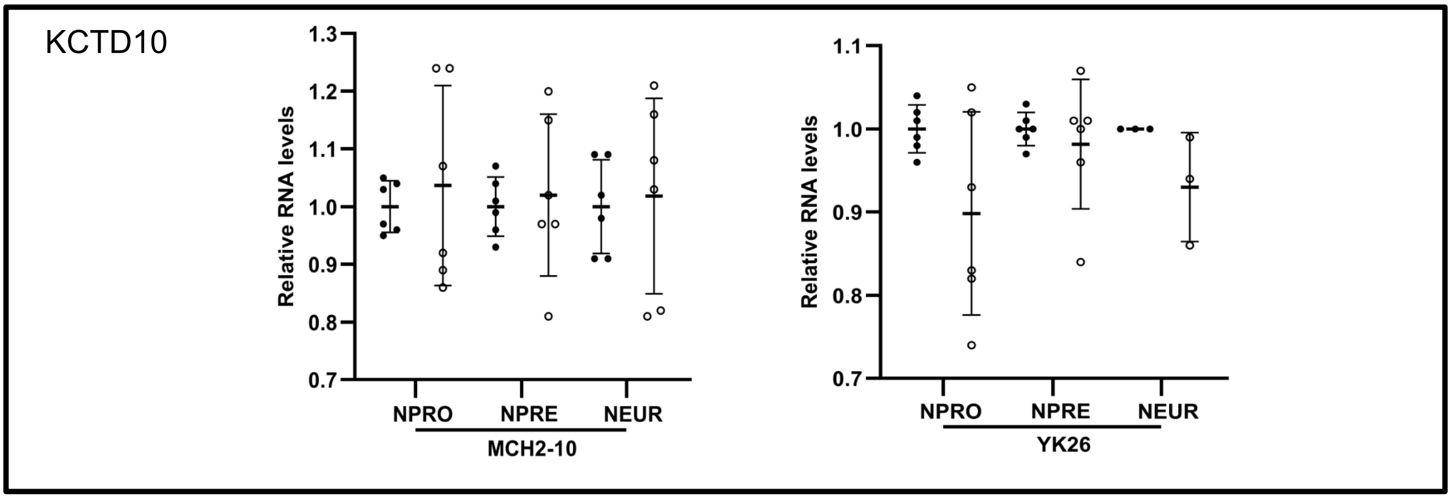
KCTD10
Relative RNA levels
1.3
1.2
1.1
1.0
0.9
0.8
0.7
NPRO
NPRE
NEUR
MCH2-10
1.1
1.0
0.9
0.8
0.7
YK26

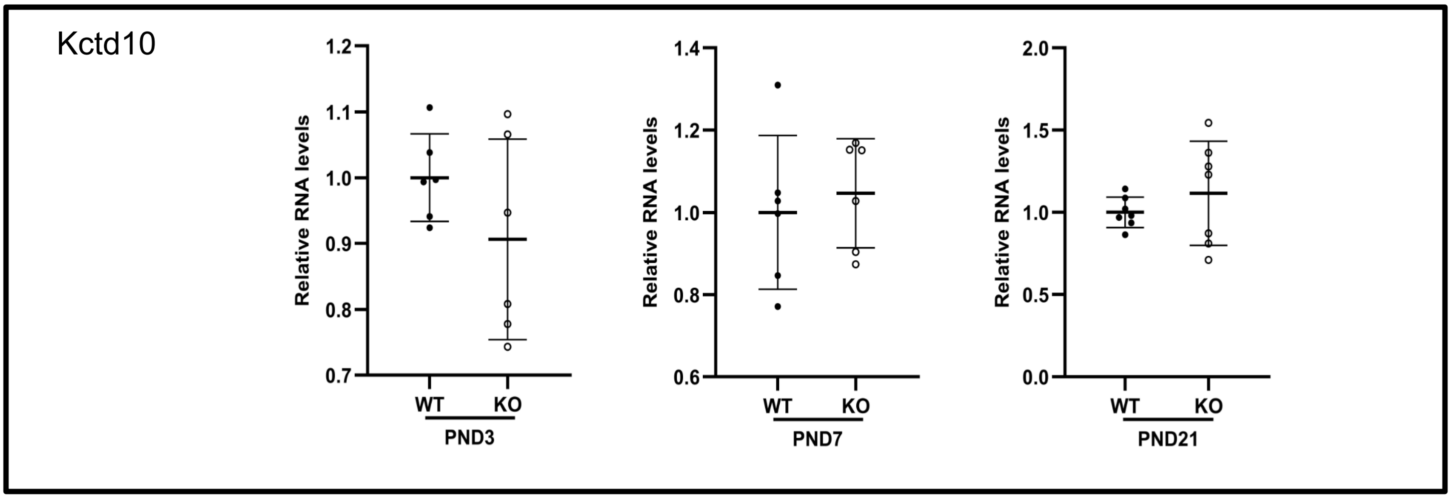
Kctd10
Relative RNA levels
1.2
1.1
1.0
0.9
0.8
0.7
WT
KO
PND3
1.4
1.2
1.0
0.8
0.6
PND7
2.0
1.5
1.0
0.5
0.0
PND21

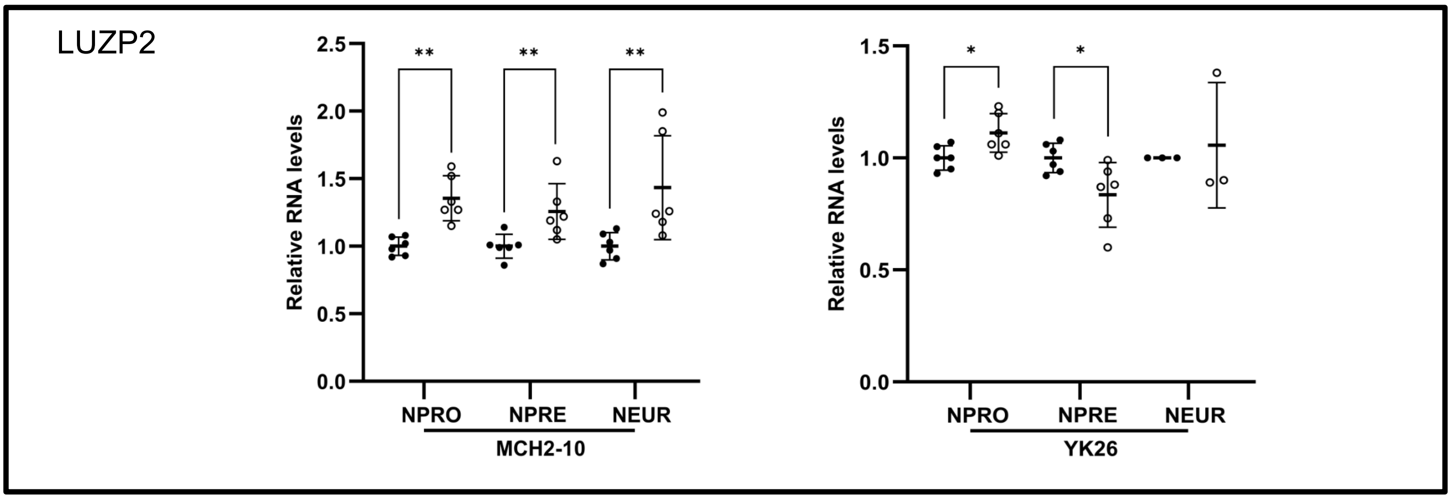
LUZP2
Relative RNA levels
2.5
2.0
1.5
1.0
0.5
0.0
**
NPRO
NPRE
NEUR
MCH2-10
1.5
1.0
0.5
0.0
*
YK26

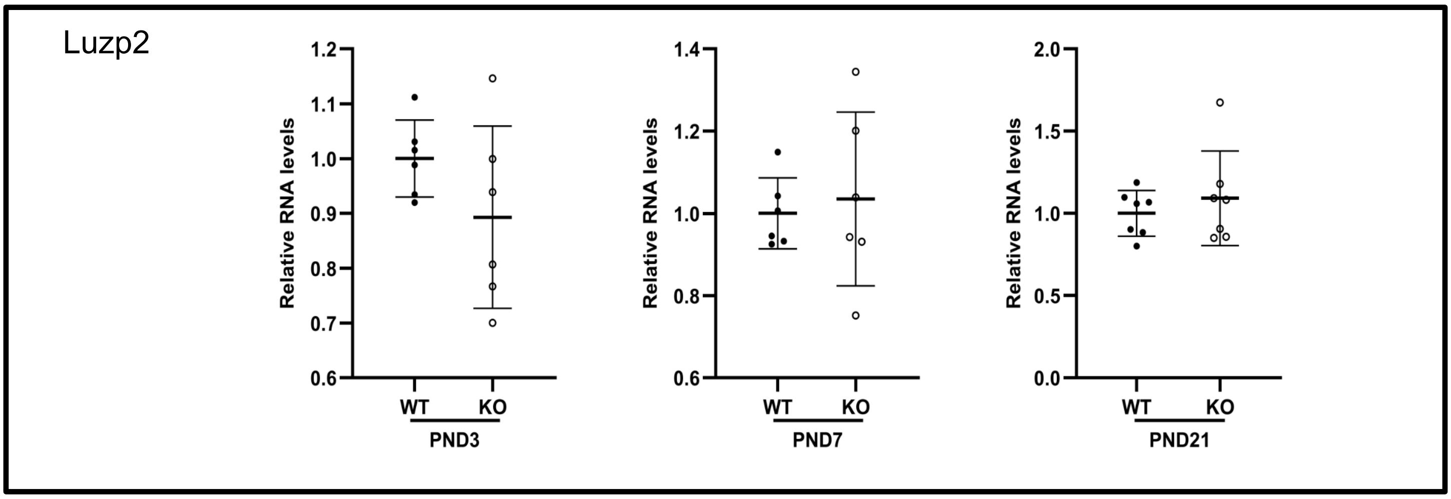
Luzp2
Relative RNA levels
1.2
1.1
1.0
0.9
0.8
0.7
0.6
WT
KO
PND3
1.4
1.2
1.0
0.8
0.6
PND7
2.0
1.5
1.0
0.5
0.0
PND21

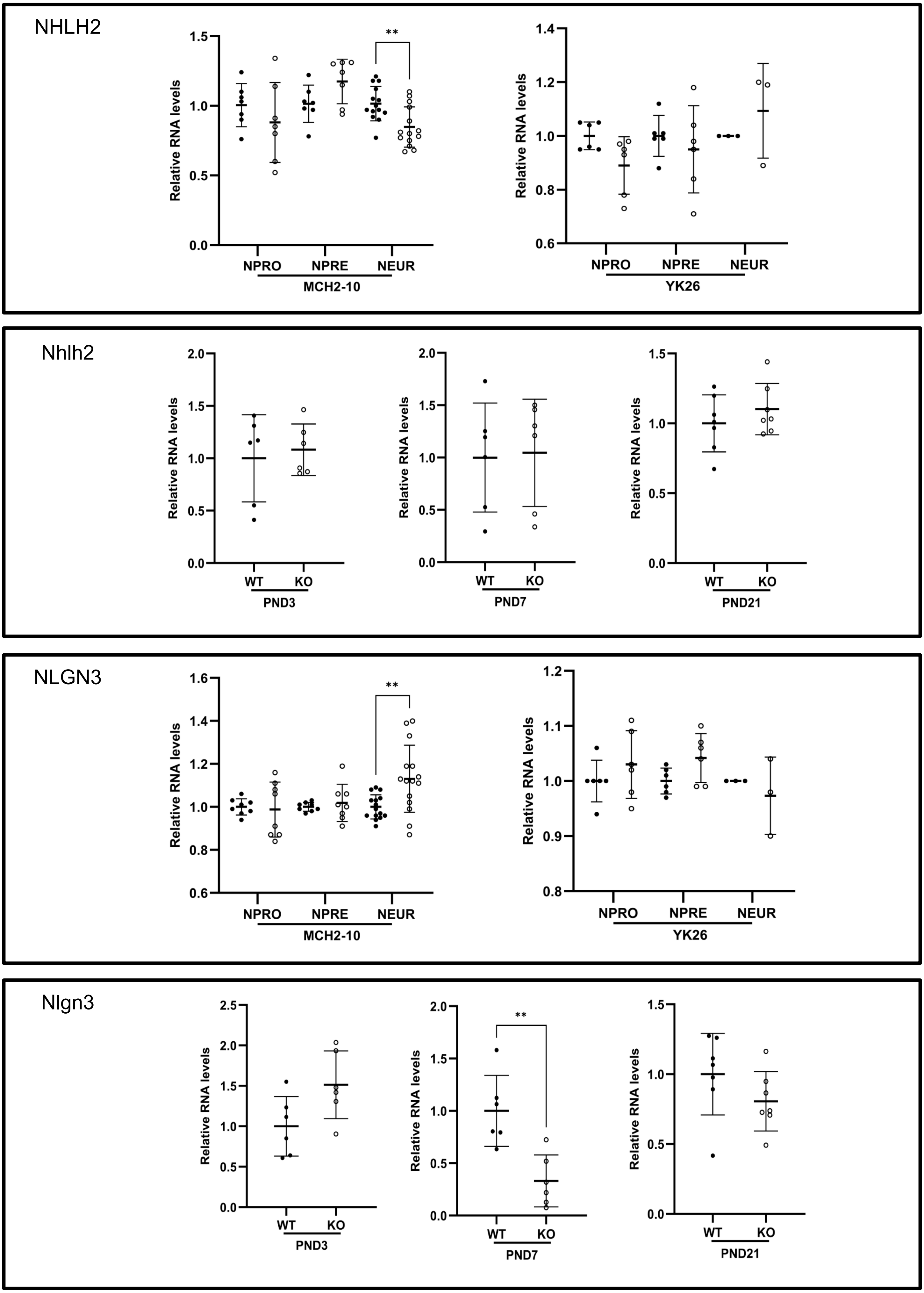
NHLH2
Relative RNA levels
**
NPRO
NPRE
NEUR
MCH2-10
YK26
Nhlh2
WT
KO
PND3
PND7
PND21
NLGN3
Nlgn3

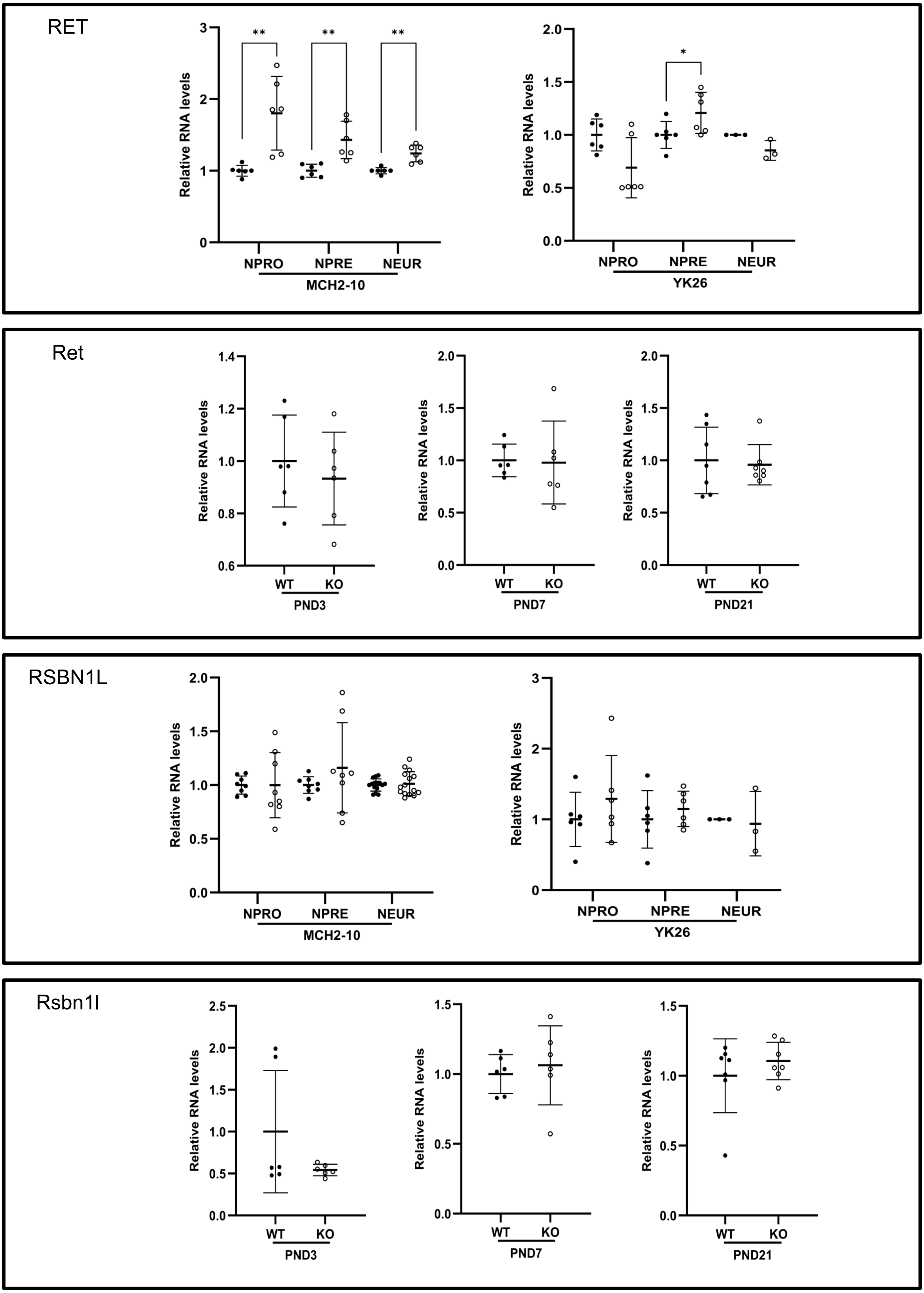

RET
Relative RNA levels
**
*
NPRO
NPRE
NEUR
MCH2-10
YK26
Ret
WT
KO
PND3
PND7
PND21
RSBN1L
Rsbn1l

SNX33

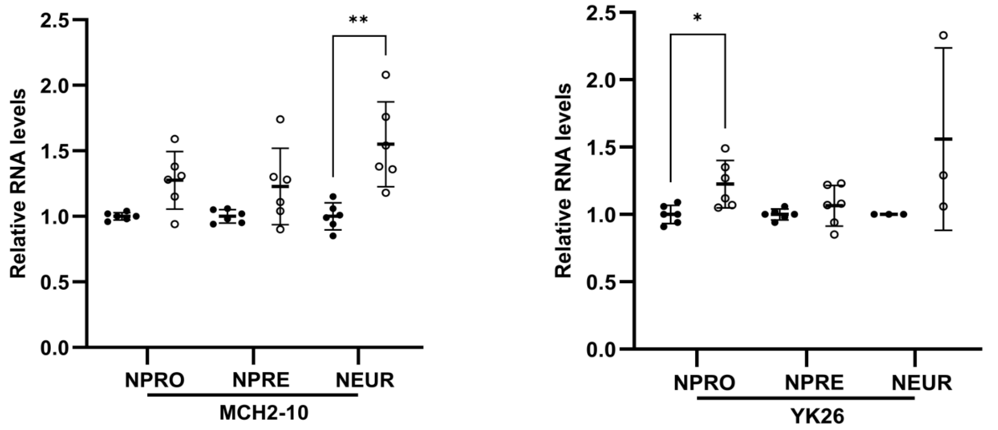


Snx33

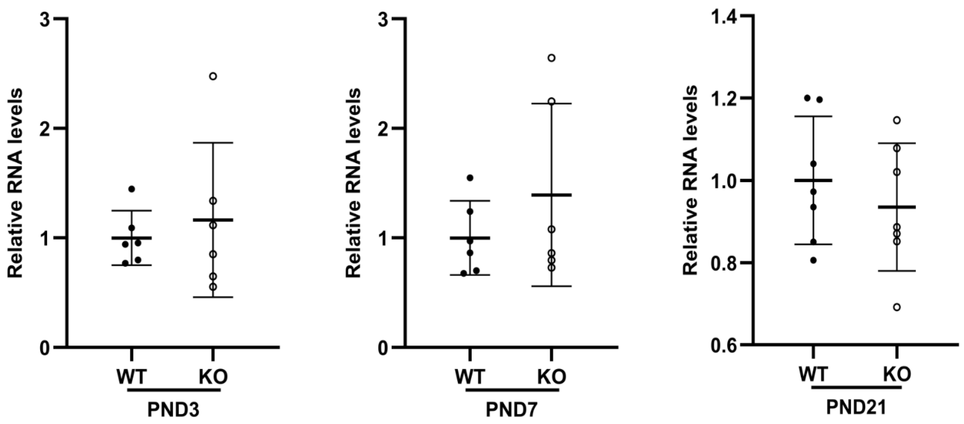


SYK

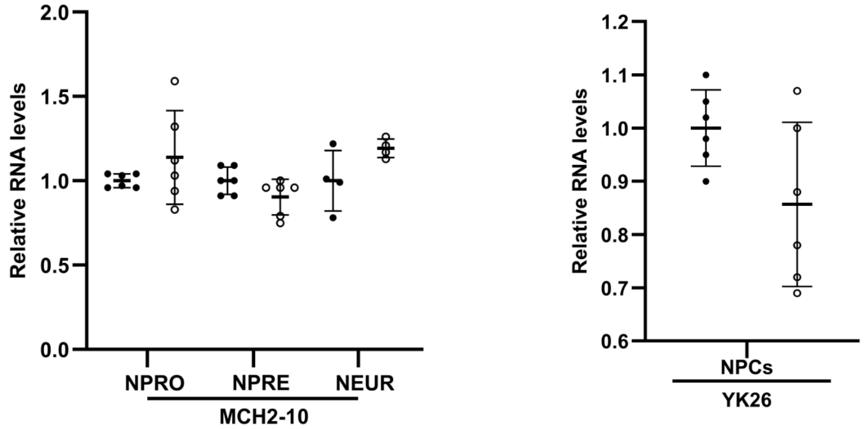


Syk

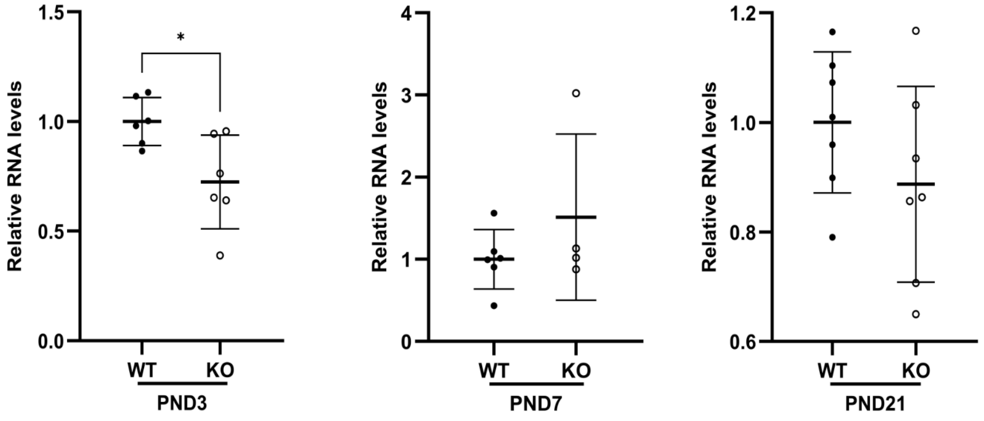

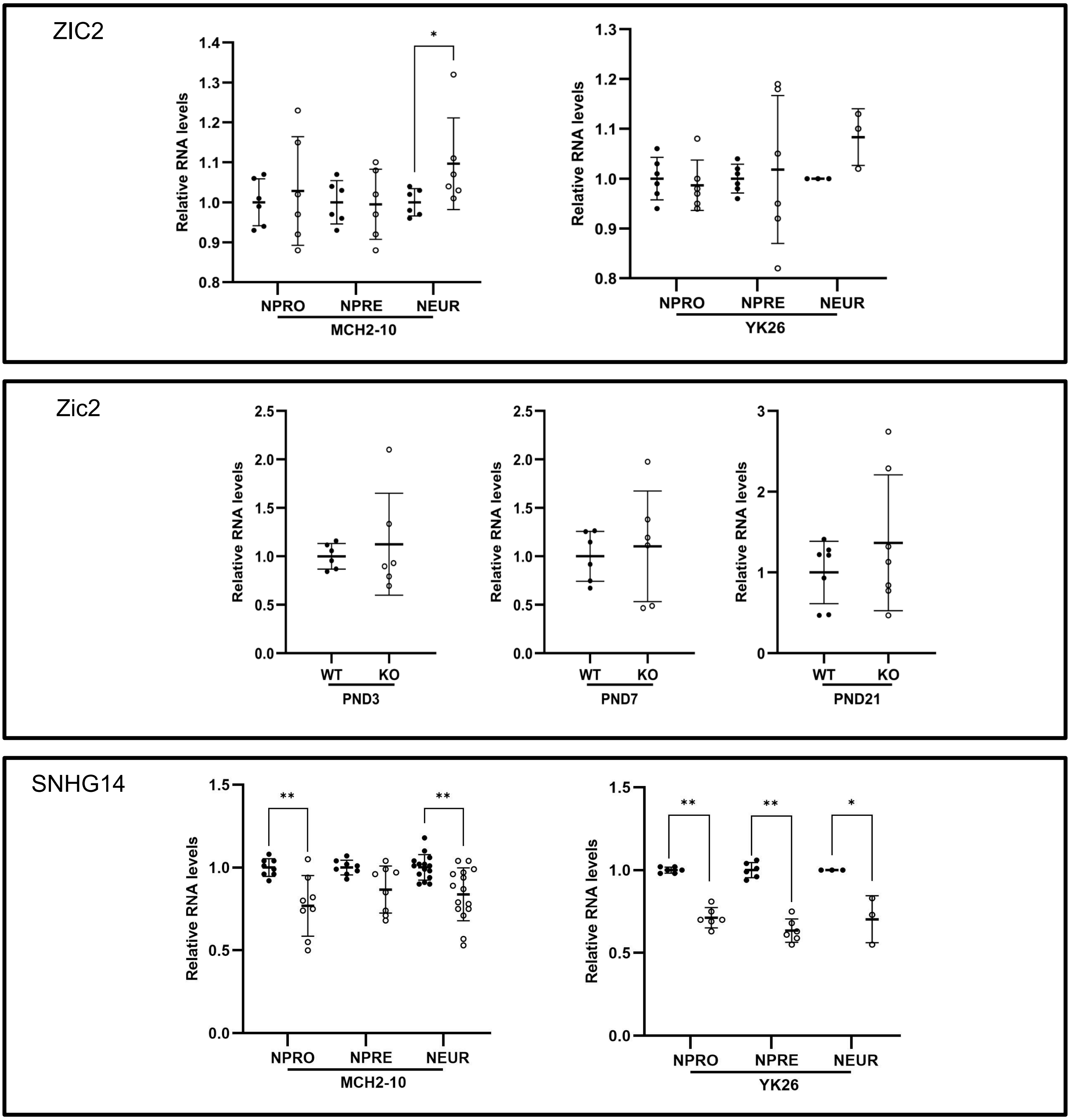
ZIC2
Relative RNA levels
NPRO
NPRE
NEUR
MCH2-10
YK26
Zic2
WT
KO
PND3
PND7
PND21
SNHG14

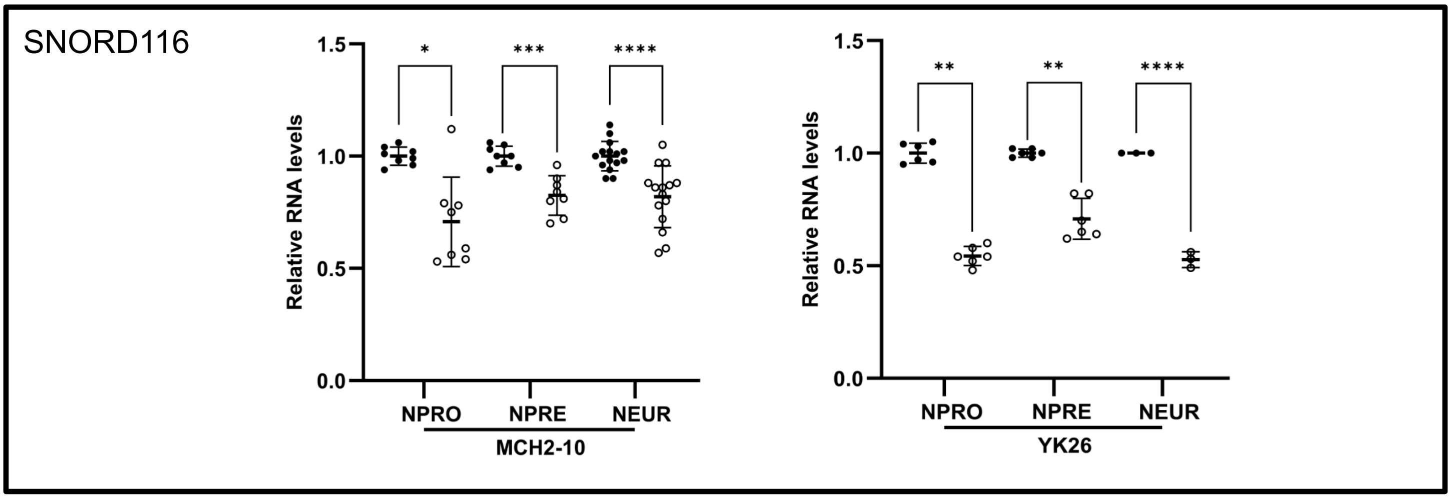
SNORD116
1.5
1.0
0.5
0.0
Relative RNA levels
*
***
****
NPRO
NPRE
NEUR
MCH2-10
**
**
****
NPRO
NPRE
NEUR
YK26

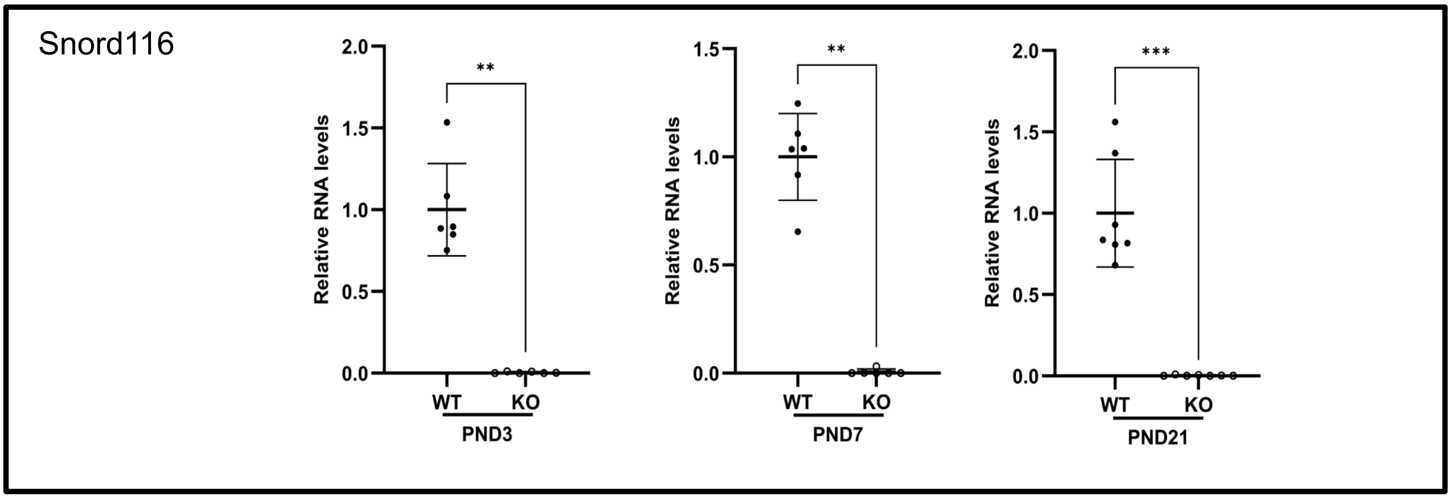
Snord116
2.0
1.5
1.0
0.5
0.0
Relative RNA levels
**
WT
KO
PND3
**
WT
KO
PND7
***
WT
KO
PND21

Supplementary Figure 3

A

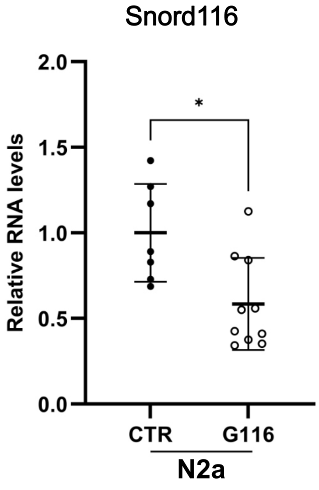


B

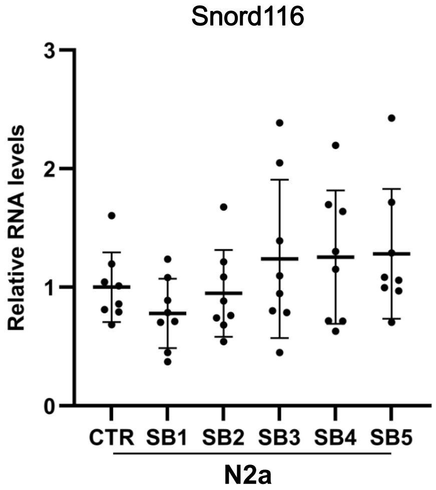

Supplementary Figure 4

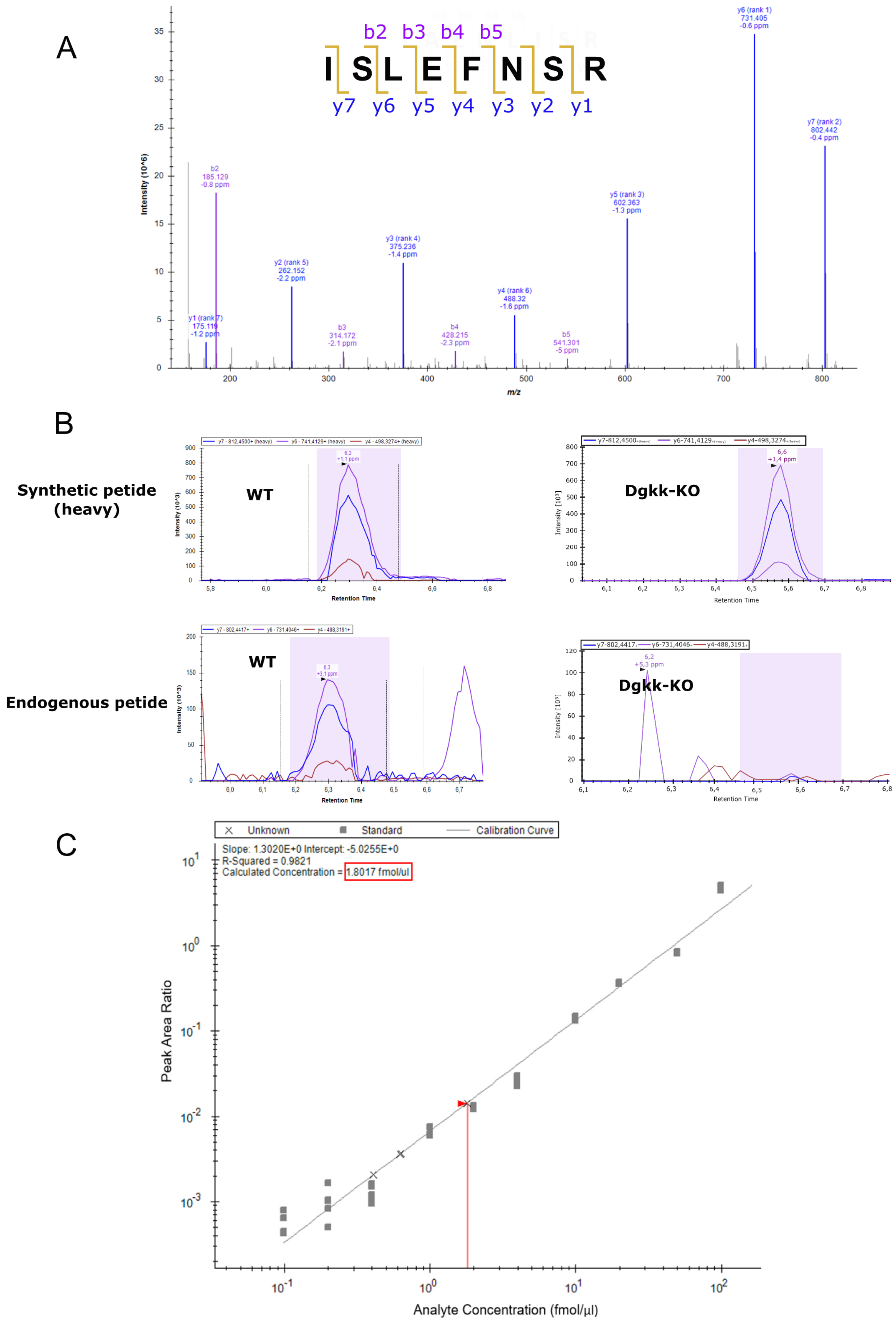

Supplementary Figure 5

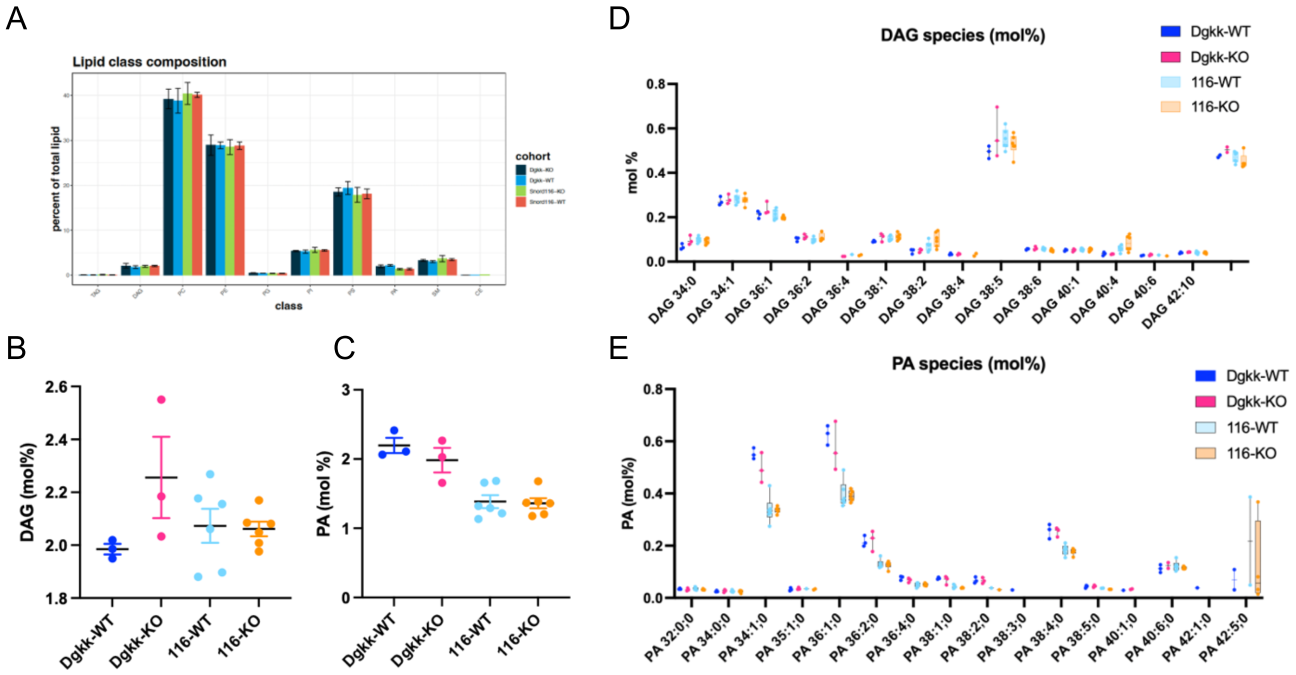

Supplementary Figure 6

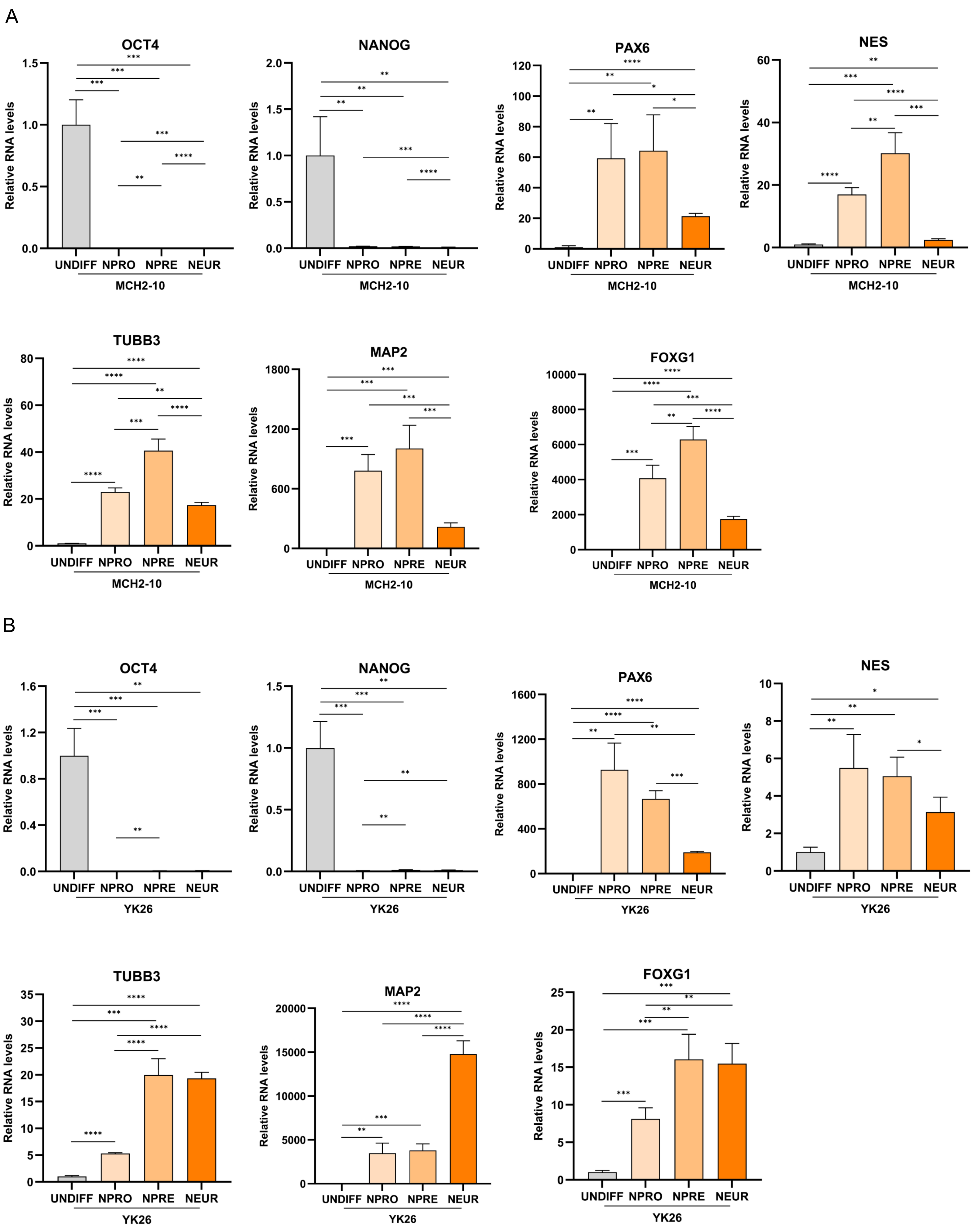
